# Analytic combinatorics for bioinformatics I: seeding methods

**DOI:** 10.1101/205427

**Authors:** Guillaume J. Filion

## Abstract

Seeding heuristics are the most widely used strategies to speed up sequence alignment in bioinformatics. Such strategies are most successful if they are calibrated, so that the speed-versus-accuracy trade-off can be properly tuned. In the widely used case of read mapping, it has been so far impossible to predict the success rate of competing seeding strategies for lack of a theoretical framework. Here I present an approach to estimate such quantities based on the theory of analytic combinatorics. In a nutshell, the strategy is to specify a combinatorial construction of reads where the seeding heuristic fails, translate this specification into a generating function using formal rules, and finally extract the probabilities of interest from the singularities of the generating function. I use this approach to construct simple estimators of the success rate of the seeding heuristic under different types of sequencing errors. I also show how the analytic combinatorics strategy can be used to compute the associated type I and type II error rates (mapping the read to the wrong location, or being unable to map the read). Finally, I show how analytic combinatorics can be used to estimate average quantities such as the expected number of errors in reads where the seeding heuristic fails. Overall, this work introduces a theoretical and practical framework to find the success rate of seeding heuristics and related problems in bioinformatics.

## 1 Introduction

High throughput sequencing is changing the face of biology [7,15]. More data is of course better, but recently, technical improvements have started to outpace the progress of algorithms [14]. When the problems are too large, one has to replace exact algorithms by heuristics that are much faster, but do not guarantee to return the right result. Good heuristics are all about understanding the input data. With the right data model, we can calculate the risk of not returning the right answer and adjust the algorithm to achieve more precision or more speed. When the data is poorly understood, heuristics may be slow or inefficient for unknown reasons.

A particular area of bioinformatics where heuristics have been in use for a long time is the field of sequence alignment [12]. Computing the best alignment between two sequences is carried out by dynamic programming in time *O*(*mn*), where *m* and *n* are the sequence lengths [2]. When at least one of the sequences is long (*e.g*. a genome), this is prohibitive and heuristics are required.

The most studied heuristics for sequence alignment are called seeding methods [16]. In a nutshell, the idea is to search short regions of the two sequences that are very similar and use them as candidates to anchor the dynamic programming alignment, which is performed only locally *i.e*. between subsequences of the input. These short regions of high similarity are called “seeds”. The benefit of the approach is that seeds can be found in short time. The risk is that they may not exist.

This strategy was most famously implemented in BLAST for the purpose of finding local homology between proteins [1]. By working out an approximate distribution of the identity score for the seeds [8,9], the authors were able to calibrate the BLAST heuristic very accurately in order to gain speed. The algorithm always performs the minimum amount of work for a desired confidence level.

Seeding methods are also heavily used in the mapping problem (figure 1), where the original sequence of a read must be found in a reference genome. The dicovery of indexing methods based on the Burrows-Wheeler transform [3] was instrumental to develop short read mappers such as BWA and Bowtie [10,11]. With such indexes, one can know the number of occurrences of a substring in a genome in time independent of the genome size. This yields a powerful seeding strategy whereby all the substrings of the read are queried in the genome. The main limitation is that the seeds must have exactly the same sequence in the read as in the genome.

The heuristic should be calibrated from the probability that a seed of given length can be found in the read, but this problem has not been fully solved. The answer depends on the types and frequencies of errors, which are often context-dependent. Overall, the lack of theory to model seeding probabilities hinders progress on this line of research.

Here I address this problem for arbitrary error models using the powerful theory of analytic combinatorics. This field of research was most significantly developed by Andrew Odlyzko, Robert Sedgewick and Philippe Flajolet [4-6]. The theory is now mature and used to tackle many problems outside the analysis of algorithms. However, it has not yet been fully realized how useful it can be for bioinformatics.

This document is predominantly written for bioinformaticians and people with a working knowledge of sequencing technologies and their applications. Accordingly, the focus will be on explaining the mathematical concepts, rather than the technological aspects. Also, my goal here is not to push the boundaries of analytic combinatorics, but to explain how its simplest concepts are useful to solve common problems in bioinformatics. I have opted for simplicity, to the detriment of generality and rigor. The results presented here are only a basic introduction to analytic combinatorics; the field is currently much more advanced and I refer the interested to the original literature [6].

**Figure 1:**
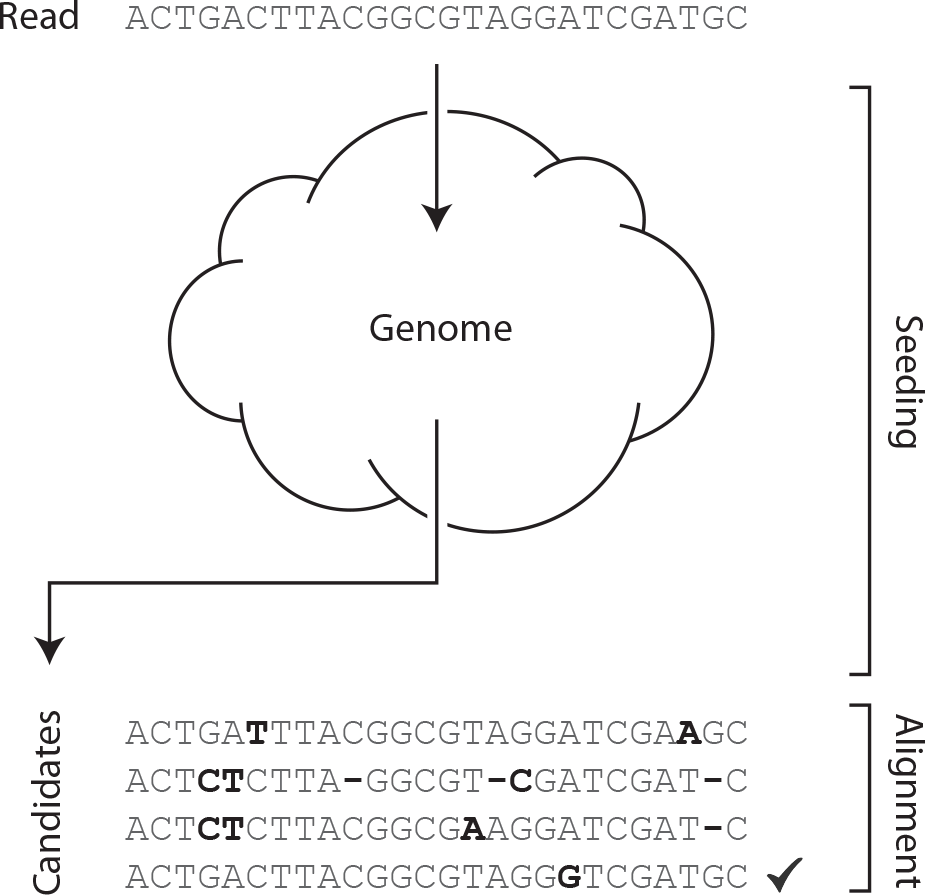
Seeding heuristic in the mapping problem. The sequencing read belongs to an unknown portion of the genome; the task is to find its original sequence (and usually its location). Because of possible read errors, the read may not be identical to the original sequence. Most algorithms feature a seeding stage and an alignment stage. The purposes of seeding and alignment are to output a short list of candidate sequences, and to determine which is the best, respectively. Seeding is a heuristic because it does not guanrantee that the overall best sequence is in the list of candidates.

## 2 Elements of analytic combinatorics

In order to develop a theoretical framework for the seeding problem, we first need to familiarize ourselves with the basic concepts of analytic combinatorics. This section is a general introduction to the concepts and tools, together with some more specialized material that will prove useful to tackle the problem at hand.

### 2.1 Weighted generating functions

The central object of analytic combinatorics is the *generating function*. Here we will develop a theory based on the slightly more general concept of *weighted* generating function.

#### Definition 1.

Let 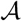 be a set of combinatorial objects characterized by a size and a weight. The **weighted generating function** of 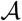 is defined as

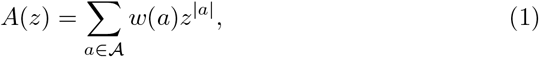

where |*a*| and *w(a*) denote the **size** and **weight** of the object a, respectively. Expression (1) also defines a sequence (*a_k_*)*_k≥0_* such that

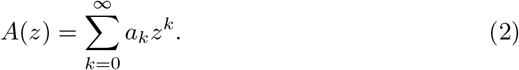

By definition 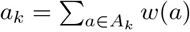, where *A_k_* is the class of objects of size k in 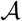. The number *a_k_* is called, the **total weight of objects of size** *k*.

#### Remark 1.

Expression (2) shows that the terms *a_k_* are the coefficients of the Taylor series expansion of the function *A(z*).

#### Remark 2.

If the weight of every object 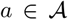 is 1, then *A(z*) is called a simple generating function, and in expression (2) 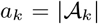 is the number of objects of size *k* in 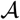.

#### Remark 3.

Typical combinatorial objects include binary trees, permutations, derangements, multisets etc., but here we will focus exclusively on sequences of symbols from finite alphabets. In what follows, the reader can think of “objects” as “finite sequences of symbols”.

Expressions (1) and (2) are of course equivalent. Depending on the context, we will use one or the other.

#### Example 1.

Let 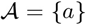 and 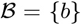 be alphabets with a single letter (of size 1). Assume *w(a*) = *p* and *w(b*) = *q*. The weighted generating functions of 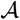 and 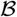 are then *A(z*) = *pz* and *B(z*) = *qz*, respectively.

The motivation for definition 1 is that operations on combinatorial objects translate into operations on their weighted generating functions. With the methods detailed below, we will obtain the weighted generating functions of elaborate combinatorial objects from simple ones. Using equation (2), we will extract the weight or probability of objects of size *k* from such weighted generating functions.

This approach is counter-intuitive at first, but propositions 4 and 5 below will show how we can compute probabilities that are inacessible to more frontal approaches.

Let us start with the most straightforward ways to obtain new weighted generating functions from those already available: additions and multiplications. If *A*(*z*) and *B*(*z*) are the weighted generating functions of two mutually exclusive sets 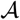 and 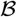, the weighted generating function of 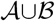 is *A*(*z*)+*B*(*z*), as appears immediately from expression (1).

#### Example 2.

With the definitions of example 1, the weighted generating function of the alphabet 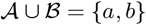 is *pz* + *qz* = *A*(*z*) + *B*(*z*).

Size and weight can be defined for pairs of objects in 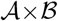 as |(*a*, *b*) | = |*a*| + |*b*| and *w*(*a*, *b*) = *w*(*a*)*w*(*b*). In other words the sizes are added and the weights are multiplied. With this convention, the weighted generating function of the Cartesian product 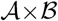 is *A*(*z*)*B*(*z*). This simply follows from expression (1) and

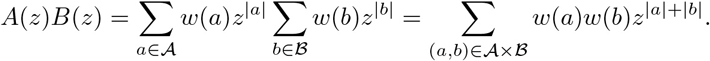

#### Example 3.

With the definitions of example 1, 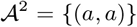 contains a single pair of letters, with size 2 and weight *p*^2^. Its weighted generating function is *p*^2^*z*^2^ = *A*(*z*)^2^.

#### Example 4.

Still with the definitions of example 1, 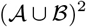 contains the four pairs of letters (*a, a*), (*a,b*), (*b,a*) and (*b,b*). They have size 2 and respective weight *p*^2^, *pq, qp*, and *q*^2^, so the weighted generating function of 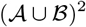 is (*p*^2^ + 2*pq* + *q*^2^)*z*^2^ = (*A*(*z*)+ *B*(*z*))^2^.

We can further extend the definition of size and weight to any finite Cartesian product in the same way. The sizes are always added and the weights are always multiplied. The generating function of a cartesian product then comes as the product of their generating functions. This allows us to construct finite sequences of objects.

#### Example 5.

With the definitions of example 1, 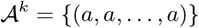 contains a single *k*-tuple of letters, with size *k* and weight *p^k^*, so its weighted generating function is *p*^*k*^*z*^*k*^. Since the sets 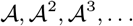 are mutually exclusive, the weighted generating function of their union is

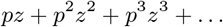

This infinite Taylor series can be expressed as a simple function. For any *k* ≥ 1, (1 – *pz*)(*pz* + (*pz*)^2^ + … + (*pz*)^*k*^) = *pz* – (*pz*)^*k*+1^. If |*z*| < 1/*p*, the term (*pz*)^*k*+1^ vanishes as *k* increases. So the weighted generating function of 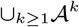 is defined for |*z*| < 1/*p* and is equal to

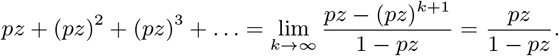

Example 5 is fundamental. It can be generalized to any set 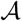 of combinatorial objects.

#### Proposition 1.

Let 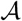 be a set with weighted generating function *A*(*z*). The weighted generating function of 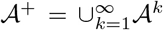, called the set of **non-empty sequences** of elements of 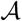 is defined for |*A*(*z*)| < 1 and is equal to

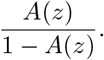

Proof. For k ≥ 1, the weighted generating function of 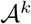 is A(*z*)^*k*^. Since the sets 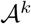 are mutually exclusive, the weighted generating function of their union is *A*(*z*) + *A*(*z*)^2^ + *A*(*z*)^3^ + … = *A*(*z*)/(1 – *A*(*z*)), provided |*A*(*z*)| < 1.

We also introduce the **empty object** *ε*, which has size 0 and weight 1. Its weighted generating function is thus 1. By convention, we define the zeroth power of a combinatorial set 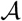 as 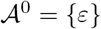. With this definition, we can state a variant of the previous proposition.

#### Proposition 2.

Let 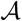 be a set with weighted generating function *A*(*z*). The weighted generating function of 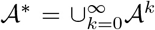, called the set of sequences of elements of 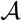 is defined for |*A*(*z*)| < 1 and is equal to

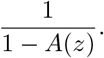

*Proof*. 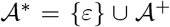. By proposition 1, the weighted generating function is 1 + *A*(*z*)/(1 – *A*(*z*)) = 1/(1 – *A*(*z*)), provided |*A*(*z*)| < 1.

#### Remark 4.

These expressions are not defined for *A*(*z*) = 1, i.e. when 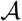 contains only the empty object. In other words, one cannot construct sequences of empty ojects. More generally, we will always assume that the sets used to construct sequences do not contain the empty object ε. Otherwise, ε is present in each 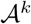 and they are not mutually exclusive.

#### Remark 5.

If |*A*(*z*) > 1, the function *A*(*z*)/(1 – *A*(*z*)) is well defined but the series *A*(*z*) + *A*(*z*)^2^ + … does not converge. This is not a problem, as relation (2) does not need to hold for every z. However, in what follows we will only consider the values of z for which the relation holds. In other words, we will always assume that |*z*| is smaller than the radius of convergence of the weighted generating function of interest.

#### Remark 6.

It is important to insist that the expression “sequences of” refers to the set 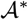 and not to the set 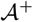, i.e. the empty object ε is a sequence of anything. For clarity, we will systematically use the expression “non-empty sequences of” when ε is excluded.

#### Example 6.

Following the definitions of example 1, 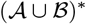 is the set of sequences of *a*’s and/or *b*’s. By proposition 2 and example 2, the weighted generating function of this set is

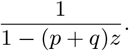

The function 1/(1 – (*p* + *q*)*z*) is just a compact representation of the infinite series 1, (*p* + *q*), (*p* + *q*)^2^, (*p* + *q*)^3^,… It thus carries all the information about the total weights of the sequences of any size. If *p* = *q* = 1, *i.e*. if we count the total amount of sequences with simple generating functions (see remark 2), we obtain 1/(1 – 2*z*) = 1 + 2*z* + 4*z*^2^ + 8*z*^3^ + 16*z*^4^ + … meaning that there are 2^*k*^ sequences of size *k*. If *p* and *q* are the respective probabilities of ocurrence of *a* and *b*, with *p* + *q* ≤ 1, then the equality 1/(1 – (*p* + *q*)*z*) = 1 + (*p* + *q*)*z* + (*p* + *q*)^2^*z*^2^ + (*p* + *q*)^3^*z*^3^ + … means that the probability that a sequence of size *k* contains only *a*’s and/or *b*’s is (*p* + *q*)^*k*^.

These simple examples are better solved by intuition. However, we will soon see that analytic combinatorics allows us to solve a wider range of problems.

### 2.2 Sequences and transfer matrices

In many combinatorial applications, one needs to count the sequences where a pattern does not occur, or where some symbols may not follow each other. A convenient way to find the weighted generating functions of such sequences is to encode this information in so-called “transfer matrices”. We will illustrate the process with an example from biology.

In many animal genomes, DNA methylation can only occur on the dinucleotide **`CG`**. Because of this property, it is interesting to count the sequences that have no **`CG`**. Sequences without **`CG`** can be thought of as walks on a directed graph with restricted transitions. Nucleotides are represtend as nodes, and edges indicate that the nucleotide at the tip can follow the nucleotide at the base. Sequences without **`CG`** correspond to walks on a complete graph where the edge from **`C`** to **`G`** is removed (see figure 2).

The same graph can also be represted as an adjacency matrix *M*, where a 1 at position (*i*, *j*) indicates that there is an edge from node *i* to node *j*, and a 0 indicates that there is no edge. For instance, the adjacency matrix of the graph of figure 2 is

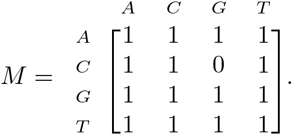

**Figure 2:**
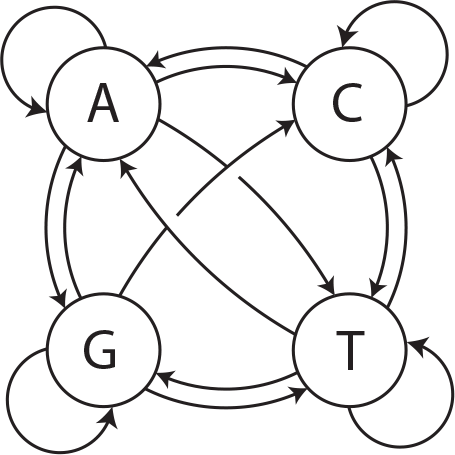
Graph representation of sequences of nucleotides without CG. A sequence without **`CG`** is equivalent to a walk on this graph. All the transitions are allowed, except **`C`** to **`G`**.

The main advantage of the adjacency matrix representation is that the powers of *M* give an analytical way to count the walks on the graph. The entry of *M*^*n*^ at position (*i,j*) is the number of walks of *n* steps that start with node *i* and end with node *j*. So we can count sequences without **`CG`** by computing the successive powers of the adjacency matrix *M* above.

The same idea can be used to find the weighted generating function of sequences of combinatorial objects. We specify the internal structure of sequences by a “transfer graph”, where the edges represent classes of combinatorial objects.

#### Definition 2.

**A transfer graph** is a directed graph whose edges are labelled by weighted generating functions(themselves representing classes of objects). In addition, a transfer graph must contain a head vertex with only outgoing edges, and a tail vertex with only incoming edges. The graph can be represented as a matrix called a **transfer matrix**.

Following the edges of a transfer graph from the head vertex to the tail vertex describes a sequence of combinatorial objects. The associated weighted generating function is the product of the functions labelling the edges (thus an absent edge is associated to the function 0).

Transfer graphs have the general structure below. The head vertex is represented as an open circle and the tail vertex as a closed circle. The edge labelled *ψ*(*z*), from the head vertex to the tail vertex represents additional sequences that we may add to fit a particular definition. Typically, *ψ*(*z*) = 1, as the only sequence that needs to be added explicitly is the empty object *ε*.

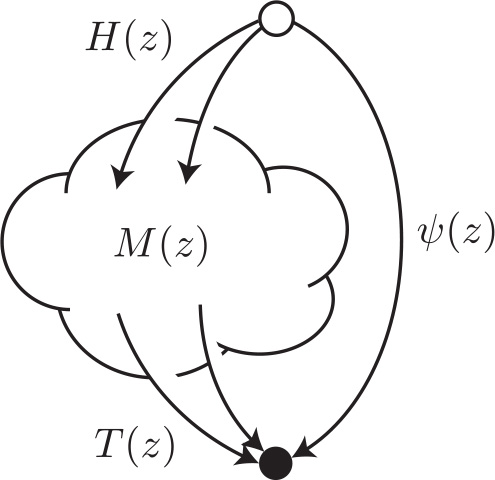

The “body” of the graph, represented as a cloud, contains the remaining m vertices. *M*(*z*) is the *m* × *m* matrix representation of the corresponding subgraph. *H*(*z*) is a vector of *m* weighted generating functions associated with the *m* edges from the head vertex to the body of the graph, and *T*(*z*) is a vector of *m* weighted generating functions associated with the edges from the body of the graph to the tail vertex. *H*(*z*) and *T*(*z*) are called the “head” and “tail” vectors, respectively.

#### Remark 7.

By convention, rows correspond to outgoing edges and columns to incoming edges. The head vertex always corresponds to the first row/column of the tansfer matrix and the tail vertex to the last.

#### Remark 8.

Transfer graphs are reminiscent of deterministic finite automata (DFA). The initial and accept states of a DFA correspond to the head and tail vertices of a transfer graph. The alphabet of a DFA corresponds to the combinatorial objects that can be concatenated (or to their weighted generating functions) and the transition function corresponds to the transfer matrix. We will not make further use of this analogy in this document.

#### Definition 3.

Concatenating the objects labelling the edges of a transfer graph in all possible ways for all possible paths from the head to the tail vertex describes the **sequences generated by the associated transfer matrix**.

#### Example 7.

Let 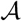 be a class of combinatorial objects with weighted generating function *A*(*z*). Sequences of objects from 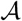 can be thought of as walks on the graph below.

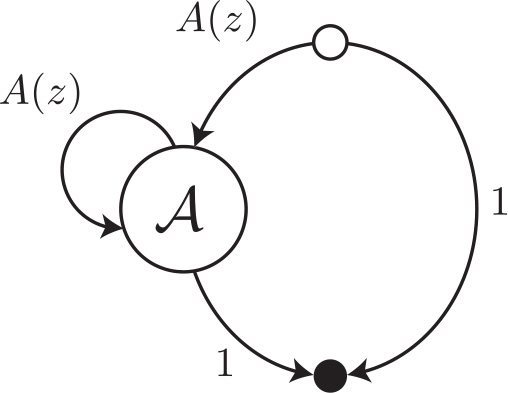

The edge from the vertex 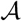 to the tail vertex marks the end of the sequence, by appending a final empty object *ε* with weight 1. The transfer matrix of this graph is the 3 × 3 matrix

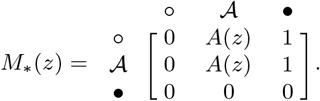

The head vector *H*(*z*) has dimension 1 and is equal to *A*(*z*). The tail vector *T*(*z*) also has dimension 1 and is equal to 1. *M*(*z*), the matrix of the body of the graph is the 1 × 1 matrix [*A*(*z*)].

*

#### Example 8.

Let 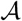 and 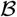 be mutually exclusive classes of combinatorial objects with respective weighted generating function *A*(*z*) and *B*(*z*). Sequences of objects from 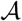 or 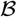 can be thought of as walks on the graph below.

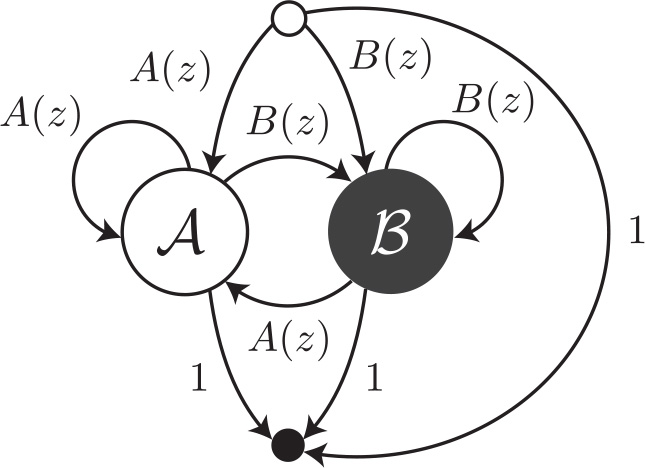

The transfer matrix of this graph is the 4 × 4 matrix

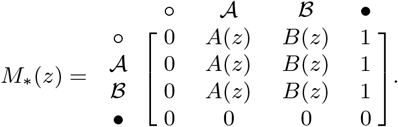

The head vector *H*(*z*) has dimension 2 and is equal to (*A*(*z*), *B*(*z*))^⊤^. The tail vector *T*(*z*) also has dimension 2 and is equal to (1,1)^⊤^. *M*(*z*), the matrix of the body of the graph is the 2 × 2 matrix

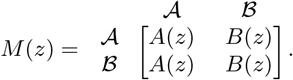

*

#### Example 9.

With the assumptions of example 8, sequences of objects from 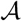 or 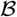 can also be thought of as walks on a the graph below.

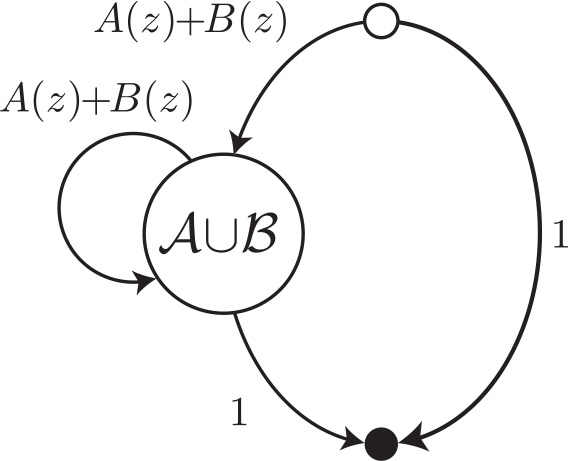

Indeed, because 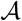 and 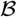 are mutually exclusive, we know that the weighted generating function of 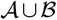 is *A*(*z*) + *B*(*z*). The transfer matrix of this graph is the 3 × 3 matrix

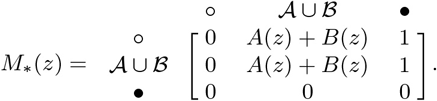

The head vector *H*(*z*) has dimension 1 and is equal to *A*(*z*) + *B*(*z*). The tail vector *T*(*z*) also has dimension 1 and is equal to 1. *M*(*z*), the matrix of the body of the graph is the 1 × 1 matrix *M*(*z*) = [*A*(*z*) + *B*(*z*)].

*

Examples 8 and 9 show that the same class of combinatorial sequences may be associated with multiple transfer graphs and transfer matrices. There are several ways to construct (and interpret) a weighted generating function.

#### Example 10.

Let 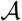 and 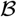 be mutually exclusive classes of combinatorial objects with respective weighted generating function *A*(*z*) and *B*(*z*). Sequences of objects from 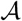 or 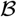 that end with 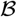 but without two consecutive objects from 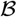 can be thought of as walks on the graph below.

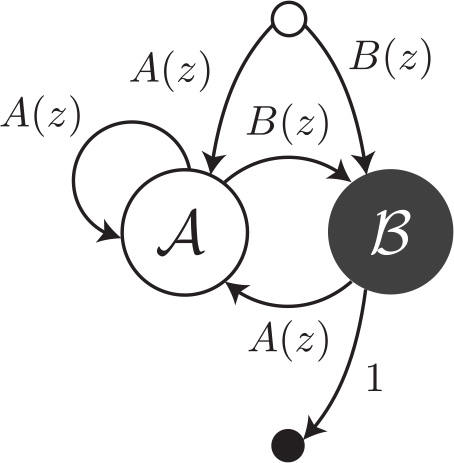

The edge with weighted generating function 1 from the head vertex to the tail vertex is no longer present because the empty object *ε* does not end with 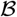. The transfer matrix of this graph is the 4 × 4 matrix

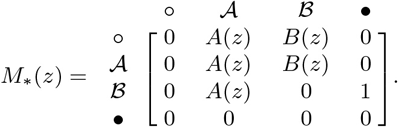

The head vector *H*(*z*) has dimension 2 and is equal to (*A*(*z*), *B*(*z*))^⊤^. The tail vector *T*(*z*) also has dimension 2 and is equal to (0,1)^⊤^. *M*(*z*), the matrix of the body of the graph is the 2 × 2 matrix

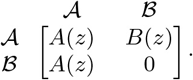

*

Let us now return to sequences of nucleotides without **`CG`**. To specify a transfer matrix, let us assume that **`G`**’s and **`C`**’s occur with frequency *p*/2 and that **`A`**’s and **`T`**’s occur with frequency *q*/2, where *q* = 1 – *p*. The weighted generating functions of the single nucleotides are thus *C*(*z*) = *G*(*z*) = *pz*/2 and *A*(*z*) = *T*(*z*) = *qz*/2. With these definitions, the transfer matrix of the graph shown in figure 2 is

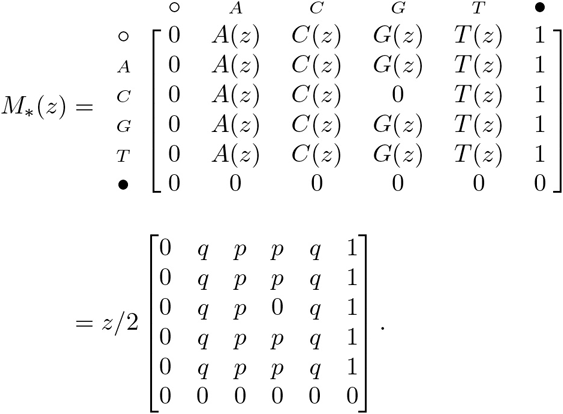

We still need to find the weighted generating function of sequences without **`CG`**. Proposition 3 below shows how this is done.

#### Lemma 1.

Given a transfer graph and its transfer matrix *M_∗_*(*z*), the weighted generating function of paths of *n* segments from vertex *i* to vertex *j* is the entry of *M_∗_*(*z*)^*n*^ at position (*i,j*).

*Proof*. Proceed by induction. For *n* =1, this is the definition of the transfer matrix. Assume that the property holds for some *n* ≥ 1. A path of *n* + 1 segments from vertex *i* to vertex *j* is a path of *n* segments from vertex *i* to some vertex *k*, followed by a single segment from vertex *k* to vertex *j*. Using the induction hypothesis and taking the union on all vertices 1 ≤ *k* ≤ *m* + 2 (recall that the transfer graph contains a head vertex, a tail vertex and *m* additional vertices), the weighted generating function of such paths is

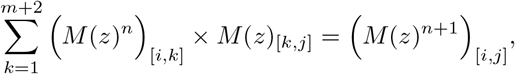

which concludes the proof by induction.

#### Proposition 3.

The weighted generating function of the sequences generated by a transfer matrix *M*_∗_(*z*) is the top right entry of the matrix (*I* – *M*_∗_(*z*))^−1^, assuming that all the eigenvalues of *M*(*z*) have modulus less than 1.

*Proof*. Top right entries correspond to paths from the head to the tail vertex (see remark 7). From lemma 1, the weighted generating function is the top right entry of the matrix

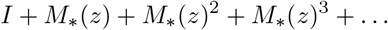

Observe that (*I*−*M*_∗_(*z*))·(*I*+*M*_∗_(*z*)+*M*_∗_(*z*)^2^ + …+*M*_∗_(*z*)^*n*^) = *I*−*M*_∗_(*z*)^*n*+1^. If all the eigenvalues of *M*_∗_(*z*) have *modulus* less than 1, the right hand side converges to the identity matrix *I* as *n* goes to infinity. This translates to

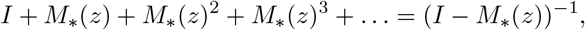

which concludes the proof.

#### Example 11.

Let us continue example 7 where we found that the tranfer matrix of sequences of objects from 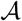 is

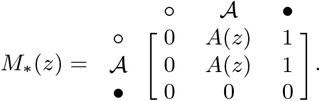

The weighted generating function of such sequences is the top right entry of the matrix

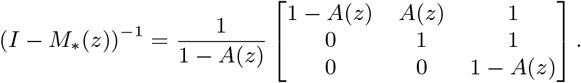

In other words, the weighted generating function of sequences of objects from 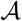 is 1/(1 – *A*(*z*)), consistently with proposition 2.

#### Example 12.

Let us continue examples 8 and 9 where we found two transfer matrices for sequences of objects from 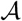 or from 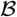. The first transfer matrix is

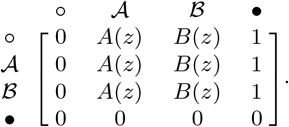

Applying proposition 3, we see that the weighted generating function is the top right entry of the matrix

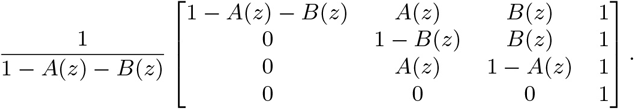

The second transfer matrix is

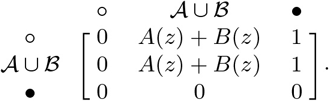

Applying proposition 3 again, we see that the weighted generating function is also the top right entry of the matrix

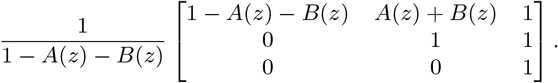

Both terms are equal to 1/(1−*A*(*z*)−*B*(*z*)). This illustrates that there are different ways to construct a sequence and its weighted generating function. This also illustrates that some constructions are simpler than others.

We now introduce a simplified way to find weighted generating functions from transfer matrices.

#### Proposition 4.

Given a transfer matrix

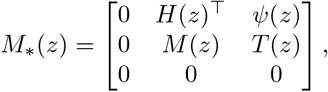

where *M*(*z*) is a *m* × *m* matrix, *H*(*z*) and *T*(*z*) are vectors of dimension *m* and *ψ*(*z*) has dimension 1, the weighted generating function of the sequences generated by *M*_∗_(*z*) is

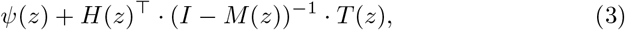

assuming that all the eigenvalues of *M*(*z*) have modulus less than 1.

*Proof*. We need to compute the top-right entry of the matrix (*I* – *M*_∗_(*z*))^−1^. Using the matrix inversion formula with the cofactor matrix, this term is equal to (−1)^*m*+2^*C*/ det(*I* – *M*_∗_(*z*)), where *C* is the determinant

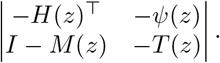

Developing det(*I* – *M*_∗_(*z*)) along the first column and then along the last row, we obtain det(*I* – *M*_∗_(*z*)) = (−1)^*m*^det(*I* – *M*(*z*)). Developping *C* along the first row and then along the last column, we obtain

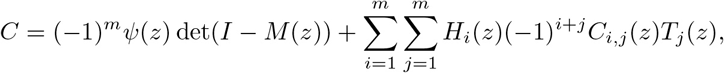

where *C*_*i,j*_ is the cofactor of *I* – *M*(*z*) at position (*i,j*). Using once more the matrix inversion formula, we obtain

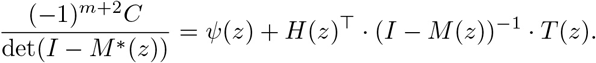

#### Example 13.

Let us continue example 10 where we found that the transfer matrix of sequences of objects from 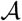 or from 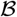 that end with an object from 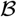 but do not contain two objects from 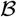 in a row is

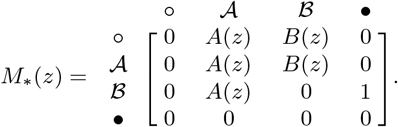

We have seen that *H*(*z*) = (*A*(*z*), *B*(*z*))^⊤^, *T*(*z*) = (0,1)^⊤^, *ψ*(*z*) = 0 and

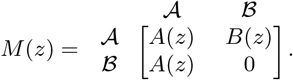

Using proposition 4, the weighted generating function of the sequences generated by *M*_∗_(*z*) is

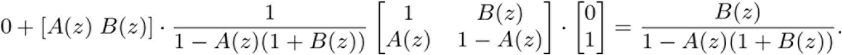

We will sometimes represent transfer graphs as their bodies (associated with the matrix *M*(*z*), not with *M*_∗_(*z*)), and specify separately *ψ*(*z*), *H*(*z*) and *T*(*z*). This will simplify the graphical representations, and using proposition 4 instead of proposition 3 will simplify the computations.

Now returning to sequences of nucleotides without **`CG`**, we have *ψ*(*z*) = 1, *H*(*z*) = (*A*(*z*), *C*(*z*), *G*(*z*), *T*(*z*))^⊤^ = (*qz*/2, *pz*/2, *pz*/2, *qz*/2), *T*(*z*) = (1,1,1,1)^⊤^, and

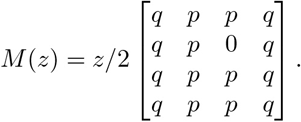

The inverse of *I* – *M*(*z*) is

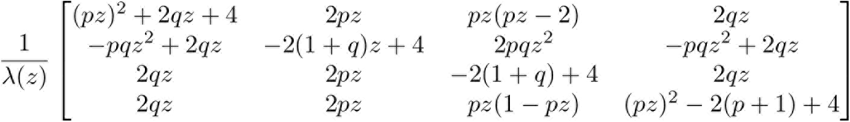

where λ(*z*) = 4 – 4*z* + (*pz*)^2^ is the determinant of *I* – *M*(*z*). Now applying proposition 4, the weighted generating function of sequences of nucleotides without **`CG`** comes out as

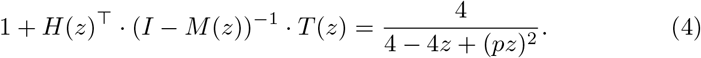

Getting to this weighted generating function was not straightforward, but the expression is relatively simple. This is commonly the case when taking the analytic combinatorics route: approaches that first look contrived and counterintuitive eventually lead to simple results.

In this section we have seen that transfer matrices allow us to find the weighted generating functions of potentially intricate sequences of combinatorial objects. In the next section we will see how we can compute the probabilities of occurrence of such sequences from their weighted generating function.

### 2.3 Asymptotic estimates

We know from expression (2) that we can recover the total weight (*i.e*. the probability) of objects of size *k* from the Taylor expansion of their weighted generating function. We will often need to extract *a*_*k*_ in expressions of the form 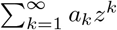 so we introduce the following symbol.

#### Definition 4.

Let *W*(*z*) be a weighted generating function. The expression

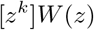

is called the **coefficient of** z^k^ **in** W and is equal to a_k_, assuming that 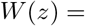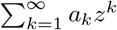 in some neighborhood of *z* = 0.

The coefficient of *z*^*k*^ in a weighted generating function is usually a quantity of interest. For instance, in expression (4) it is the probability that a nucleotide sequence of size *k* does not contain any **`CG`**.

One of the main motivations for the analytic combinatorics approach is that we can efficiently approximate the coefficients of weighted generating functions. Proposition 5 below is a simplified version of the founding theorem of the field [4].

#### Proposition 5.

If a function *W*(*z*) is the ratio of two polynomials *P*(*z*)/*Q*(*z*), and *Q* has only simple roots, then

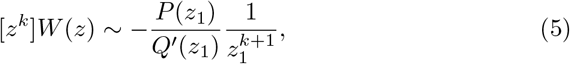

where *z*_1_ is the root of *Q* with smallest modulus.

Expression (5) is an asymptotic approximation: it becomes more accurate as *k* increases, *i.e*. as we consider objects of increasing size. This is the most fundamental property of the general analytic combinatorics strategy.

The roots of *Q* are called “singularities” of the function *W*. They are values where the function is not defined. The roots of *Q* with multiplicity 1 (*i.e*. the values for which *Q* vanishes but its derivative does not) are also referred to as “simple poles” of *W*.

Proposition 5 says that the asymptotic growth of the coefficients of the series expansion of *W*(*z*) is dictated by the singularity of smallest *modulus*, also known as the “dominant singularity”. We first state a lemma that will be useful to demonstrate proposition 5.

#### Lemma 2.

*If* |*z*| < *a, then*

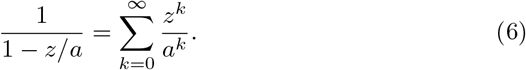

*Proof*. Proceed as in example 5, replacing *p* by 1/*a*.

We now prove proposition 5.

*Proof*. Let *z*_1_, *z*_2_,…, *z*_*n*_ be the complex roots of *Q*, sorted by increasing order of *modulus*. Since *Q* has only simple roots, there exists constants *β*_1_,…, *β*_*n*_ such that the partial fraction decomposition of the rational function *P*(*z*)/*Q*(*z*) can be written as

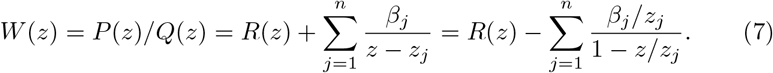

In the expression above, *R* is a polynomial that is nonzero if and ony if the degree of *P* is higher than the degree of *Q*. Either way, the coefficient of *z*^*k*^ in *R*(*z*) is 0 for *k* higher than the degree of *R*, so the coefficients of *R* do not contribute to [*z*^*k*^]*W*(*z*) for large *k*. We can thus assume *R*(*z*) = 0 without loss of generatlity. From lemma 2 we can expand each of the *n* terms of the sum as

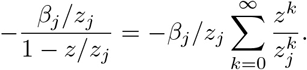

Substituting the expression above in (7), we obtain

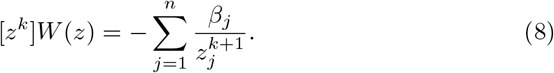

Since *z*_1_ is the root with smallest *modulus*, the sum above is dominated by the term 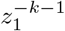 as *k* increases, so the coefficient of *z*^*k*^ in *W*(*z*) is asymptotically equivalent to

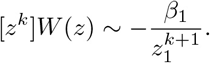

To find the value of *β*_1_, we factorize *Q*(*z*) as (*z* – *z*_1_)*Q*_1_(*z*), which is possible because *z*_1_ is a root of *Q*, and we write

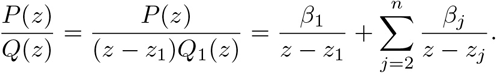

Multiplying through by (*z* – *z*_1_) and setting *z* = *z*_1_, we obtain the expression *β*_1_ = *P*(*z*_1_)/*Q*_1_ (*z*_1_). Differentiating the expression *Q*(*z*) = (*z* – *z*_1_)*Q*_1_(*z*) shows that *Q′*(*z*_1_) = *Q*_1_(*z*_1_), and thus that *β*_1_ = *P*(*z*_1_)/*Q′*(*z*_1_), which concludes the proof.

#### Remark 9.

Expression (8) is not an approximation, it is the exact value of the coefficient of *z*^*k*^ in *W*(*z*) = *P*(*z*)/*Q*(*z*). By keeping more than one term in the sum, we can obtain more accurate estimates, and by keeping all the terms we obtain the exact value.

#### Remark 10.

The asymptotic estimate (5) converges exponentially fast to the coefficient as *k* increases. To see this, divide the exact expression (8) by its leading term 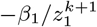 and obtain

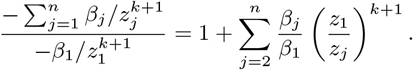

Since |*z*_1_| < |*z*_*j*_| for 2 ≤ *j* ≤ *n*, the error terms are *O*(|*z*_1_/*z*_2_|^*k*^) so they decrease exponentially fast as *k* increases.

#### Remark 11.

Proposition 5 does not hold if *z*_1_ is not a simple pole. The proof can be beneralized, but the resulting asymptotic formula is different. Proposition 10 in section 5.2 shows the coefficient asymptotics for poles of second order.

Recall from section 2.2 that the weighted generating function of nucleotide sequences without **`CG`** is

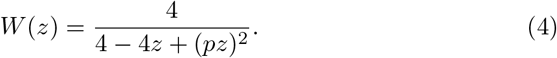

Here *Q*(*z*) =4 – 4*z* + (*pz*)^2^ has two distinct roots, 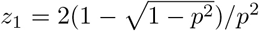 and 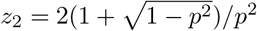. Since *Q*′(*z*) = 2*p*^2^*z* – 4, proposition 5 yields

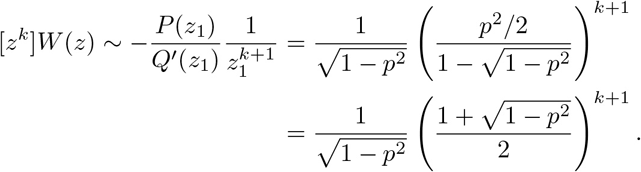

In this case we can also obtain the exact value by using the second singularity. The probability that a sequence of size *k* contains no **`CG`** is exactly equal to

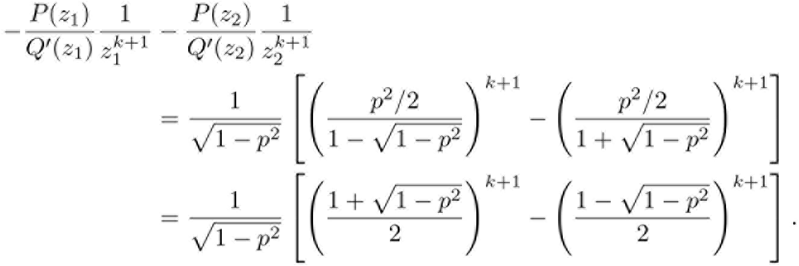

If all the nucleotides have the same frequency, then *ps* =1/2 and the probability is approximately equal to (0.9330127^*k*+1^ – 0.0669873^*k*+1^)/1.154701. From *k* = 5, the second term of the sum is over one million times smaller than the first term.

This example illustrates the elegance and the efficiency of the analytic combinatorics approach. It also shows how accurate the approximate solution can be.

In conclusion, we summarize the analytic combinatorics strategy as follows: (*i*) define simple objects associated with simple generating functions, (*ii*) combine these objects into more complex structures, (*iii*) translate those combinations into more complex generating functions, and (*iv*) extract coefficients using approximation theorems such as proposition 5.

## 3 Exact seeding

### 3.1 Reads, error symbols and error-free intervals

From the experimental point of view, a sequencing read is the result of an assay on some polymer of nucleic acid. The output of the assay is the decoded sequence of monomers that compose the molecule. Three types of sequencing errors can occur: *substitutions, deletions* and *insertions*. A substitution is a nucleotide that is different in the molecule and in the read, a deletion is a nucleotide that is present in the molecule but not in the read, and an insertion is a nucleotide that is absent in the molecule but present in the read.

For our purpose, the focus is not the nucleotide sequence *per se*, but whether the nucleotides are correct. Thus, we need only four symbols to describe a read: one for each type or error, plus one for correct nucleotides. In this view, a read is a finite sequence of letters from an alphabet of four symbols.

Errors in the read are initially unknown. In practice, the only way to detect them is to align the read with the reference sequence in order to highlight the differences. This operation may be difficult to perform, especially if the read has many errors. While exposing the theory, it will sometimes seem that the errors are known before aligning the read. Such cases will serve illustrative and didactic purposes, but they never occur in practice.

A read can be seen as a sequence of symbols. Figure 3 shows the typical structure of a read, together with the symbols that we will use for correct nucleotides, substitutions, deletions, and insertions.

**Figure 3:**
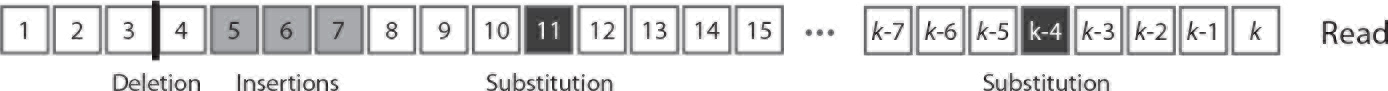
Read as sequence of symbols. Reads consist of correct nucleotides (white boxes), substitutions (black boxes), deletions (vertical bars) and insertions (grey boxes).

A read can be partitioned uniquely into maximal sequences of identical symbols called “intervals”. Intervals of correct nucleotides will have a particular importance, so we introduce the somewhat simpler term “error-free interval”.

#### Definition 5.

An **error-free interval** is a sequence of correct nucleotides that cannot be extended left or right.

Instead of sequences of symbols, reads can be seen as sequences of either error-free intervals or error symbols (see figure 4). We will use this construction throughout section 3. As developed below, this will allow us to control the size of the largest error-free interval.

These formal definitions will allow us to specify the weighted generating functions of the reads of interest and then approximate their probability of occurrence using the results of section 2.

**Figure 4:**
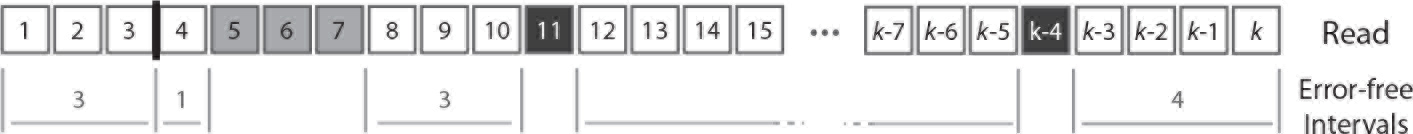
Read as sequence of error-free intervals or error symbols. Consecutive correct nucleotides can be lumped together in error-free intervals. The representation of a read as a sequence of either error-free inetervals or error symbols is unique. The error-free intervals are highlighted in this example, together with their size.

### 3.2 Exact seeds

One of the most common operations is to map the sequenced reads in a reference genome, *i.e*. find the identity of the fragment that was sequenced. The exact search problem can be solved efficiently, provided the right indexing data structures are available, but sequencing errors make the task more difficult.

In most mapping algorithms, the search starts by looking for perfect matches between a fragment of the read and a sequence from the genome. This match is called a “seed”, and it allows to quickly eliminate most of the search space in order to focus on a few promising candidate sequences.

This strategy greatly accelerates the mapping process, but it is not failsafe. Because of sequencing errors, it could be that no seed is found, or worse, that a match is found at the wrong location of the genome. Thus, error-free intervals hold the key to the problem. If the read contains a long enough error-free interval, it may be used as a seed. We will use a definition of “seed” that corresponds to this intuition.

#### Definition 6.

***An exact γ-seed** is an error-free interval of size at least γ*.

#### Remark 12.s

In what follows, we will often refer to an “exact γ-seed” as an “exact seed” or as a “seed” when the concrete value of *γ* is either clear from the context or irrelevant.

The main goal of section 3 is to construct estimators of the probability that a read contains an exact γ-seed, based on our knowledge of the typical sequencing errors. Our strategy is to construct the weighted generating functions of reads that *do not* contain any exact γ-seed. For this, we will decompose such reads as sequences of either error symbols or error-free intervals of size less than γ. We will obtain their weighted generating functions from proposition 4 and use proposition 5 to approximate their probability of occurrence.

We will find the weighted generating function of all the reads *R*(*z*), and then the weighted generating function of reads without exact γ-seed *S*_γ_(*z*). The probability that a read of size *k* has no exact γ-seed will then be equal to the total weight of reads of size *k* without seed, divided by the total weight of reads of size *k*, *i.e*.

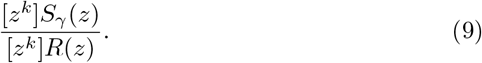

To introduce the concepts progressively, we will first describe simplified models where some types of errors are disallowed, and gradually increase the complexity towards a model with all error types.

### 3.3 Substitutions only

In the simplest model, we assume that errors can be only substitutions, and that they occur with the same probability *p* for every nucleotide. We will refer to this model as the “uniform substitutions error model”. Due to its simplicity, we will be able to go at greater depth and obtain more advanced results with this model (especially in section 4). Importantly, the model is not overly simple and it has some real practical applications. For instance, it describes reasonably well the error model of the Illumina platforms, where *p* is around 0.01 [13].

We will first use proposition 4 to obtain the weighted generating function *R*(*z*) of all reads under this error model, from which we will deduce the weighted generating function *S*_γ_(*z*) of reads without exact γ-seed. We will then use proposition 5 to derive an approximation formula for the probability that a read does not contain exact γ-seed.

Under the uniform substitutions model, reads are sequences of either substitutions or error-free intervals. They can be thought of as a walk on the graph shown in figure 5. The symbol Δ_0_ stands for error-free intervals and the symbol *S* stands for single substitutions. *F*(*z*) and *pz* indicate the weighted generating functions of error-free intervals and substitutions, respectively. The fact that an error-free interval cannot follow another error-free interval is a consequence of the definition: two consecutive intervals must be merged into a larger interval.

#### Remark 13.

In all the error models described in section 3, the error-free intervals cannot have size 0. The reason is that a read must correspond to exactly one path in the transfer graph and that sequences of combinatorial classes containing the empty object ε are not uniquely defined, (see remark 4).

A substitution is a single nucleotide and thus has size 1. Because substitutions have probability *p*, their weighted generating function is *pz*. Conversely, the weighted generating function of correct nucleotides is *qz*. Error-free intervals are non-empty sequences of correct nucleotides, so by proposition 2 their weighted generating function is

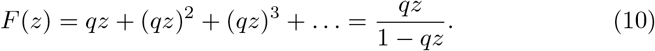

The transfer matrix associated with the body of the transfer graph shown in figure 5 is

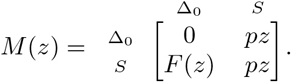

A read can start with an error-free interval or with a substitution, so the head vector *H*(*z*) is equal to (*F*(*z*), *pz*)^⊤^. Similarly, a read can end with an error free interval or with a substitution, so the tail vector *T*(*z*) is equal to (1,1)^⊤^. Here *ψ*(*z*) = 1, as we still need to include reads of size 0, *i.e*. the empty sequence *ε*. Applying proposition 4, the weighted generating function of reads is *R*(*z*) = *ψ*(*z*) + (*F*(*z*), *pz*)^⊤^ · (*I* – *M*(*z*))^−1^ · *T*(*z*), or

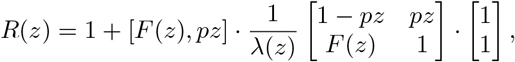

where λ(*z*) = 1 – *pz*( 1 + *F*(*z*)) is the determinant of *I* – *M*(*z*). Finishing the computations, we obtain

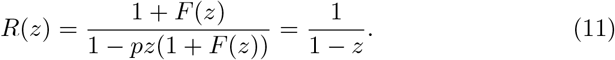

Since 1/(1 – *z*) = 1 + *z* + *z*^2^ + …, this means that for any *k* ≥ 0, the total weight of reads of size *k* is equal to 1. As a consequence, ratio (9) reduces to the numerator [*z*^*k*^]*S*_γ_(*z*).

To find the weighted generating function of reads without exact γ-seed, we need to limit error-free intervals to a maximum size of γ – 1. To do this, we can replace *F*(*z*) in (11) by its truncation *F*_γ_(*z*) = *qz* + (*qz*)^2^ + … + (*qz*)^γ−1^. We obtain

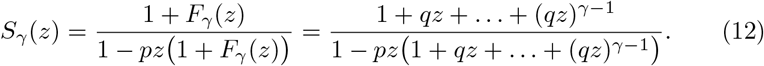

**Figure 5:**
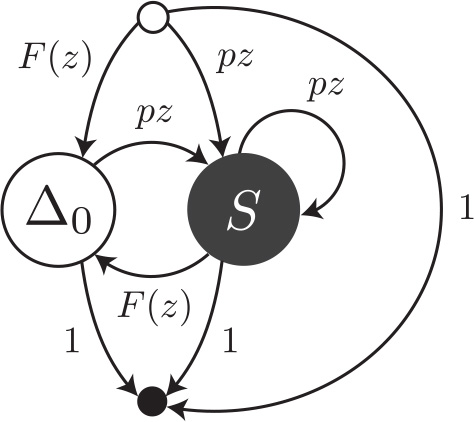
Transfer graph of reads with uniform substitutions. Reads are viewed as sequences of error-free intervals (symbol Δ_0_) or substitutions (symbol *S*). *F*(*z*) and *pz* are the weighted generating functions of error-free intervals and individual substitutions, respectively.

#### Remark 14.

Viewing reads as sequences of either error-free intervals or substitutions was important to obtain the weighted generating function of reads without seed: it allowed us to just replace *F*(*z*) by *F*_γ_ (*z*) in order to find the expression immediately.

To compute ratio (9) we now need to compute the coefficient of *z*^*k*^ in *S*_γ_(*z*). As explained in section 2.3, we extract an asymptotic estimate of it using proposition 5. For this, we need to find the singularities of *S*_γ_ (*z*).

It is interesting to look in detail into the problem of finding the singularities of *S*_γ_(*z*). For simplicity, we start with a concrete case. The left panel of figure 6 shows the values of the denominator of *S*_γ_(*z*) with *p* = 0.1 and 7 = 17. *S*_17_ has one real root greater than 1. The remaining singularities of *S*_17_ are complex and seem to be evenly spaced on a circle, as can be seen on the right panel of figure 6. This is only a visual impression. In fact the singularities do not lie on a perfect circle and their rotation angles are not exactly regular.

It is “fortunate” that the dominant root of *S*_17_ is a real number because we can use efficient numerical methods to approximate it (e.g. bisection or Newton-Raphson method). The following proposition shows that this is no accident: the dominant singularity of *S*_γ_ is always a real positive number greater than 1.

#### Proposition 6.

For 0 <*p*< 1, *q* =1 – *p* and γ ≥ 1, *S*_γ_ as expressed in (12) has exactly one positive real singularity. This is the dominant singularity and it is greater than 1.

*Proof*. Write *S*_γ_(*z*) = *P*(*z*)/*Q*(*z*) and search the roots of the denominator *Q*(*z*) = 1 – *pz*(1 + *qz* + … + (*qz*)^γ−1^). First, we show that *Q* has exactly one positive real root greater than 1. For real *z* > 0, *pz* and 1 + *qz* + … + (*qz*)^γ−1^ are strictly increasing, so *Q*(*z*) is strictly decreasing. Since *Q*(1) = *q*^*d*^ > 0 there is no root in the interval (0,1). Since lim_*z*_→_∞_ *Q*(*z*) = −∞, *Q* vanishes for a real number greater than 1.

Second, we show that this is the root with smallest *modulus*. Express the complex roots of *Q* as Re^*iθ*^, with *R* > 0 and 0 ≤ *θ* < 2π. They satisfy the equation

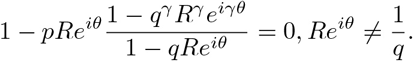

Multiplying through by 1 −*qRe*^iθ^, we obtain an equation of which we separate the real and the imaginary parts to obtain

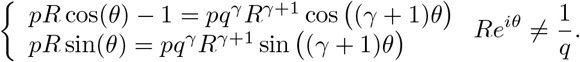

Squaring and summing, we obtain the following equation

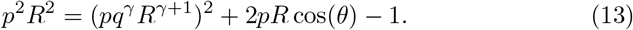

Considering this an equation of *R* > 0, the solution is minimal when 2*pR* cos(*θ*) is maximal, *i.e*. when *θ* = 0. In other words, if there is a positive real root, its *modulus* is the minimum among the roots. We have seen above that there exsists exactly one, so it is the dominant singularity of *S*_γ_.

We now have all the tools to approximate the coefficients of *S*_γ_ (*z*) using proposition 5 and obtain the approximate probability that a read contains no seed.

**Figure 6:**
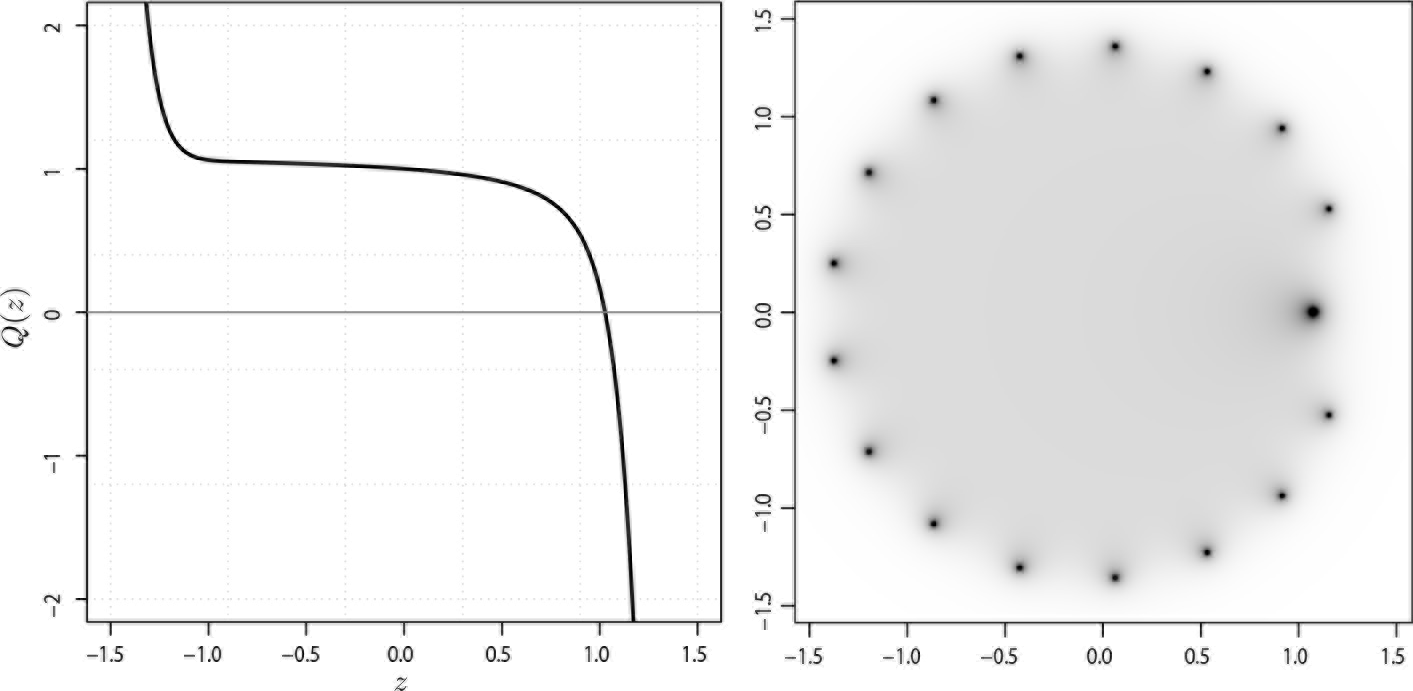
Singularities of *S*_17_. *Q*(*z*) denotes the denominator of *S*_17_(*z*) from expression (12) with *p* = 0.1 and γ = 17. *Left*: The values of *Q*(*z*) are represented as a bold line for *z* real. *Right*: The values of |1/*Q* (*z*) | are represented for *z* complex by the heat map on the complex plane (the numbers indicate the real and imaginary parts of *z*). Darker pixels correspond to higher values. Sixteen singularities of *S*_17_ lie close to a circle. The remaining seventeenth is the one that appears on the left panel. It is the dominant singularity because it is the closest to the origin.

#### Proposition 7.

The probability that a read of size *k* has no seed under the uniform substitutions model is asymptotically equivalent to

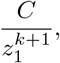

where z_1_ is the only real positive root of 1 – *pz*(1 + *qz* + … + (*qz*)^γ−1^), and where

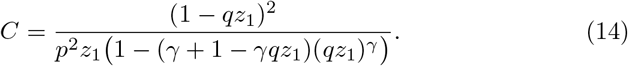

*Proof*. Apply propositions 5 to *S*_γ_(*z*) and then proposition 6, together with the fact that for any singularity *z* of *S*_γ_ we have 1 + *qz* + … + (*qz*)^γ−1^ = 1/*pz*.

We now illustrate proposition 7 with a concrete example explaining how the calculations are done in practice.

#### Example 14.

Let us approximate the probability that a read of size *k* = 100 has no seed for γ =17 and for a substitution rate *p* = 0.1. To find the dominant singularity of *S*_17_, we need to solve 1−0.1*z*× (1 + 0.9*z*+ … + (0.9*z*)^16^) = 0. We rewrite the equation as 1 – 0.1*z* × (1 – (0.9*z*)^17^)/(1 – 0.9*z*) = 0 and use bisection to solve it numerically, yielding *z*_1_ ≈ 1.0268856. Substituting this value in (14) yields *C* ≈ 1.396145, so the probability that a read contains no seed is approximately 1.396145/1.0268856^101^ ≈ 0.095763. For comparison, a 99% confidence interval obtained by performing 10 billion random simulations is 0.09575 – 0.09577. The computational cost of the analytic combinatorics approach is infinitesimal compared to the random simulations, and the precision is much higher for *k* = 100.

Overall, the analytic combinatorics estimates are close to the exact values. Figure 7 illustrates the precision of the estimates for different values of the error rate *p* and of the read size *k*.

### 3.4 Substitutions and deletions

To not jump too fast into the difficulties, we will now describe a semi-realistic model where errors can be deletions or substitutions, but not insertions. As in the case of uniform substitutions, we assume that every nucletoide call is false with a probability *p* and true with a probability 1 – *p* = *q*. Here, we also assume that the “space” between consecutive nucleotides can contain a deletion with probability *δ*.

A deletion may be adjacent to a substitution, or lie in between two correct nucleotides. In the first case, the deletion does not interrupt any error-free interval so it does not change the probability that the read contains a seed. For this reason, we ignore deletions next to substitutions. More precisely, we assume that they can occur, but whether they do has no importance for the problem.

**Figure 7:**
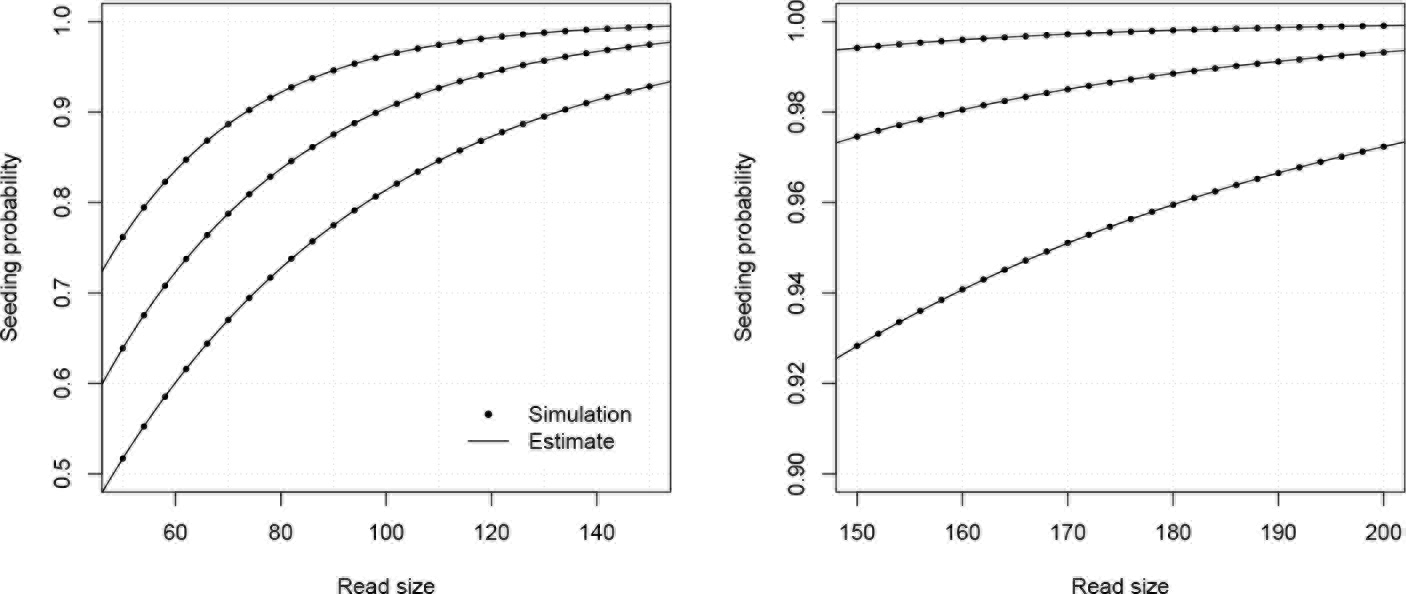
Example estimates for substitutions only. The analytic combinatorics estimates of proposition 7 are benchmarked against random simulations. Shown on both panels are the probabilities that a read of given size contains a seed, either estimated by 10,000,000 random simulations (dots), or by proposition 7 (lines). The curves are drawn for γ =17 and *p* = 0.08, *p* = 0.10 or *p* = 0.12 (from top to bottom).

Under this error model, a read can be thought of as a walk on the graph shown in figure 8. The graph is almost the same as the one shown figure 5; the only difference is the edge labelled *δF*(*z*) from Δ_0_ to Δ_0_. This edge represents the fact that under this error model, an error-free interval can follow another one if a deletion with weighted generating function *δ* is present in between (as illustrated for instance in figure 4).

The weighted generating function of error-free intervals *F*(*z*) has a different expression from that of section 3.3. When the size of an error-free interval is 1, the weighted generating function is just *qz*. For a size *k* > 1, there are *k* – 1 “spaces” between the nucleotides, so the weighted generating function is (1 – *δ*)^*k*−1^(*qz*)^*k*^. Summing for all the possible sizes, we obtain the weighted generating function of error-free intervals as

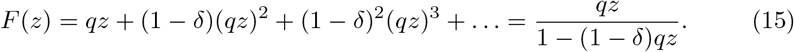

The transfer matrix associated with the body of the transfer graph shown in figure 8 is

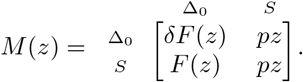

**Figure 8:**
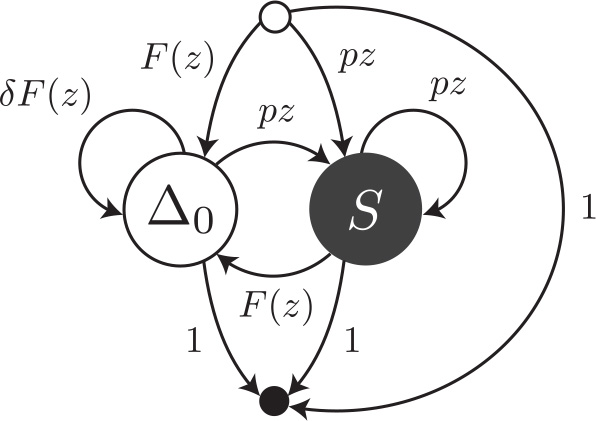
Transfer graph of reads with uniform substitutions and deletions. Reads are viewed as sequences of error-free intervals (symbol Δ_0_) or substitutions (symbol *S*). An error-free interval can follow another one if a deletion is present in between. *F*(*z*) and pz are the weighted generating functions of error-free intervals and individual substitutions, respectively. *δF*(*z*) is the weighted generating function of a deletion followed by an error-free interval.

The head and tail vectors are the same as in section 3.3, *i.e H*(*z*) = (*F*(*z*), *pz*)^⊤^, *T*(*z*) = (1,1)^⊤^ and *ψ*(*z*) = 1. Applying proposition 4 yields

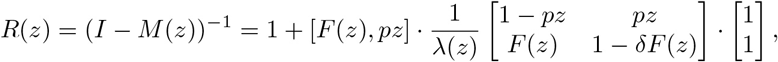

where λ(*z*) = 1 – *pz* – (*pz*(1 – *δ*) + *δ*)*F*(*z*) is the determinant of *I* – *M*(*z*). Finishing the computations we obtain

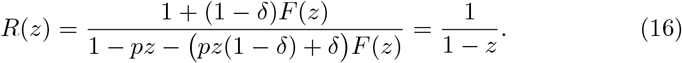

As in section 3.3 the result is 1/(1 – *z*) = 1 + *z* + *z*^2^ + …, which means that the total weight of reads of size *k* is equal to 1 for every *k* ≥ 0. This also means that ratio (9) reduces once again to its numerator [*z*^*k*^]*S*_γ_(*z*).

To find the weighted generating function of reads without exact γ-seed, we need to bound the size of error-free intervals to a maximum of γ – 1, *i.e*. to replace *F*(*z*) by its truncation *F*_γ_(*z*) = *qz* + (1 – *δ*)(*qz*)^2^ +… + (1 – *δ*)^γ−2^(*qz*)^γ−1^. With this, the weighted generating function of reads without seed is

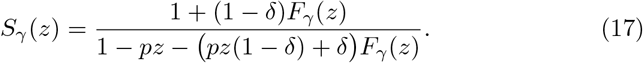

Applying proposition 5 to this expression, we obtain the following approximation formula.

#### Proposition 8.

The probability that a read of size k has no seed under the model of uniform substitutions and deletions is asymptotically equivalent to

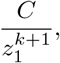

where z_1_ is the only real positive root of the denominator of *S*_γ_(*z*), *i.e.* the root of 1 – *pz* – (*pz*(1 – *δ*) + *S*)(*qz* + (1 – *δ*)(*qz*)^2^ + … + (1 – *δ*)^γ−2^(*qz*)^γ−1^1*, and*

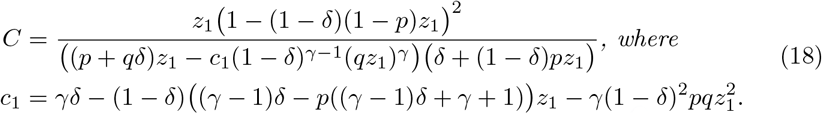

We illustrate proposition 8 with a concrete example explaining how the calculations are done in practice.

#### Example 15.

Let us approximate the probability that a read of size *k* = 100 has no seed for γ = 17, *p* = 0.05 and *δ* = 0.15. To find the dominant singularity of *S*_17_ we solve 1 – 0.05*z* – (0.0425*z* + 0.15)(0.95*z* + 0.85(0.95*z*)^2^ + … + 0.85^1B^(0.95*z*)^16^) = 0. We write it as 1 – 0.05*z*-(0.0425*z* + 0.15)(0.95*z*-0.85^16^(0.95*z*)^17^)/(1-0.8075*z*) =0 and use bisection, yielding *z*_1_ ≈ 1.006705. Now substituting the obtained value in (18) gives *C* ≈ 1.096177, so the probability that a read contains no seed is approximately 1.096177/1.006705^101^ ≈ 0.558141. For comparison, a 99% confidence interval obtained by performing 10 billion random simulations is 0.55813-0.55816.

Once again, the analytic combinatorics estimates are close to the exact values. Figure 9 illustrates the precision of the estimates for different values of the deletion rate *δ* and of the read size *k*.

**Figure 9:**
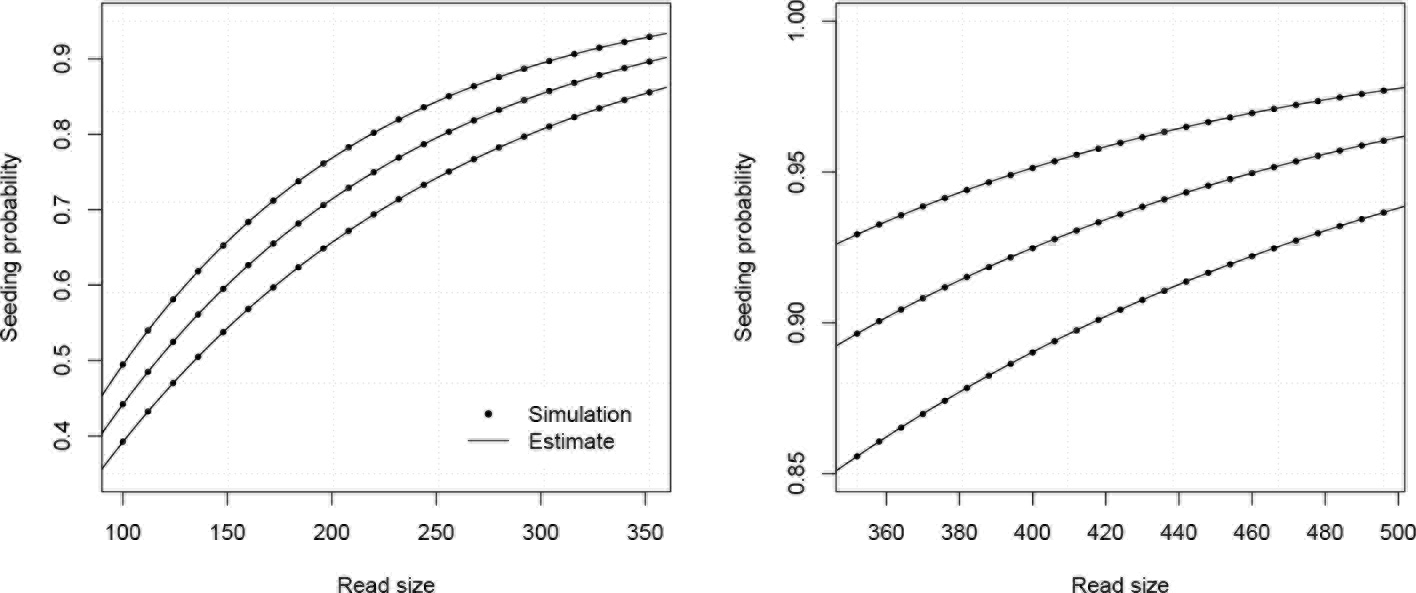
Example estimates for substitutions and deletions. The analytic combinatorics estimates of proposition 8 are benchmarked against random simulations. Shown on both panels are the probabilities that a read of given size contains a seed, either estimated by 10,000,000 random simulations (dots), or by proposition 8 (lines). The curves are drawn for γ = 17, *p* = 0.05 and *δ* = 0.14, *δ* = 0.15 or *δ* = 0.16 (from top to bottom).

### 3.5 Substitutions, deletions and insertions

Here we consider the model that we will refer to as “full error model” because all types of errors are allowed. Introducing insertions brings two additional difficulties. The first is that a substitution is indistinguishable from an insertion followed by a deletion (or a deletion followed by an insertion). By convention, we will count all these cases as substitutions. As a consequence, a deletion can never be found next to an insertion. The second difficulty is that insertions usually come in bursts. This is also the case of deletions, but we could neglect it because this does not affect the size of the interval (all deletions have size 0).

To model insertion bursts, we need to assign a probability *r* to the first insertion, and a probability *r̃* > *r* to all subsequent insertions of the burst. We will still denote the probability of a substitution *p* and that of a correct nucleotide *q*, but here *p* + *q* + *r* = 1. We will also assume that an insertion burst stops with probability 1 – *r̃* at each position of the burst.

Under this error model, reads can be thought of as a walk on the graph shown in figure 10. The symbols Δ_0_, *S* and *I* stand for error-free intervals, single substitutions and single insertions, respectively. The body of the graph is represented separately from the head and tail vertices to avoid overloading the figure.

The terms *F*(*z*), *pz* and *δF*(*z*) are the same as in section 3.4. The terms *rz* and *r̃z* are the weighted generating functions of the first inserted nucleotide and of all subsequent nucleotides of the insertion burst, respectively. The burst terminates with probability 1 – *r̃* and is followed by an error-free interval or by a substitution. The total weight of these two cases is *p* + *q* < 1, so we need to further scale the weighted generating functions by a factor *p* + *q* =1 – *r*.

The expression of the weighted generating function of error-free intervals *F*(*z*) is the same as in section 3.4, namely

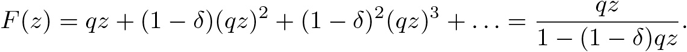

The transfer matrix associated with the body of the transfer graph shown in figure 10 is

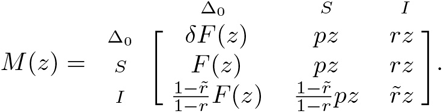

Here the head vector *H*(*z*) is equal to (*F*(*z*), *pz*, *rz*)^⊤^, the tail vector *T*(*z*) is equal to (1,1,1)^⊤^, and *ψ*(*z*) = 1. The expression of (*I* – *M*(*z*))^−1^ is omitted because it is cumbersome, but according to proposition 4 all we need is the value of *ψ*(*z*) + *H*(*z*)^⊤^ · (*I* – M(*z*))^−1^ · *T*(*z*), which is equal to

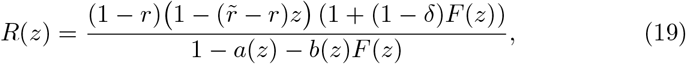

where *a*(*z*) and *b*(*z*) are second degree polynomials defined as

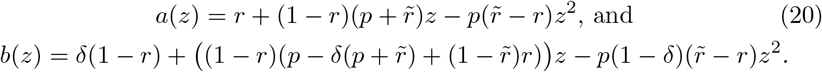

Substituting in (19) the expressions of *F*(*z*), *a*(*z*) and *b*(*z*), the terms cancel out and we find

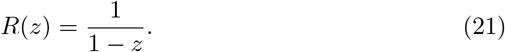

Again, we obtain the simple expression 1/(1 – *z*) = 1 + *z* + *z*^2^ + … where all the coefficients are equal to 1. This means that once again, ratio (9) reduces to the numerator [*z*_*k*_]*δ*_γ_(*z*) without exact γ-seed, we replace *F*(*z*) in expression (19) by its truncated version

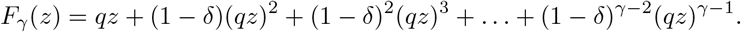

We obtain the following expression

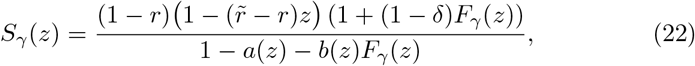

where *a*(*z*) and *b*(*z*) are defined as in (20).

**Figure 10:**
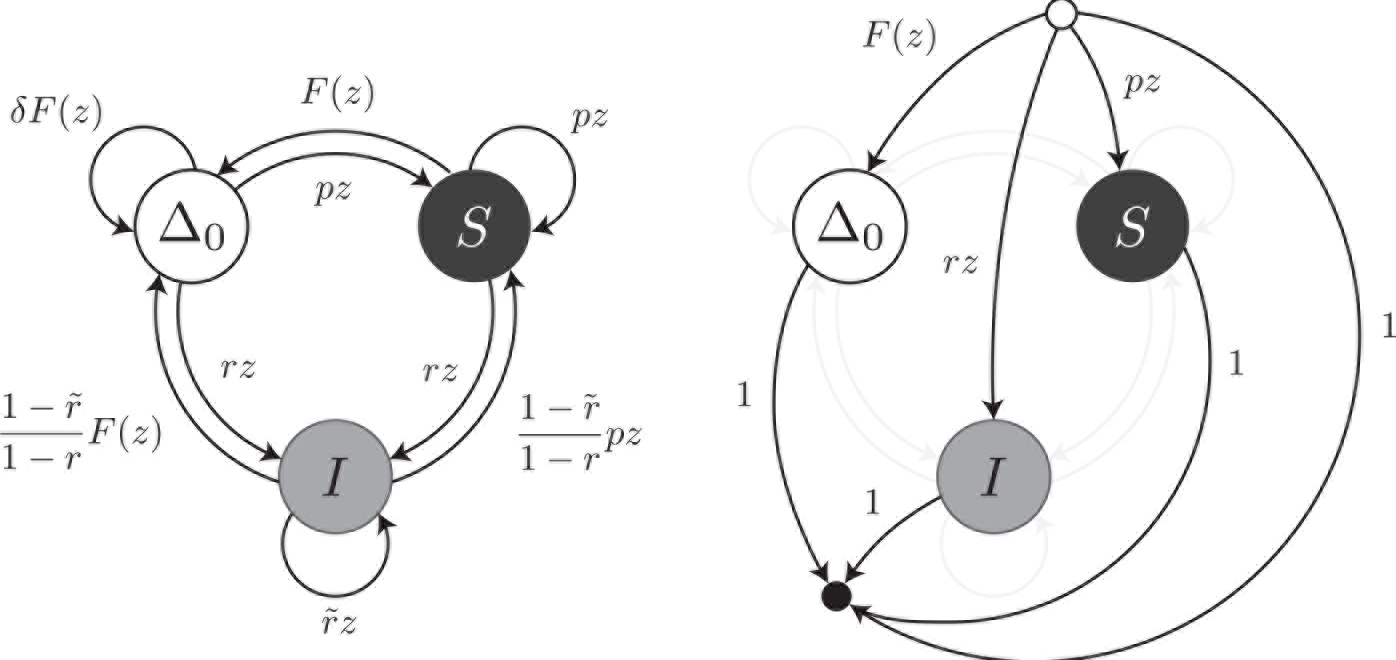
Transfer graph of reads with full error model. Reads are viewed as sequences of error-free intervals (symbol Δ_0_), substitutions (symbol *S*) or insertions (symbol *I*). To not overload the figure, the body of the transfer graph is shown on the left, and the head and tail edges are shown on the right. *F*(*z*) and *pz* are the weighted generating functions of error-free intervals and individual substitutions, respectively. *δF*(*z*) is the weighted generating function of a deletion followed by an error-free interval. *rz* and *r̃z* are the weighted generating functions of the first and all subsequent insertions of a burst, respectively. (1 – *r̃*)*F*(*z*)/(1 – *r*) is the weighted generating function of an error-free interval following an insertion and (1 – *r̃*)*pz*/(1 – *r*) is the weighted generating function of a substitutition following an insertion.

#### Remark 15.

Note that when *r* = *r̃* = 0, *a*(*z*) = *pz* and *b*(*z*) = *pz*( 1 – *δ*) + *δ*, so expression (22) becomes

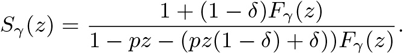

This is expression (17), i.e. the model described in section 3.4. When we also have *δ* = 0, this expression further simplifies to

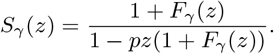

This is expression (12), i.e. the model described in section 3.3. In other words, the error models described previously are special cases of the full error model.

As in the previous sections, we can use proposition 5 to obtain asymptotic approximations for the probability that the reads contain no seed.

#### Proposition 9.

The probability that a read of size k has no seed under the full error model is asymptotically equivalent to

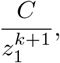

where *z*_1_ is the only real positive root of the polynomial 1 – *a*(*z*) – *b*(*z*)*F*_γ_(*z*) = 1 – *a*(*z*) – *b*(*z*)(*qz* + (1 – *δ*)(*qz*)^2^ + … + (1 – *δ*)^γ−2^(*qz*)^γ−1^), *and C* = *c*_1_/*c*_2_, with

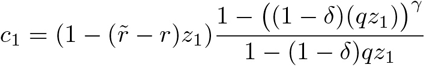

and

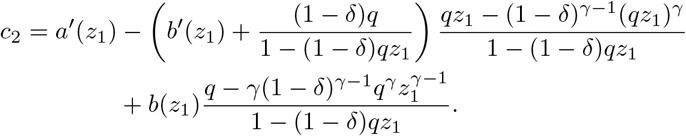

We illustrate proposition 9 with a concrete example explaining how the calculations are done in practice.

#### Example 16.

Let us approximate the probability that a read of size *k* = 100 has no seed for γ = 17, *p* = 0.05, *δ* = 0.15, *r* = 0.05 and *r̃* = 0.45. With these values, *a*(*z*) = 0.05 + 0.475*z*-0.02*z*^2^ and *b*(*z*) = 0.1425 + 0.00375*z*-0.017*z*^2^. We need to solve 0.95-0.475*z* + 0.02*z*^2^-(0.1425 + 0.00375*z*-0.017*z*^2^)(0.9*z* + 0.85(0.9*z*)^2^ + … + 0.85^15^(0.9*z*)^16^) = 0. We rewrite the equation as 0.95-0.475*z* + 0.02*z*^2^-(0.1425 + 0.00375*z*-0.017*z*^2^)(0.9*z*-0.85^15^(0.9*z*)^16^)/(1-0.765*z*) = 0 and use bisection to solve it numerically, yielding *z*_1_ ≈ 1.00295617. Using proposition 9, we obtain *C* ≈ 1.042504, so the probability that a read contains no seed is approximately 1.042504/1.00295617^101^ ≈ 0.773749. For comparison, a 99% confidence interval obtained by performing 10 billion random simulations is 0.77373 – 0.77376.

Once again, the analytic combinatorics estimates are close to the exact values. Figure 11 illustrates the precision of the estimates for different values of the insertion rate *r* and of the read size *k*.

**Figure 11:**
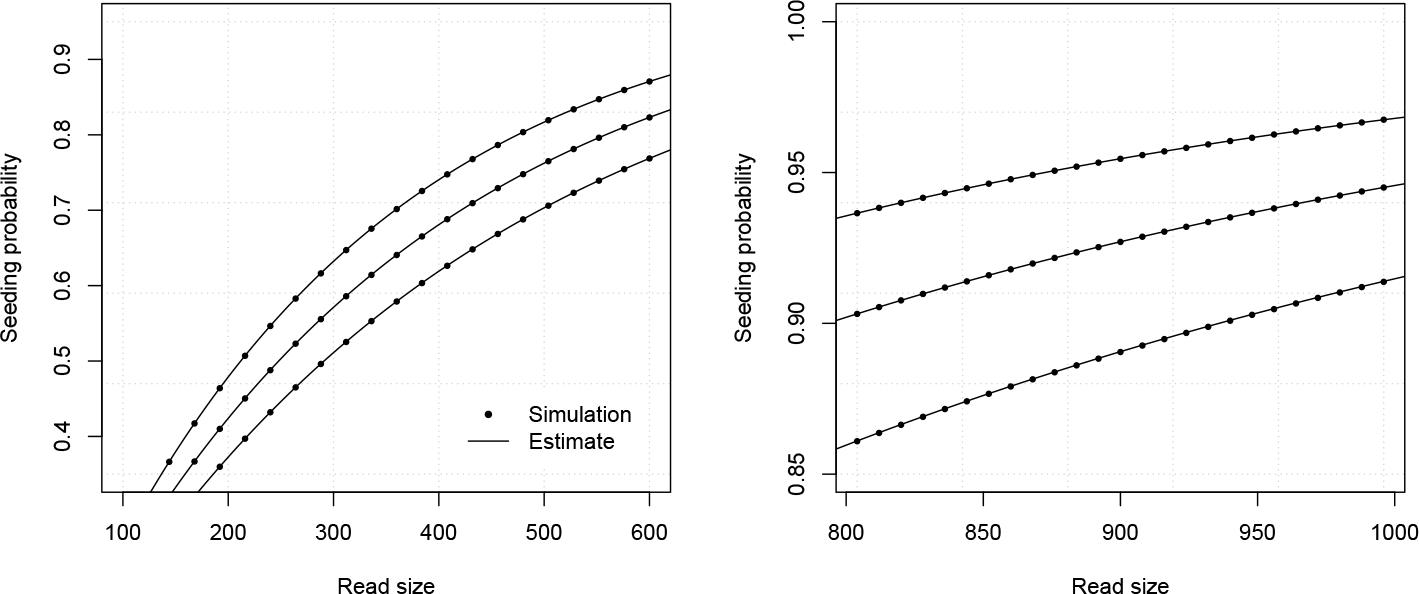
Example estimates for substitutions, deletions and insertions. The analytic combinatorics estimates of proposition 9 are benchmarked against random simulations. Shown on both panels are the probablities that a read of given size contains a seed, either estimated by 10,000,000 random simulations (dots), or by proposition 9 (lines). The curves are drawn for γ = 17, *p* = 0.05, *δ* = 0.15, *r* = 0.45 and *r̃* = 0.04, *r* = 0.05 or *r* = 0.06 (from top to bottom).

### 3.6 Empirical error models

In the theory developed so far, we introduced different kinds of errors because they have different probabilities and different sizes, but the nature of the error is irrelevant. To know whether the read contains a perfect seed, only their distribution matters.

An important consequence is that we can develop custom error models based on empirical estimates of the error distribution. This option is not only valid for modeling sequencing errors, but also for modeling differences occurring through biological processes, such as mutations or macroevolution.

We introduce error-only intervals, which will encapsulate the available information about the size of error patches. Treating deletions separately will allow us to simplify the exposition, so we will consider that error-only intervals are always non-empty.

#### Definition 7.

An **error-only interval** is a non-empty sequence of errors that cannot be extended left or right.

We assume that every nucleotide has a constant probability *q* of being correct. With probability *p* =1 – *q*, the nucleotide is incorrect and thus initiates an error-only interval. If *p*_*k*_ is the empirical probability that the error-only interval has size *k* for 1 ≤ *k* ≤ *n*, we can write the weighted generating function of error-only intervals as *pE*(*z*), where *E*(*z*) = *p*_1_*z* + *p*_2_*z*^2^ + … + *p*_*n*_*z*^*n*^.

The weighted generating function of deletions is denoted 5 for consistency with the previous sections. As before, we will ignore deletions adjacent to error-only intervals. Even if they occur, they have no consequence as they never interrupt a potential seed.

Reads under the empirical error model can be thought of as walks on the transfer graph shown in figure 12. An error-free interval can be followed by an error-only interval, or by another error-free interval if a deletion is present in between. An error-only interval can only be followed by an error-free interval.

The head and tail vectors are (*F*(*z*), *pE*(*z*))^⊤^ and (1,1), respectively; and *ψ*(*z*) = 1. The transfer matrix associated with the body of the transfer graph shown in figure 12 is

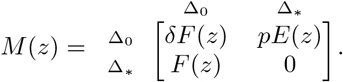

Computing *ψ*(*z*) + *H*(*z*) ^1^ · (*I* – *M*(*z*)) ^−1^ · *T*(*z*) as per proposition 4 yields

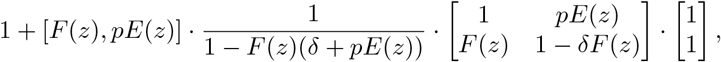

which eventually simplifies to

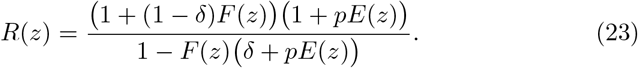

As previously, the weighted generating function of error-free intervals is *F*(*z*) = *qz* + (1 – *δ*)(*qz*)^2^ + (1 – *δ*)^2^(*qz*)^3^ + … = *qz*/(1 – (1 – *δ*)*qz*). Substituting this value in the equation above, we obtain

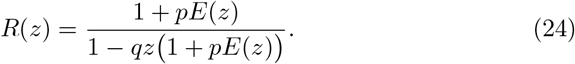

Epxression (24) is in general not equal to 1/(1 – *z*) because *E*(*z*) is not equal to *z*/(1 – *pz*). Since the total weight of reads of size *k* is not equal to 1, we cannot simplify ratio (9) as in the previous cases. We will need to include [*z*^*k*^]*R*(*z*) in our calculations, which we will approximate with proposition 5. This implies that we have to find the root with smallest *modulus* of the polynomial 1 – *qz*(1 + *δ* + *E*(*z*)).

**Figure 12:**
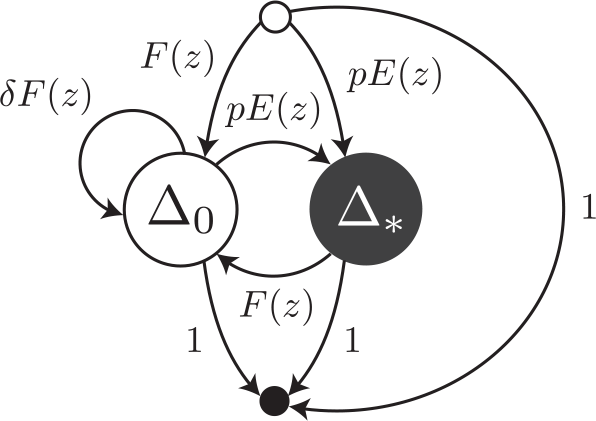
Transfer graph of reads under the empirical error model. Reads are viewed as sequences of error-free intervals (symbol Δ_0_) or error-only intervals (symbol Δ_∗_). An error-free interval can follow another one if a deletion is present in between. *F*(*z*) and *pE*(*z*) are the weighted generating functions of error-free intervals and error- only intervals, respectively. *δF*(*z*) is the weighted generating function of a deletion followed by an error-free interval.

#### Remark 16.

Why [*z*^*k*^]*R*(*z*) = 1 for some models but not for others? The combinatorial “atoms” can be inhomogeneous, leading to a weight deficit for certain sequence sizes. For instance, sequences of the object aa with weighted generating function *z*^2^ cannot have odd size. Empirical error-only intervals are typically bounded whereas error-free intervals are not. This imbalance introduces a weight deficit for some sequence sizes.

As in the previous sections, to find the weighted generating function of reads without an exact γ-seeds, we need to replace *F*(*z*) in expression (23) by its truncation *F*_γ_(*z*) = *qz* + (1 – *δ*)(*qz*)^2^ + … + (1 – *δ*)^γ−2^(*qz*)^γ−1^. The terms only partially cancel out and we obtain

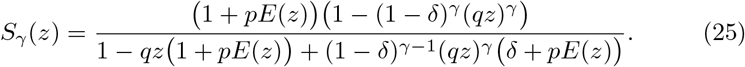

#### Remark 17.

When *R*(*z*) *and S*_γ_ (*z*) are defined by (24) and (25), ratio(9) is always less than or equal to 1. To see this, note that expression (23) can be written as

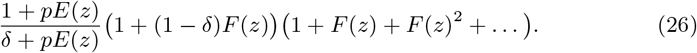

From this expression, it is clear that replacing *F*(*z*) by its truncation *F*_γ_ (*z*) decreases the coefficient of *z*^*k*^ (recall that all the weights are positive). This is equivalent to [*z*^*k*^]*R*(*z*) ≥ [*z*^*k*^]*S*_γ_(*z*), confirming that the probability of occurrence of reads without seed is always less than or equal to 1.

To estimate the total weight of reads without seed, we need to find the root with smallest *modulus* of 1 – *qz*(1 + *pE*(*z*)) + (1 – *δ*)^γ−1^(*qz*)^γ^(*δ* + *pE*(*z*)). The solution depends on the particular expression of *E*(*z*). Even though the process can be automated using proposition 5, every case is different and we cannot give an explicit formula here.

#### Example 17.

Assume that empirical measurements suggest that *q* = 0.9, *δ* = 0.15 and that error-only intervals of size 1, 2 or 3 have probabilities 0.5, 0.33 and 0.17, respectively (in this example, error-only intervals of size greater than 3 are never observed). This implies *E*(*z*) = 0.5*z* + 0.33*z*^2^ + 0.17*z*^3^.

If we choose seeds of size γ = 17, the weighted generating function of reads in expression (24) becomes

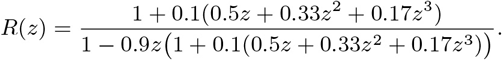

The smallest root of the denominator is approximately equal to 1.008754. The multiplicative constant of proposition 5 is approximately equal to 0.962590, so the total weight of reads of size *k* is approximately equal to 0.962590/1.008754^*k*+1^.

The denominator of (25) can be expressed in the form that is simpler to compute 1 – (1 – *δ*)*qz* – *qz*( 1 – (1 – *δ*)^γ−1^ (*qz*)^γ−1^) (*δ*+*pE*(*z*)). The weighted generating function of reads without seed becomes

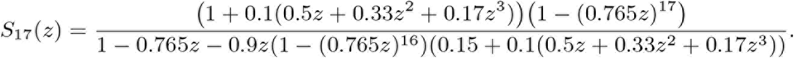

The smallest root of the denominator is approximately 0.0122100. The multiplicative constant of proposition 5 is approximately equal to 0.949377. So the total weight of reads without seed is approximately equal to 0.949377/1.012098^*k*+1^.

Combining these two results, the probability that a read of size *k* has no seed is the ratio of the total weights computed above. This is approximately equal to 0.9862735/1.003315^*k*+1^.

### 3.7 Worst case for approximations

So far, all the examples showed that the analytic combinatorics approximations are accurate. Indeed, the main motivation for our approach is to find estimates that converge exponentially fast to the target value. Does this mean that we can always use the approximations in place of the true values?

To find out, we need to describe the behavior of the estimates in the worst conditions. The approximations become more accurate as the size of the sequence increases, *i.e*. as the reads become longer. This is somewhat inconvenient: the read size is usually fixed by the technology or by the problem at hand, so the user does not have easy ways to improve the accuracy. Overall, the approximations described above just tend to be less accurate for short reads.

The second aspect is convergence speed. In proposition 5 it was shown that the rate of convergence is dominated by the ratio between the two smallest singularities of the weighted generating function. This means that convergence is fastest when the dominant singularity is significantly closer to 0 than the other singularities. Conversely, convergence is slowest when at least one other singularity is almost as close to 0.

The worst case for the approximation is thus when the reads are small and when the parameters are such that singularities have relatively close *moduli*. In the error model of uniform substitutions, this corresponds to small values of the error rate *p*.

To see this, recall that the singularities *Re*^*iθ*^ of *S*_γ_(*z*) expressed in (12) are bound by the equation

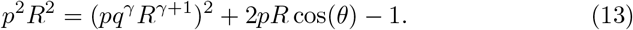

Considering this an equation of *R* > 0, the solution is minimal when 2*pR* cos(*θ*) is maximal, and *vice versa*. Thus, the two equations for the smallest and largest solutions 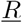 and R̅ are respectively

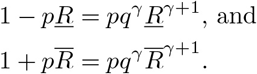

Observe that 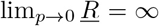, otherwise *z*(1 + *qz* + … + (*qz*)^γ−1^) would be bounded and the equality *Q*(*z*) = 1 – *pz*(1 + *qz* + … + (*qz*)^γ−1^) = 0 could not hold. This implies 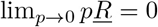 and 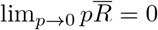. Otherwise, if we had for instance 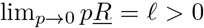, taking the limit of the first equation above would yield 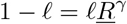, inconsistent with the fact that 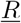 is unbounded.

This last fact entails 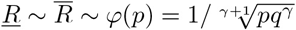. Indeed, we have 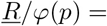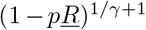 and 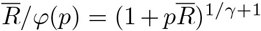, which both tend to 1 as *p* vanishes. This means that the *moduli* of all the singularities get closer to each other as *p* decreases. As a consequence, the speed of convergence of the approximation of proposition 5 diminishes (but it remains exponential).

In practical terms, the situation above describes the specifications of the Illumina technology, where errors are almost always substitutions, occurring at a frequency around 1% on current instruments. Since the reads are often around 50 nucleotides, the analytic combinatorics estimates of the seeding probabilities are typically less accurate than suggested in the previous sections.

Figure 13 shows the accuracy of the estimates in one of the worst cases. The curve is clearly distinct from the simulation at the chosen scale, but the absolute difference is never higher than approximately 0.015 (and lower for read sizes above 40). Whether this error is acceptable depends on the problem. Very often, the error on the measure of *p* is ±0.01 or greater, which is a more serious limitation on the precision than the convergence speed of the estimates. In most practical applications, the approximation error of proposition 5 can be tolerated even in the worst case, but it is important to bear in mind that it may not be totally negligible for reads of size 50 or lower.

If precision is crucial, remember that the exact solution can be found by using all the singularities of the weighted generating function *S*_γ_ (see remark 9). If computational speed is an issue, the singularities and associated proportionality constants can be precomputed for a range of values of *p* and γ.

**Figure 13:**
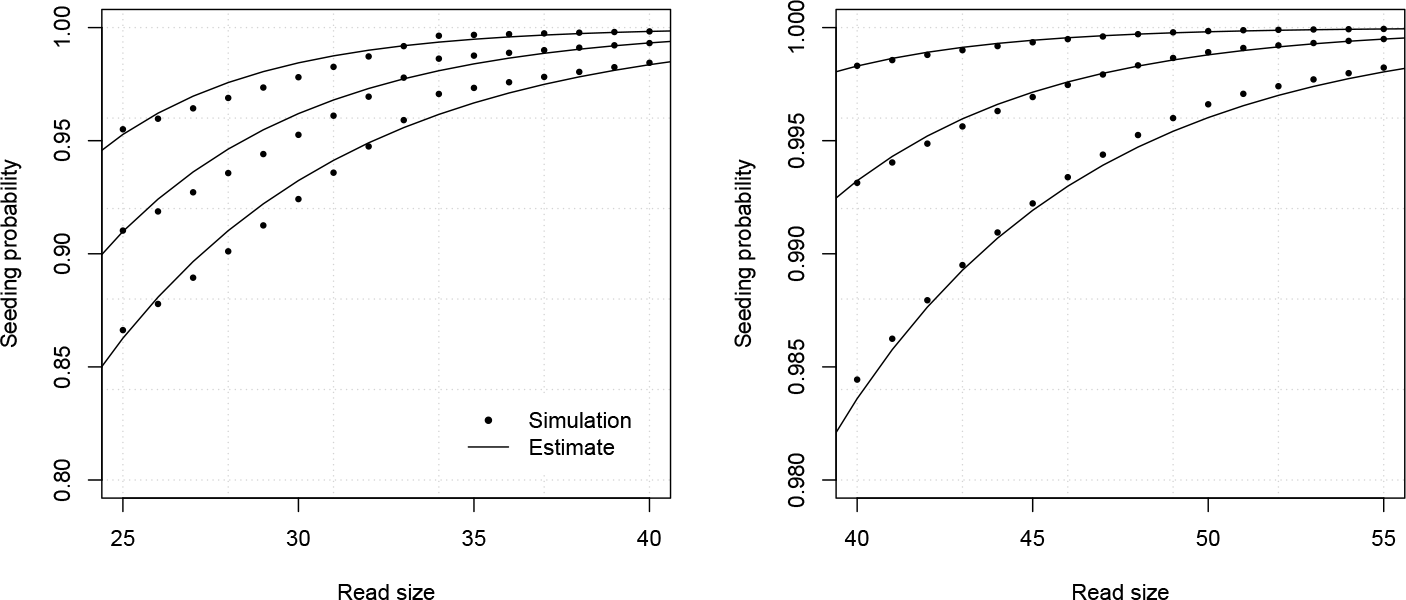
Example worst case for seeding with substitutions only. The analytic combinatorics estimates of proposition 7 are benchmarked against 10,000,000 random simulations. Shown on both panels are the probabilities that a read of given size contains a seed of size γ = 17, either estimated by random simulations (dots), or by proposition 7. The curves are drawn for *p* = 0.005, *p* = 0.010 or *p* = 0.015 (from top to bottom). The largest difference between the estimates and the simulations is around 0.015.

### 3.8 Oversimplified error models

It may be tempting to replace the somewhat complex error models of sections 3.4 and 3.5 by the simpler uniform subsitutions model of section 3.3 with an equivalent error rate. An intuitive approach would be to set the only parameter *q* = 1 – *p* of this model to the value of *q*(1 – *δ*) of the more complex ones, because those represent the probability of decoding a nucleotide without error.

However, this approach is inaccurate because insertions and deletions can have a strong influence. To show this, let us revisit example 16 with an approximate substitution model instead of the full error model.

#### Example 18.

In example 16, we computed the approximate probability that a read of size *k* = 100 contains no exact 17-seed, where *p* = 0.05, *δ* = 0.15, *r* = 0.05 and *r̃* = 0.45. In such conditions, the probability of decoding a nucleotide without error (given that it is not in an insertion burst) is (1 – *p* – *r*)(1 – *δ*) = 0.765. Now using a substitution model where *q* = 0.765 and thus *p* = 0.235, the analytic combinatorics estimate comes out as 1.03726/1.002591^101^ ≈ 0.79866. This number is outside the 99% confidence 0.77373 – 0.77376 and the uniform substitution model underestimates the probability that the read contains a seed by more than 2 percentage points in this case.

The approximation error does not vanish. Actually, in the conditions of example 18 it increases with the read size *k*. Figure 14 shows the same data as figure 11, now fitted by an approximate substitution error. The error can be as high as 5 percentage points.

**Figure 14:**
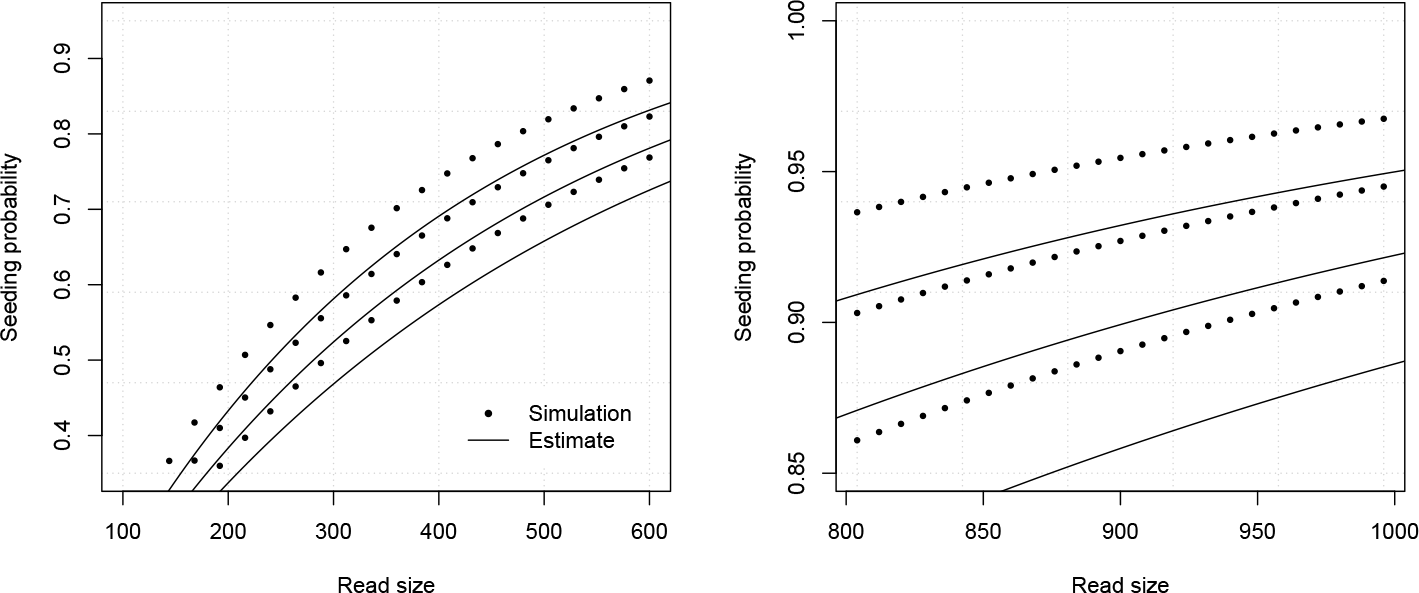
Example estimates with oversimplified error model. Shown on both panels are the probablities that a read of given size contains a seed. The dots are obtained by 10,000,000 random simulations of the full error model with γ = 17, *p* = 0.05, *δ* = 0.15, *r* = 0.45 and *r̃* = 0.04, *r* = 0.05 or *r* = 0.06 (from top to bottom)-they are the same as in figure 14. The lines represent the analytic combinatorics estimates of the oversimplified uniform substitution model computed from proposition 7 with *p* = 0.2265 or *p* = 0.2350 or *p* = 0.2435 (top to bottom).

The conclusion is that the quality of the approximation collapses when using approximate error models. One may argue that an error of 5 percentage points may be tolerable, but the actual amount of error when using an approximate error model is unknown in genereal. Using simplified error models is unsafe because convergence is slow (the term “convergence” can still be used because the probability that the read contains a seed always tends to 1 as the size increases) and the error cannot be controlled.

Nevertheless, complexity comes at a cost. It is important to remember that the parameters of the error model may be inaccurate, in which case the error is also not controlled. If these parameters are hard to estimate, it may be preferrable to use a simplified model. But in general, the amount of data available in a sequencing run is sufficient to estimate the parameters of the error model. In practice, the safest approach to compute seeding probabilities is to use the full error model described in section 3.5 and estimate the four parameters from the alignment data generated during the mapping. If *δ* or *r* are small, they may be set to 0 for simplicity, yielding the error models of section 3.4 or section 3.3 (see remark 15).

## 4 Advanced seeding methods

Until now we have seen models of increasing complexity, but the transfer graphs had a fixed layout, and the weighted generating functions were ratios of polynomials of degree γ, the minimum size of the seed (with the notable exception of the empirical error model, for which the degree is not specified *a priori*). In relation to different problems, we will now explore other categories of models where the layout of the transfer graph depends on γ, and where the degree of the polynomials to solve will increase much faster than γ.

Because of this increased complexity, we will only consider the simplest uniform substitution error model. Otherwise, the strategy remains unchanged: we will formulate combinatorial problems, view their solutions as walks on transfer graphs, encode these graphs as transfer matrices, obtain the weighted generating functions of the solutions from proposition 4 and approximate the answers using proposition 5.

### 4.1 Inexact seeds

We now consider a more challenging problem. The ongoing development of algorithms and data structures makes it possible to search inexact seeds, *i.e*. sequences that are very similar but not identical to the target. This comes at a greater computational cost than finding exactly similar sequences. If this cost is mitigated, it may be worth searching inexact seeds because the chances are higher to identify the target.

The theory below is implemented within the framework of the uniform substitutions error model. Adaptations to more complex error models do not present new theoretical challengnes, but the many cases to consider make the models cumbersome, at the cost of concision and clarity. Let us begin with a definition that will simplify the following discussion.

#### Definition 8.

**A single substitution interval** is a non-empty sequence of nucleotides that cannot be extended left or right, and that contains at most one substitution and no other error. An **inexact γ-seed** is a single substitution interval of size γ or greater.

We emphasize once more that our concern is the case of *one* substitution; the cases with more errors, or with errors of other types are not considered. An exact γ-seed is an inexact γ-seed, but the converse is not true. Figure 15 illustrates the relationshiop between substitutions and single substitution intervals. Note that single substitution intervals can overlap. In the models below, this is something that we will have to account for explicitly.

We already computed the weighted generating function of reads under the uniform substitutions model in section 3.3. Namely, in equation (11) we found *R*(*z*) = 1/(1 – *z*) = 1 + *z* + *z*^2^ + … from which we concluded that the total weight of reads of size *k* ≥ 0 is 1. We now need to find the weighted generating function of reads that do not contain an inexact γ-seed. For this, we introduce a new type of combinatorial object.

**Figure 15:**
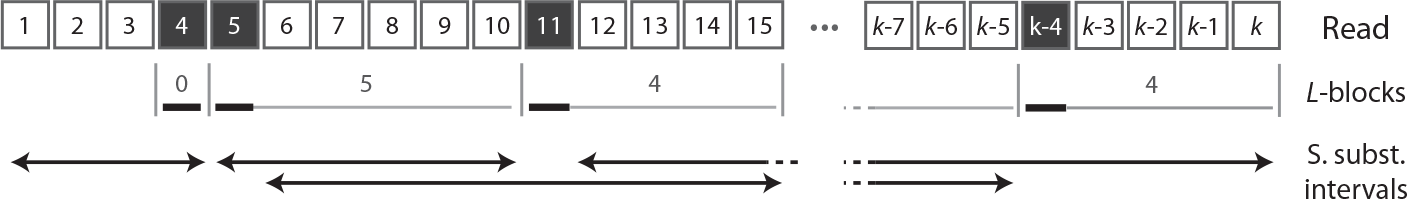
Inexact seeding. Substitutions (black squares) occur uniformly at random within the read. They delimit L-blocks and single substitution intervals. L-blocks consist of a substitution followed by an error-free interval. There is always a substitution on the left of an L-block: the error-free interval at the head of the read is not an L-block. Single substitution intervals (arrows) are the longest stretches of the read that contain at most one sbustitution.

#### Definition 9.

An **L-block** is a substitution followed by an error-free interval.

The point of this definition is that a read is an error-free interval, followed by a sequence of *L*-blocks. In a read without inexact γ-seeds, not all the combinations of *L*-blocks are possible. For instance, an *L*-block with γ – 2 correct nucleotides can only be followed by a substitution, *i.e*. an *L*-block with 0 correct nucleotide (otherwise the concatenation of these two *L*-blocks forms an inexact γ seed). A substitution with γ – 3 correct nucleotides can be followed by an *L*-block with 0 or 1 correct nucleotide, *etc*.

To give a concrete example, reads without inexact 5-seed can be seen as walks on the graph shown in figure 16. The numbers on the vertices indicates the amount of correct nucleotides in an *L*-block. There cannot be an *L*-block with 4 or more correct nucleotides, otherwise the read would contain an inexact 5-seed. The edges capture the relationships between consecutive *L*-blocks. The weighted generating function of *L*-blocks with i correct nucleotides is designated by *ℓ*_*i*_ (*z*), for *i* = 0,1, 2, 3.

The head and tail vectors are shown on the right panel of figure 16. Reads can start with up to 3 correct nucleotides. The weighted generating function of error-free intervals of size *i* is designated *f*_i_(*z*), for *i* = 0,1, 2, 3. Finally, any *L*-block can mark the end of the read, so all of them are connected to the tail vertex by 1, the weighted generating function of the empty oject *ε*. We do not need to connect the head and the tail node with such an edge, because as we will see below, the empty sequence is already described by the transfer graph.

The weighted generating function of an *L*-block with i correct nucleotides is *ℓ*_i-block_(*z*) = pz × *qz* × … × *qz* = *pz*(*qz*)^*i*^, *i.e*. the weighted generating function of a substitution followed by *i* correct nucleotides. In a similar way, the weighted generating function of a stretch of *i* correct nucleotides is *f*_*i*_(*z*) = *qz* × … × *qz* = (*qz*)^*i*^.

Because *f*_0_(*z*) = 1, we now see that on the graph shown in figure 16, a walk starting from the head vertex and going to the tail vertex through the vertex labelled 0 has weighted generating function 1. This corresponds to the empty sequence, so there is no need to add it again to the graph, so *ψ*(*z*) = 0.

In the general case γ > 0, the matrix of the body of the transfer graph shown in figure 16 is

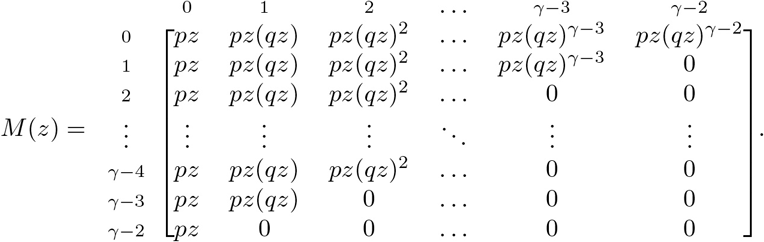

The head vector *H*(*z*) is equal to (1, *qz*, (*qz*)^2^,…, (*qz*)^γ−2^)^⊤^, the tail vector *T*(*z*) is equal to (1,1,…, 1)^⊤^, and *ψ*(*z*) = 0. The expression of the weighted generating function of reads without inexact γ-seed is omitted here. By proposition 4, it can be computed as *S*_γ_(*z*) = *H*(*z*) · (*I* – *M*(*z*))^−1^ · *T*(*z*).

**Figure 16:**
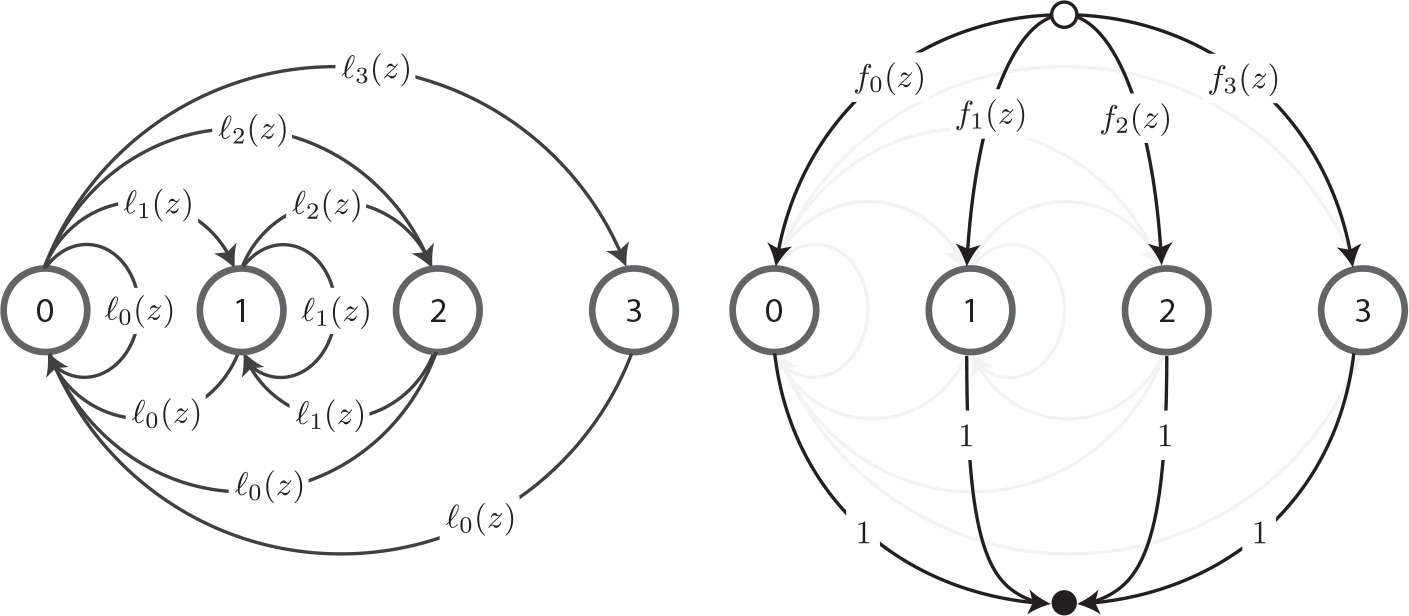
Transfer graph of reads without inexact 5-seed. Reads are viewed as sequences of *L*-blocks. To not overload the figure, the body of the transfer graph is shown on the left, and the head and tail edges are shown on the right. The labels on the vertices represent the number of correct nucleotides in the *L*-blocks. The terms *ℓ*_*i*_(*z*) are the weighted generating functions of *L*-blocks with *i* correct nucleotides for *i* = 0,1, 2, 3. The terms *f_i_(z*) are the weighted generating functions of error-free intervals of size i = 0,1, 2, 3.

#### Example 19.

Let us revisit example 14, where we approximated the probability that a read of size *k* = 100 has no seed for γ = 17 and for *p* = 0.1; but this time we allow the seed to contain one substitution. The matrix *M*(*z*) has dimension 16 and the weighted generating function *S*_17_(*z*) can be written as *P*(*z*)/*Q*(*z*), where *P* and *Q* are polynomials. Using numerical methods to find the root of *Q* with smallest *modulus*, we obtain *z*_1_ ≈ 1.079244. Likewise, we obtain —*P*(*z*_1_)/*Q*^/^(*z*_1_) ≈ 2.326700, so the probability that the read does not contain an inexact seed is approximately 2.326700/1.079244^101^ ≈ 0.0010511. For comparison, a 99% confidence interval obtained by performing 10 billion random simulations is 0.001049 – 0.001054. Recall that in the same conditions, the chances that the read does not contains an exact seed were approximately 9.6%.

The approximations have the typical accuracy of analytic combinatorics estimates. Figure 17 shows the precision of the estimates for different values of the substitution rate *p* and of the read size *k*. Observe that the probability that the read contains an inexact seed is substantially higher than the probability that it contains an exact seed (compare with figure 7).

**Figure 17:**
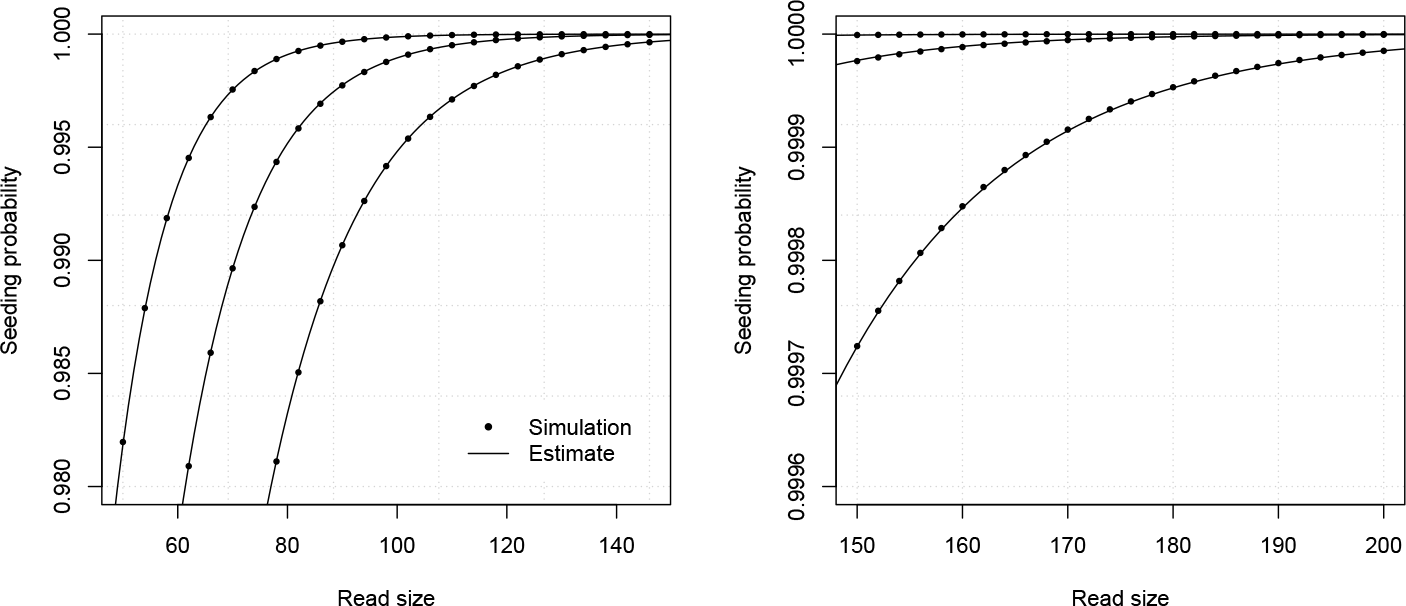
Example estimates for inexact seeding with substitutions only. The analytic combinatorics estimates described in this section are benchmarked against random simulations. Shown on both panels are the probabilities that a read of given size contains an inexact seed, either estimated by random simulations (dots), or by the method described above (lines). The curves are drawn for γ = 17, *p* = 0.08, *p* = 0.10 or *p* = 0.12 (from top to bottom). 10,000,000 simulations are run on the left panel and 100,000,000 on the right one.

Unfortunately, the expressions of the weighted generating functions given by proposition 4 are very cumbersome, and they may take a long time to evaluate. For instance, the degrees of *P* and *Q* for γ = 17 are 135 and 136, respectively.

An option is to precompute the dominant singularities and associated multiplicative constants for a useful range of values of γ (up to 30 is sufficient for most applications) and of the parameter *p* (up to 0.25 is sufficient for most applications). Once these values are stored, the estimates can be calculated rapidly.

### 4.2 Type I errors

Our concern so far was whether reads contain a seed, *i.e*. an error-free (or almost free) interval of sufficient size. If this is the case, the target sequence is among the candidates and we implicitly assume that it will be identified during the alignment stage. But we did not discuss what happens if the read does not contain a seed.

There are two possible cases. The first is that the mapping procedure returns no hit. The conclusion is then that the sequence does not belong to the genome, or that it cannot be mapped. In analogy with statistical testing, we will call this a “type II error”. The second case is that the mapping procedure returns a hit. This cannot be a true hit because the target was discarded during the seeding stage. We will call this a “type I error”. Type I errors are the most difficult to detect, because nothing distinguishes them from true hits, at least at the seeding stage.

It is important to emphasize that in this version of the mapping problem, the sum of type I and type II error rates is the probability that the read does not contain a seed (because we assume that there is a mapping error if and only if the read does not contain a seed). The split between type I and type II errors depends on the genome, and more particularly on the configuration of its repeated sequences. A stretch of γ nucleotides matches a random genome with probability *O*(4 ^−γ^). However, genomes are not a succession of random nucleotides. Large sequences are often duplicated, to the extent that the greater part of eukaryotic genomes can be considered repetitive. In addition, repeats are usually not exactly identical, but only similar. This means that the *O*(4 ^−γ^) term can be spectacularly underestimated.

A complete treatment of type I errors is presently not possible. We will focus on a single case that will be the basis for further development. More specifically, we will assume that the target sequence has exactly one duplicate in the genome, differing only by uniform substitutions (we will see below how to relax these assumptions). The problem then amounts to finding the probability that the duplicate but not the target sequence is discovered during the seeding step. Figure 18 shows how this can occur.

**Figure 18:**
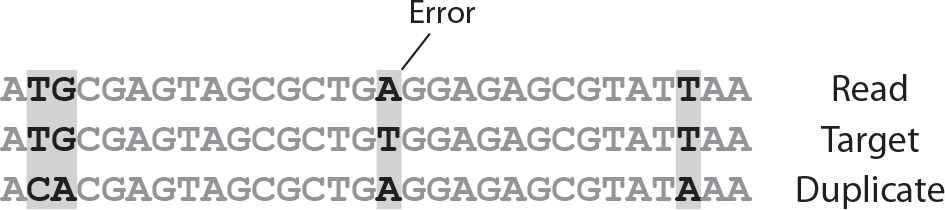
Example type I error. A read of 30 nucleotides contains a single error at the sixteenth position. By chance, the erroneous nucleotide is identical to the duplicate sequence at this position. This creates a match of 24 nucleotides for the duplicate sequence, while the longest match for the target is 15 nucleotides. Thus, using seeds of size greater than 15 would result in a type I error (even though the duplicate has three differences with the read and the target has only one).

#### Remark 18.

With only one reference sequence, “error-free” and “matching” nucleotides are the same. With more than one reference, we must take care of distinguishing the two, because matching nucleotides can be incorrect (they can match any of the duplicates).

Instead of tackling this problem directly, we will set a more general framework where a read can match two similar sequences denoted (+) and (−). A read now consists of four kinds of nucleotides: those matching both sequences, those matching the (+) sequence only, those matching the (−) sequence only, and those matching none. The first kind will be referred to as a *match* and the last three as *mismatches* (single or double). The following definitions will be useful to sketch the transfer graph.

#### Definition 10.

An **R-block** is a mismatch-free interval followed by a single or a double mismatch. *A* (+) **interval** (respectively (−) **interval**) is a sequence of nucleotides matching the (+) sequence (respectively (−) sequence) that cannot be extended left or right. *A* (+) **seed** (respectively (−) **seed**) is a (+) interval (respectively (−) interval) of size γ or greater.

This information is summarized in figure 19. We will use it to find a construction for reads with neither (+) nor (−) seed. We will later show how this can be used to find the probabilities of type I errors.

**Figure 19:**
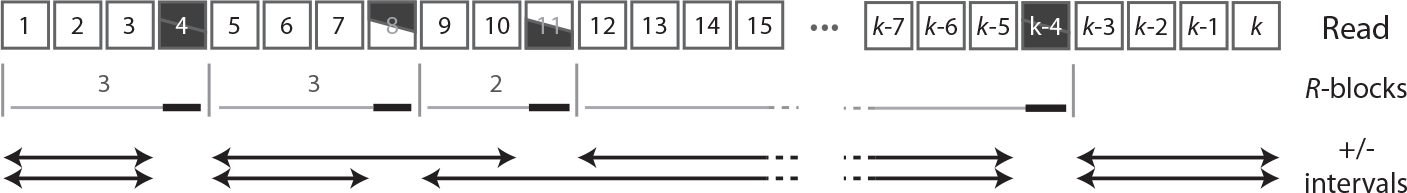
Seeding with two reference sequences. Matches are represented as white squares, mismatches against the (+) sequence as bottom black wedges, mismatches against the (−) sequence as top black wedges, and double mismatches as black squares. They delimit R-blocks, consisting of mismatch-free intervals followed by a mismatch. The last nucleotide of an R-block is always a mismatch: the mismatch-free interval at the tail of the read is not an R-block. (+) or (−) intervals are stretches of the read that match either the (+) sequence (top arrow) or the (−) sequence (bottom arrow) and that cannot be extended left or right.

#### Remark 19.

We consider that when a subsequence is both a (+) and a (−) seed (as the head of the read shown in figure 19 for instance), both sequences are in the list of candidates after seeding, i.e. both are aligned.

Reads can be viewed as sequences of *R*-blocks followed by a mismatch-free interval. Reads with neither (+) nor (−) seed have particuarly complex constraints on the arrangements of those *R*-blocks. For instance, an *R*-block of size γ (*i.e*. γ – 1 matches followed by a mismatch) must be followed by a mismatch if the final nucleotide is a single mismatch, but it can be followed by up to γ – 1 matches if this nucleotide is a double mismatch.

For the sake of clarity, we will consider a concrete example where γ = 3. The read below starts with two matches, followed by a double mismatch, followed by a mismatch against the (−) sequence. The numbers between brackets (0,1) indicate the respective amount of nucleotides matching the (−) and the (+) sequences at the end of the *R*-block. At that point, there are only five possiblities for the next *R*-block; every other combination would create a (+) seed or a (−) seed.

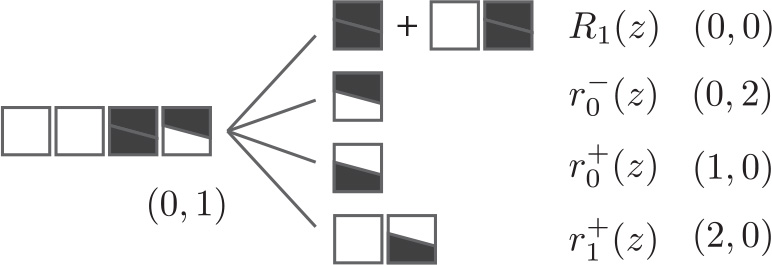

Appending a double mismatch or a match followed by a double mismatch (top case) brings the number of nucleotides matching (−) and (+) at the end of the *R*-block to (0, 0). The weighted generating function of this union of *R*-blocks is denoted *R*_1_(*z*). Appending a mismatch against the (−) sequence (second case from the top), *i.e*. an *R*-block with weighted generating function 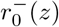 brings the amount of matching nucleotides to (0, 2). Conversely, appending a mismatch against the (+) sequence (third case from the top), *i.e*. an *R*-block with weighted generating function 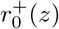 brings the amount of matching nucleotides to (1,0). Finally, appending a match followed by a mismatch against the (+) sequence (bottom case), *i.e*. an *R*-block with weighted generating function 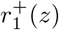 brings the amount of matching nucleotides to (2, 0).

By considering all the possible scenarios for other reads, we obtain the transfer graph shown in figure 20. Even for such a small value of γ, the graph is so dense that the edges have been separated in two panels and encoded as symbols in order to not overload the figure.

As in the example above, the numbers in the vertices indicate the amount of nucleotides matching (−) and (+) at the end of the *R*-blocks. Since *R*-blocks are terminated by a mismatch, at least one of these numbers is 0. The body of the transfer graph is only partially represented on the left panel. The edges pointing to the vertex (0,0) are displayed on the right panel, together with the head and tail edges. The four kinds of edges on the left panel correspond to different *R*-blocks with weighted generating functions 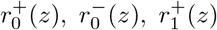 and 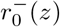. The sign indicates which sequence is *mismatched* at the end of the *R*-block and the index is the number of matches in the *R*-block.

The edges shown on the right panel for lack of space on the left panel all point to the vertex (0,0). All the vertices are connected to this vertex because appending an *R*-block terminated by a double mismatch always brings the read to this state. The weighted generating functions *R*_0_(*z*), *R*_1_(*z*) and *R*_2_(*z*) represent such *R*-blocks with *up to* 0, 1 or 2 matches, respectively.

The remaining edges shown on the right panel are the head and tail edges. In the initial state of the read, the amount of matching nucleotides is (0,0), which is indicated by the label 1 on the edge between the head vertex and the vertex (0, 0). This is equivalent to prepending the read by the empty object *ε*. Reads are terminated by a sequence of matches. The weighted generating functions *F*_0_(*z*), *F*_1_(*z*) and *F*_2_(*z*) represent mismatch-free intervals of size *up to* 0, 1 or 2, respectively. The empty sequence is already present in the graph (take a path from the head vertex to the tail vertex through (0, 0) with no match appended at the tail), and no extra sequence needs to be added, so here *ψ*(*z*) = 0.

**Figure 20:**
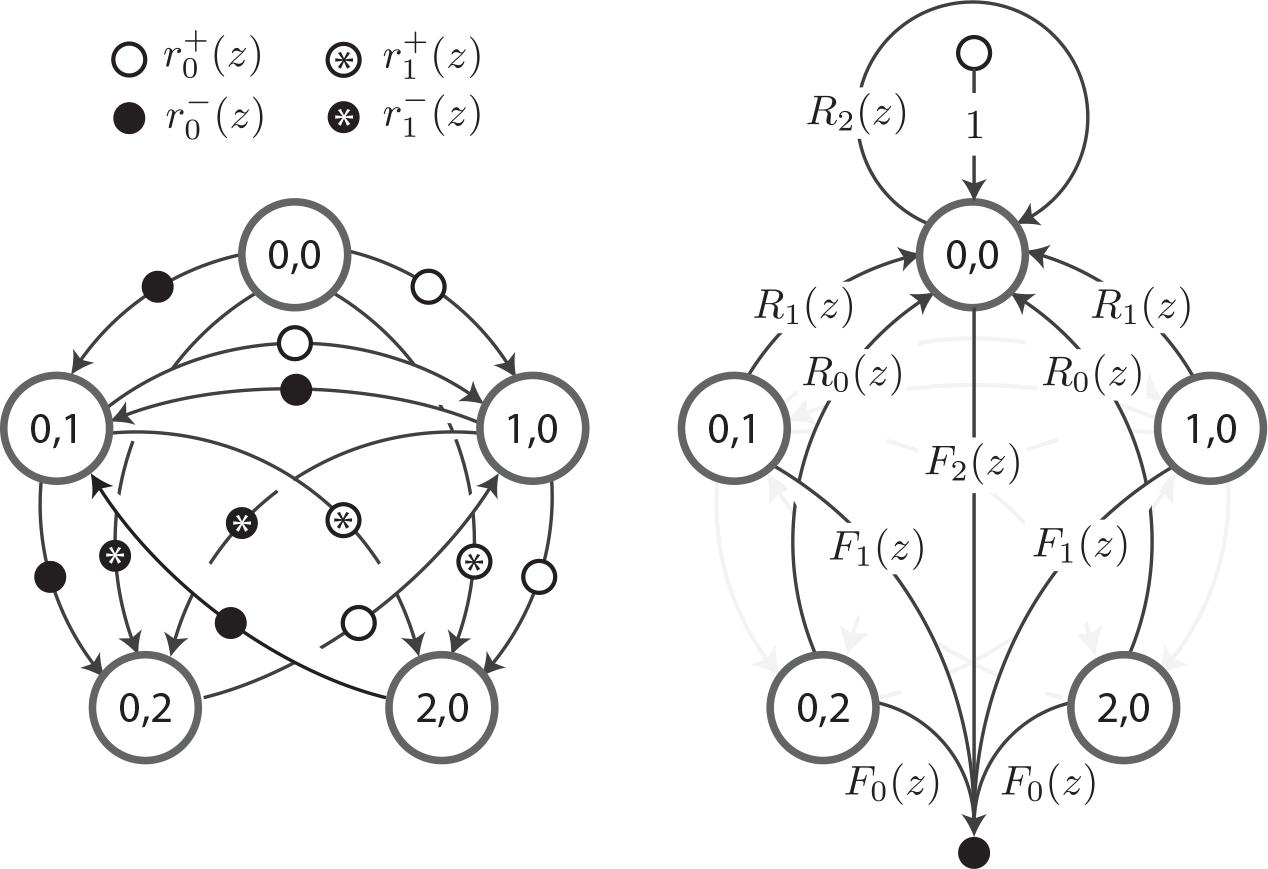
Transfer graph of reads without seed of size 3 for either (+) or (−). Reads are viewed as sequences of *R*-blocks, and they can match two sequences referred to as (+) and (−). To not overload the figure, the edges are split between the left and right panels. The numbers in the vertices represent the respective number of nucleotides matching the (−) and (+) sequences at the end of an *R*-block. Weighted generating functions r+ (z) represent *R*-blocks with i matches terminated by a mismatch against the (+) sequence (symmetrically for the (−) sequence). Weighted generating function *R*_*i*_(*z*) represent *R*-blocks with *up to i* matches terminated by a double mismatch. Weighted generating functions *F*_*i*_(*z*) represent mismatch-free intervals of *up to i* matches.

A similar logic can be applied for higher values of γ. The transfer graph becomes too cumbersome to represent but the transfer matrix has enough regularity to be specified in full. The general transfer matrix associated with the body of the transfer graph has dimension 2γ – 1 and is defined as

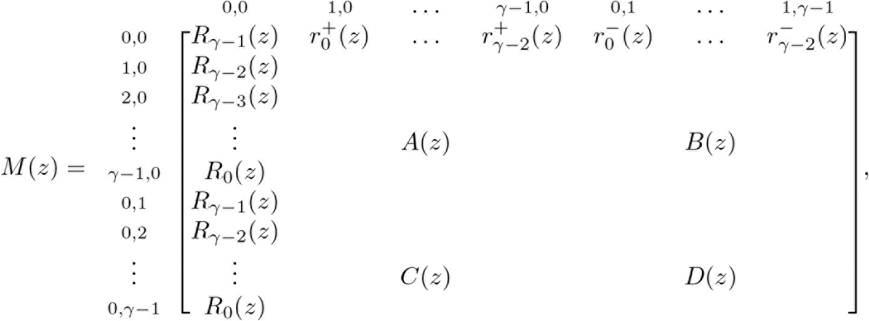

where *A*(*z*), *B*(*z*), *C*(*z*) and *D*(*z*) are γ – 1 × γ – 1 matrices defined as

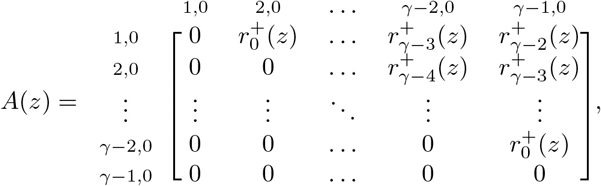

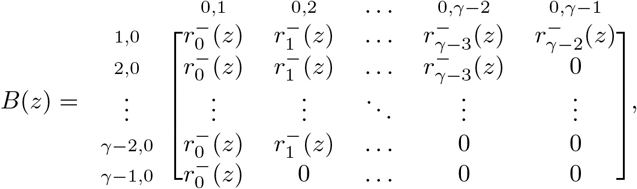

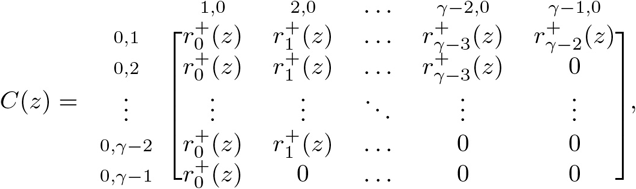

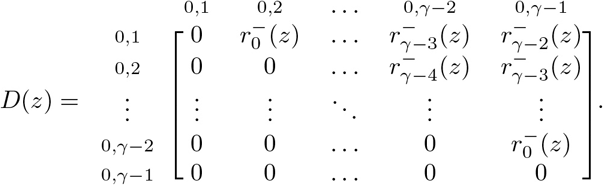

In this case the head vector *H*(*z*) is equal to (1, 0, 0,…, 0)^⊤^ and the tail vector *T*(*z*) is equal to (*R*_γ−1_(*z*), *R*_γ−2_(*z*),…, *R*_0_(*z*), *R*_γ−2_(*z*),…, *R*_0_(*z*))^⊤^. The expression of the weighted generating function of reads with neither (+) nor (−) seed is omitted here (it is fairly complex even for small values of γ). As per proposition 4, it can be computed as *H*(*z*)^⊤^ · (*I* – *M*(*z*))^−1^ · *T*(*z*).

We still need to give the exact expression of the weighted generating functions appearing in the definition of *M*(*z*). Under the assumption that mismatches occur uniformly in the read, nucleotides of each type occur with probabilities *a*, *b*, *c* or *d*, with *a* + *b* + *c* + *d* = 1, where ita is the probability that the nucleotide is a (double) match, *b* that it is a mismatch against (+), *c* that it is a mismatch against (−) and *d* that it is a mismatch against both. With these notations we obtain

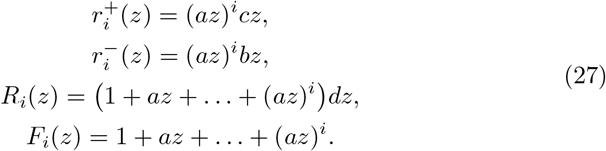

With these definitions we could find the expression of the weighted generating function of the reads with neither (+) nor (−) seed, but we will return to the original problem of computing the probability of a type I error. In this context, the (+) sequence is the target and the (−) sequence is the duplicate.

We will assume that each nucleotide is decoded incorrectly with probability *p* (substitutions due to sequencing errors) and that for each nucleotide, the duplicate sequence differs from the target with probability *κ* (divergence between the target and the duplicate). A nucleotide is a mismatch for both sequences if it is a read error (probability *p*) and if either the duplicate is identical to the target (probability 1 – *κ*) or if it is different from both the target and the decoded nucleotide (probability 2*κ*/3). The other probabilities can be computed with similar arguments, yielding *a* = (1 – *p*)(1 – *κ*), *b* = *pκ*/3, *c* = (1 – *p*)*κ* and *d* = *p*(1 – *κ*/3).

The model is completely specified by *p* and *κ*. With their actual values at hand, we can use proposition 5 to obtain an asymptotic estimate of the probability that a read has no match for the target *and* has no match for the duplicate. Call this estimated probability *P*_2_. Using the results of section 3.3, we can also compute an asymptotic estimate of the probability that the read does not contain an exact γ-seed (irrespective of potential matches to the duplicate). Call this estimated probability *P*_1_. We can thus estimate the probability that the read contains no exact γ-seed *but* contains a match for the duplicate as *P*_1_ – *P*_2_.

To clarify this equality, call *A* the event that the read has no γ-seed and *B* the event that it contains no match of size γ for either the target or the duplicate sequence. First observe that *B* ⊂ *A* so *P*(*A* ⋂ *B*^c^) = *P*(*A*) – *P*(*B*). Then, *B*^*c*^ is the event that the read contains an exact γ-seed, a stretch of size γ or greater matching the duplicate sequence, or both. Another way to describe *B*^*c*^ is that “some hit is found in the read”. So *A* ⋂ *B*^*c*^ is the event that “some hit is found, but it is not the target”, which is the definition of a type I error. Since *P*(*A*) ≈ *P*_1_ and *P*(*B*) ≈ P_2_, the probability of type I errors is indeed approximately equal to *P*_1_ – *P*_2_.

#### Example 20.

Let us approximate the probability of type I error (with one duplicate) for a read of size *k* = 100 with *γ* = 17 and for substitution rates *p* = 0.10 and *κ* = 0.10. In example 14 we found *P*_l_ ≈ 1.396145/1.0268856^101^. To compute *P*_2_, we substitute *a* = 0.81, *b* = 0.09, *c* = 0.00333, *d* = 0.09667 in the expression of the 33 × 33 transfer matrix *M*(*z*) and in the expression of *T*(*z*). We then compute the expression *H*(*z*)^⊤^ · (*I* – *M*(*z*))^−1^ · *T*(*z*) and obtain a rational function *P*(*z*)/*Q*(*z*) where *P* and *Q* are polynomials of degree 288 and 289, respectively. The root of *Q* with smallest *modulus* is the dominant singularity *z*_*L*_ of the weighted generating function. Using numerical approaches, we find *z*_*L*_ ≈ 1.0272930245. We compute the proportionality constant of proposition 5 as – *P*(*z*_1_)/*Q*Ⅎ(*z*_1_) ≈ 1.4032791 from which we obtain *P*_2_ ≈ 1.4032791/1.0272930245^101^. The type I error rate is then approximately equal to 0.0032906. For comparison, a 99% confidence interval obtained by performing 10 billion random simulations is 0.003288 – 0.003292.

Figure 21 illustrates the precision of this estimate for different values of the substitution rate *κ* and of the read size *k*. Observe that the estimates are accurate even for such low probability of occcurrence.

Computing *P*_2_ with proposition 5 requires finding a weighted generating function *P*(*z*)/*Q*(*z*) where *P* and *Q* are polynomials with too many terms to fit in this document. It is thus problematic to compute the approximations efficiently. The best option is to precompute the dominant singularities and associated multiplicative constants for a useful range of γ (up to 30 is sufficient for most applications) and of the parameters *p* and *κ* (up to 0.25 is sufficient for most applications). Once these values are stored, the estimates can be calculated rapidly from expression (5) without having to compute any weighted generating function.

At the start of this section, we assumed that the target has exaclty one duplicate. Also, *κ* is usually unknown, so are the estimates derived above useful at all? First, it is important to stress that the seeding strategy is efficient only when the target has few duplicates. Some sequences have more than 100,000 copies in the human genome. In such cases, the seeding process yields many candidates that will cost time at the alignment stage, with anyway small chances of identifying the correct target. As a consequence, most mapping algorithms bail out as soon as they gather enough evidence that the target has many duplicates. There is thus little incentive to develop accurate estimates of type I error rates for highly repeated sequences, and one should focus on the cases of intermediate amount of duplicates, say around 20.

**Figure 21:**
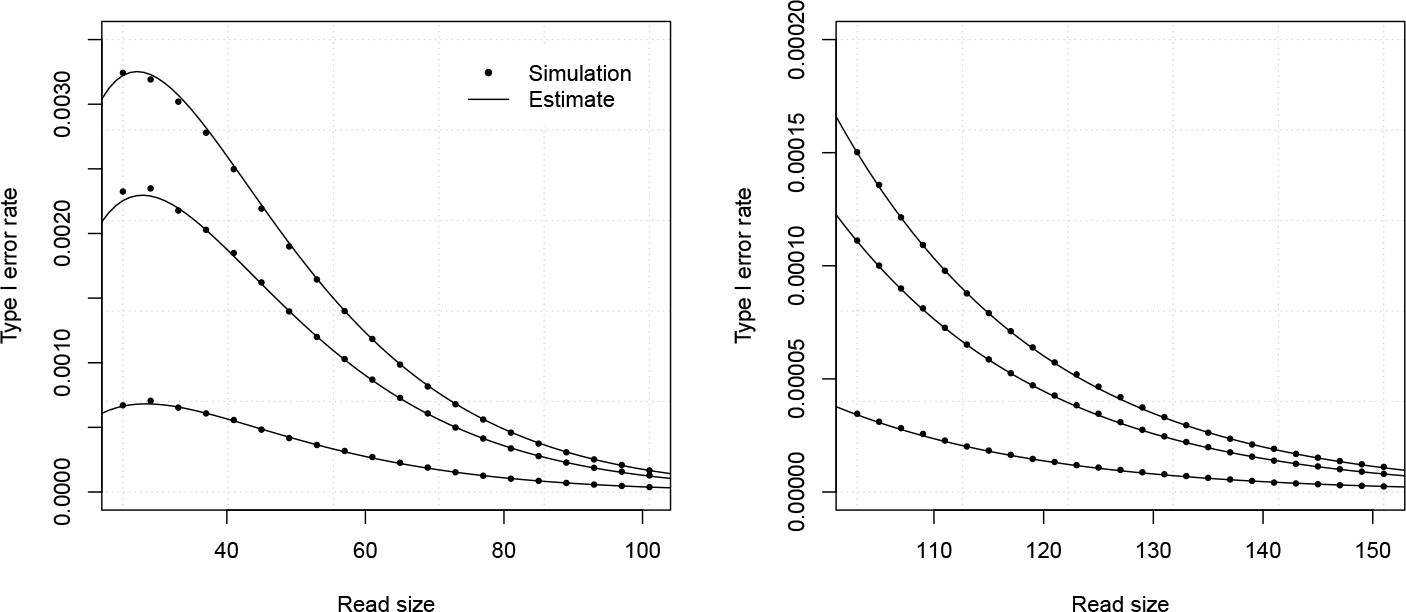
Example estimates of type I errors (one duplicate). The analytic combinatorics estimates are benchmarked against random simulations. Shown on both panels are the probabilities that a read of given size will induce a type I error, either estimated by 10,000,000 (left) or 100,000,000 (right) random simulations (dots), or by the method described above (lines). The curves are drawn for γ = 17 and *p* = 0.05, *κ* = 0.05, *κ* = 0.15 or *κ* = 0.25 (from top to bottom).

Fortunately, we can extend the theory developed above to the case of a few duplicates. The probability that a read contains no match for the duplicate given that it contains no match for the target is *P*_2_/*P*_1_. If the target has *N* extra duplicates evolving independently with mutation rate *κ*, the probability that a read contains no match for any of the duplicates given that it contains no match for the target is (*P*_2_/*P*_1_)^*N*^. The probability of a type I error is thus *P*_1_(1 – (*P*_2_/*P*_1_)^*N*^). For *N* = 1, we recover *P*_1_ – *P*_2_ and for large *N*, the value approaches *P*_1_, the probability that the read has no seed.

This approach is somewhat naive because repeated sequences do not evolve independently of each other and *N*, like *κ* is usually unknown. However, it gives a handle on the problem as some “proxy” *N* and *κ* can be estimated during the seeding step or better, stored for every position of the genome.

An additional observation is that the case of one duplicate gives a lower bound for the probability of type I error. When there is exactly one duplicate, type I errors are almost impossible to detect because the mapping process identifies a single hit that is similar to the read (as if a seed were present and the target had no duplicate). If the proportion of the genome that is repeated is *x* and *P*\* is the maximum of *P*_1_ – *P*_2_ relative to unknown *κ*, then *xP*\* is a reasonable lower bound on the probability of type I errors. This number is an *a priori* estimate that depends only on the read size and on the sequencing error rate. If more information about the read, or about the seeding and alignment processes is available, then this lower bound can vary.

#### Example 21.

Let us approximate the *a priori* lower bound of the type I error rate for reads of size *k* = 100 with γ = 17 and for sequencing error rate *p* = 0.10. In example 14 we found *P*_1_ ≈ 1.396145/1.0268856^101^, which does not depend on *κ*. We now need to compute the lower bound of *P*_2_ as *κ* varies. Using precomputed values, we find the minimum is reached for *κ* ≈ 0.07 where *P*_2_ ≈ 1.403628/1.02732^101^. This yields an extimate approximately equal to 0.0035. Assuming that approximately half of the genome has at least one duplicate sequence, we cannot guarantee a type I error rate below 0.17% with this seeding strategy.

The methods presented in this section are numerically accurate, but the models may not be faithful to the biological reality. It is still challenging to represent a genome together with the relationships between its duplicated sequences, and more generally to know the number of duplicates of a target sequence. In summary, estimates of the type I error rate are strongly dependent on our knowledge of the repeat structure of the genome. New data structures or algorithms to model genomic repeats will greatly benefit seeding heuristics in the mapping problem.

### 4.2 MEM seeds

In the exact seeding scheme considered so far, every subsequence of the genome that matches γ consecutive nucleotides of the read is considered a candidate at the seeding step. While assuring good chances of discovering the target, this approach often yields “too many” useless candidates, which costs computational time at the alignment step. An alternative scheme called MEM seeding (Maximal Exact Match) gives better empirical results. The principle is to use as seeds only the longest local matches between the read and the genome.

#### Definition 11.

A γ-MEM (Maximal Exact Match) is a sequence of at least γ nucleotides from the read, that matches a subsequence of the genome and that cannot be extended left or right.

#### Remark 20.

Note that error-free intervals and mismatch-free intervals are disjoint, whereas MEMs can be overlapping (and usually are).

Since there are fewer MEM seeds than exact γ-seeds, MEM seeding is more prone to errors than exact seeding. As we will see below, an important difference is that the combined frequencies of type I and type II errors is strictly greater than the probability that the read contains an error-free interval of size 7 or greater.

To make the problem more concrete, we will consider as in section 4.2 that the target has exactly one duplicate sequence. In a similar way, we will start by defining two potential matches referred to as (+) and (−). The definitions of *R*-block, (+) interval and (−) interval given in section 4.2 still apply.

Figure 22 highlights the properties of MEMs. Notice how a match for (+) is masked by a longer match for (−). Nucleotides from 12 to 16 match the (+) sequence, creating a potential seed. However, it is contained in a match for the (−) sequence from nucleotides 9 to 20. Since the match for (+) is not a MEM, it is not used as a seed. Cases such as this can cause type I errors to occur even when the reads contains matches of size 7 or greater for the target.

**Figure 22:**
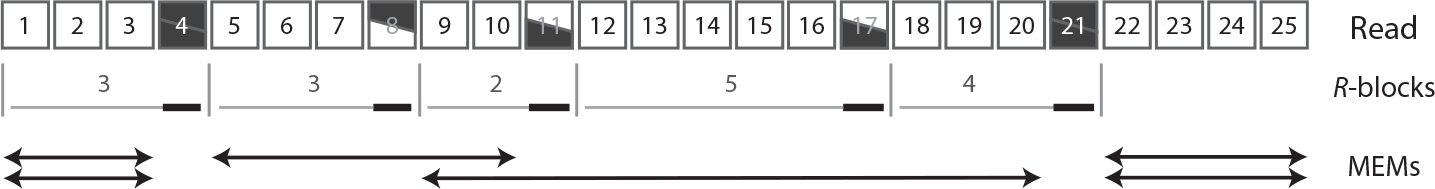
MEMs (with two references). The symbols are the same as in figure 19. Matches are represented as white squares, mismatches for the (+) sequence as bottom black wedges, mismatches for the (−) sequence as top black wedges, and double mismatches as black squares. They delimit *R*-blocks, consisting of mismatch-free intervals followed by a mismatch. The last nucleotide of an *R*-block is always a mismatch: the mismatch-free interval at the tail of the read is not an *R*-block. MEMs (arrows) are stretches of the read that match any of the two sequences and that cannot be extended left or right. MEMs matching the (+) sequence are represented at the top and MEMs matching the (−) sequence at the bottom. In this example, the central stretch of 5 nucleotides matching the (+) sequence cannot be used as seed because it is included in a stretch of 12 nucleotides matching the (−) sequence.

#### Remark 21.

We consider that when a MEM matches both sequences (as the head and the tail of the read shown in figure 22 for instance), both sequences are in the list of candidates after seeding, i.e. both are aligned.

As in section 4.2, reads are seen as sequences of *R*-blocks. We will first focus on the probability that a read does not contain a γ-MEM seed for the (+) sequence, from which we will later deduce the probabilities of type I and type II errors. This restriction imposes relatively complex constraints on the *R*-blocks. For instance, an *R*-block of size γ *(i.e*. γ – 1 matches followed by a mismatch) must be followed by a mismatch if the final nucleotide is a mismatch for the (−) sequence, but it may be followed by up to γ – 1 matches if this nucleotide is a double mismatch, or by any number of matches if it is a mismatch for the (+) sequence.

To give a concrete example, we will consider γ-MEMs of size 3. The read below starts with two matches, followed by a double mismatch, followed by a mismatch for the (+) sequence. The symbols between brackets (*, 0) indicate the respective amount of nucleotides matching the (−) and the (+) sequences at the end of the *R*-block. All the *R*-blocks terminated by a mismatch against the (+) sequence are equivalent because the subsequent *R*-block cannot be part of a MEM for the (+) sequence. So the only information we need regarding the (−) sequence is whether the number of matching nucleotides at the end or the *R*-block is greater than 0, which we represent by the * symbol. At that point, there are infinitely many possibilities for the next *R*-blocks, but they can be gathered in four distinct classes that do not create a MEM seed for the (+) sequence.

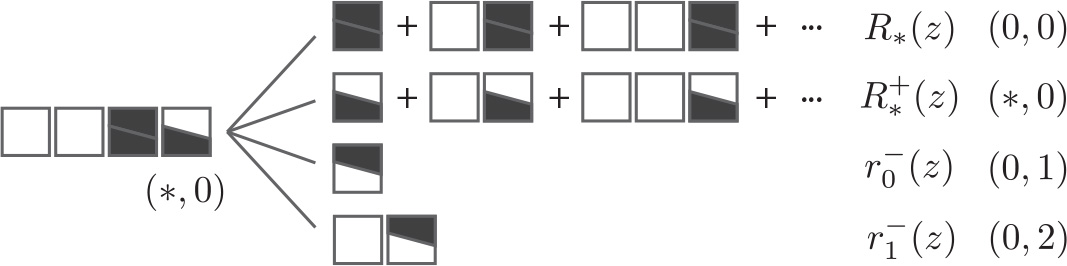

Appending any number of matches followed by a double mismatch (top case) brings the number of nucleotides matching (−) and (+) at the end of the *R*- block to (0,0). The weighted generating function of this union of *R*-blocks is denoted *R*_*_(*z*). Appending any number of matches followed by a mismatch against the (+) sequence (second case from the top) maintains the amount of matching nucleotides as (*, 0). The weighted generating function of this union of *R*-blocks is denoted 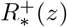. Appending a mismatch for the (−) sequence (third case from the top), *i.e*. an *R*-block with weighted generating function 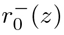 brings the amount of matching nucleotides to (0,1). Finally, appending a match followed by a mismatch for the (−) sequence (bottom case), *i.e*. an *R*-block with weighted generating function 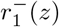 brings the amount of matching nucleotides to (0, 2). All other *R*-blocks create a MEM seed for the (+) sequence.

By considering all the possible scenarios for other reads, we obtain the transfer graph shown in figure 23. The numbers in the vertices indicate the amount of nucleotides matching (−) and (+) at the end of the *R*-blocks. Since *R*-blocks are terminated by a mismatch, at least one of these numbers is 0. As explained above, the * symbol stands for one or more nucleotides, as all these cases are equivalent. The body of the transfer graph is represented on the left panel, the head and tail edges on the right panel.

Simple *R*-blocks have weighted generating functions 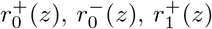 or 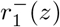. The sign indicates which sequence is *mismatched* at the end of the *R*- block and the index indicates the number of matches at the start of the *R*-block. Unions of *R*-blocks have weighted generating functions *R*_*_(*z*), 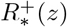 or 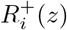. The first represents any number of matches followed by a double mismatch, the second represents any number of matches followed by a mismatch against the (+) sequence, the last represent *up to i* matches followed by a mismatch against the (+) sequence.

In the initial state of the read, the amount of matching nucleotides is (0, 0), which is indicated by the label 1 on the edge between the head vertex and the vertex (0,0). This is equivalent to prepending the read by the empty object *ε*. Reads are terminated by a sequence of matches. The weighted generating functions *F*_*_(*z*), *F*_0_(*z*), *F*_1_(*z*) and *F*_2_(*z*) represent mismatch-free intervals of any size or size *up to* 0, 1 or 2, respectively. Here, the empty sequence is present in the graph (take a path from the head vertex to the tail vertex through (0, 0) with no match appended at the tail), and no extra sequence needs to be added so *ψ*(*z*) = 0.

**Figure 23:**
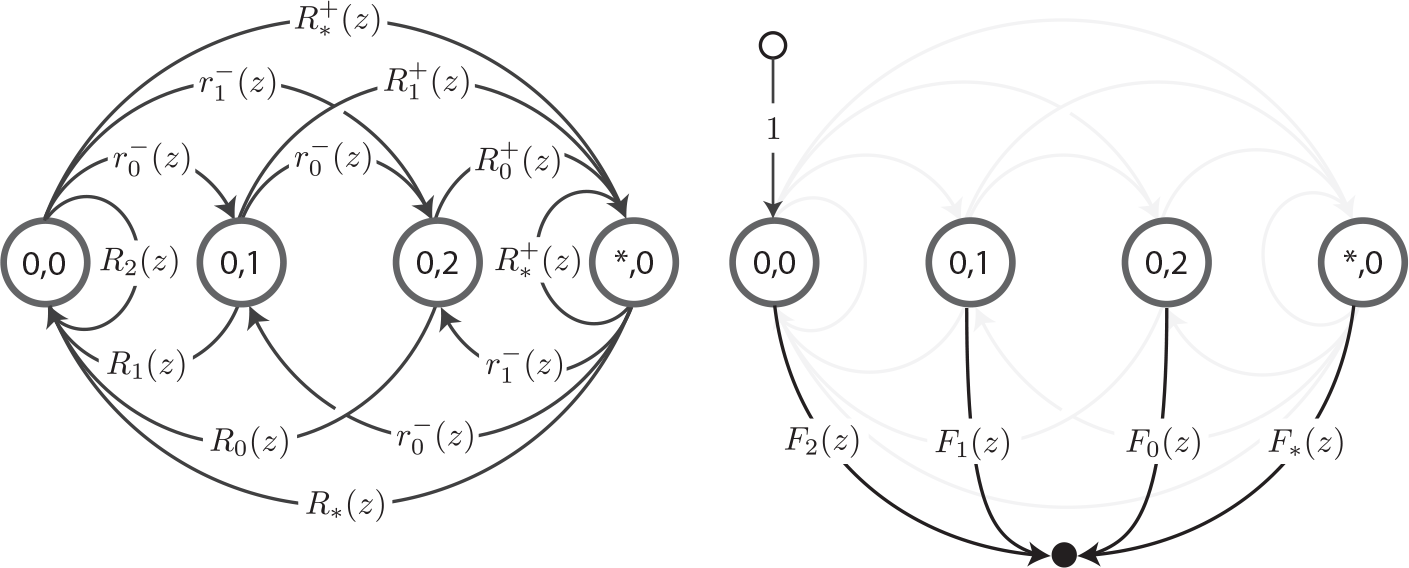
Transfer graph of reads without 3-MEM seed for (+). Reads are viewed as sequences of *R*-blocks, and they can match two sequences referred to as (+) and (−). To not overload the figure, the body of the graph is shown on the left panel, the head and tail edges are showed on the right panel. The numbers in the vertices represent the respective number of nucleotides matching the (−) and (+) sequences at the end of an *R*-block (the * symbol stands for one or more nucleotides). Weighted generating functions 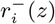 represent *R*-blocks with *i* matches terminated by a mismatch against the (+) sequence. Weighted generating functions *R*_*i*_(*z*) and 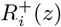 represent *R*-blocks with *up to i* matches terminated by a double mismatch or by a mismatch against (+), respectively. Weighted generating functions *F*_*i*_(*z*) represent mismatch-free intervals of size *up to i*. In the variants *R*_*_(*z*), 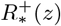 and *F*_*_(*z*), there is no upper limit on the number of matching nucleotides.

The same logic can be applied for higher values of γ. The general transfer matrix associated with the body of the transfer graph has dimension γ +1 and is defined as

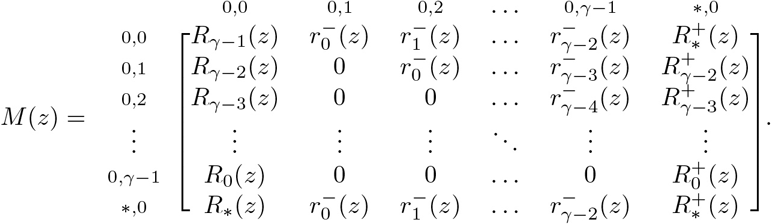

The head vector *H*(*z*) is equal to (1, 0,…, 0)^⊤^ and the tail vector *T*(*z*) is equal to (*F*_γ−1_(*z*),…, *F*_1_(*z*), *F*_0_(*z*), *F*\*(*z*))^⊤^. The weighted generating function of reads with no γ-MEM for the (+) sequence is omitted here. As per proposition 4, it can be computed as by *S*+(*z*) = *H*(*z*)^⊤^ · (*I* – *M*(*z*))^−1^ · *T*(*z*).

As in section 4.2, denote *a* the probability that the nucleotide is a double match, *b* that it is a mismatch against (+), *c* that it is a mismatch against (−) and d that it is a mismatch for both. With these definitions, expressions (27) still apply, and we also have

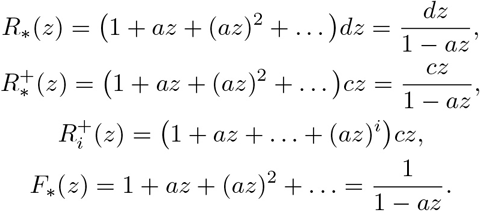

Knowing the values of *a*, *b*, *c* and *d*, we could compute the weighted generating function and use proposition 5 to obtain an asymptotic estimate of the probability that a read does not contain a γ-MEM for the (+) sequence. If we consider that the (+) sequence is the target and that the (−) sequence is the duplicate, this is the probability that we set out to find, i.e the probability that a read contains a γ-MEM seed when the target sequence has exactly one duplicate. The model is completely specified by the substitution rates *p* and *κ*, and as in section 4.2 *a* = (1 – *p*)(1 – *κ*), *b* = *pκ*/3, *c* = (1 – *P*)*k* and *d* = *p*(1 – *κ*/3). The example below shows how the computations are carried out in practice.

#### Example 22.

Let us approximate the probability that a read does not contain a MEM seed (where the target has one duplicate) for a read of size *k* = 100 with γ = 17 and for *p* = 0.10 and *κ* = 0.10. First, we substitute in the 18 × 18 transfer matrix *M*(*z*) and in *T*(*z*) the values *a* = 0.81,6 = 0.09, *c* = 0.00333, *d* = 0.09667. We then compute the expression 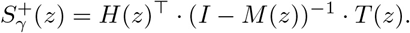 and obtain a rational function *P*(*z*)/*Q*(*z*) where *P* and *Q* are polynomials of degree 35 and 36, respectively. The root of *Q* with smallest *modulus* is the dominant singularity z_1_ of the weighted generating function. Using numerical approaches, we find *z*_1_ ≈ 1.0266331946. We compute the proportionality constant of proposition 5 as —*P*(*z*_1_)/*Q*′(*z*_1_) ≈ 1.3893346 from which we obtain the final estimate 1.3893346/1.0266331946^101^ ≈ 0.09769252. For comparison, a 99% confidence interval obtained by performing 10 billion random simulations is 0.097682 0.097698.

Figure 24 illustrates the precision of this estimate for different values of the error rate *p* and of the read size *k*. The values of the parameters are chosen to match those of figure 7 where we used exact seeds. In the conditions above, the seeding probabilities are alomst indistinguishable. This does not mean that the cost of using MEM seeds is always negligible. When using MEMs, the seeding probability decreases as the number of duplicates increases (here we assume that there is only one duplicate). In addition we are not separating type I and type II errors, which typically have different costs.

This raises the question of how to compute the type I error rate when using MEM seeds. The work above provides all the elements we need to approximate it. Call the P_3_ the approximate probability that the read does not contain a γ-MEM seed. Using the results from section 4.2, we can compute an asymptotic estimate of the probability that a read does not contain an exact γ-seed *and* does not contain any stretch of size γ or greater matching the duplicate sequence. We previously called this estimated probability *P*_2_. We can estimate the probability of type I errors as *P*_3_ – *P*_2_.

**Figure 24:**
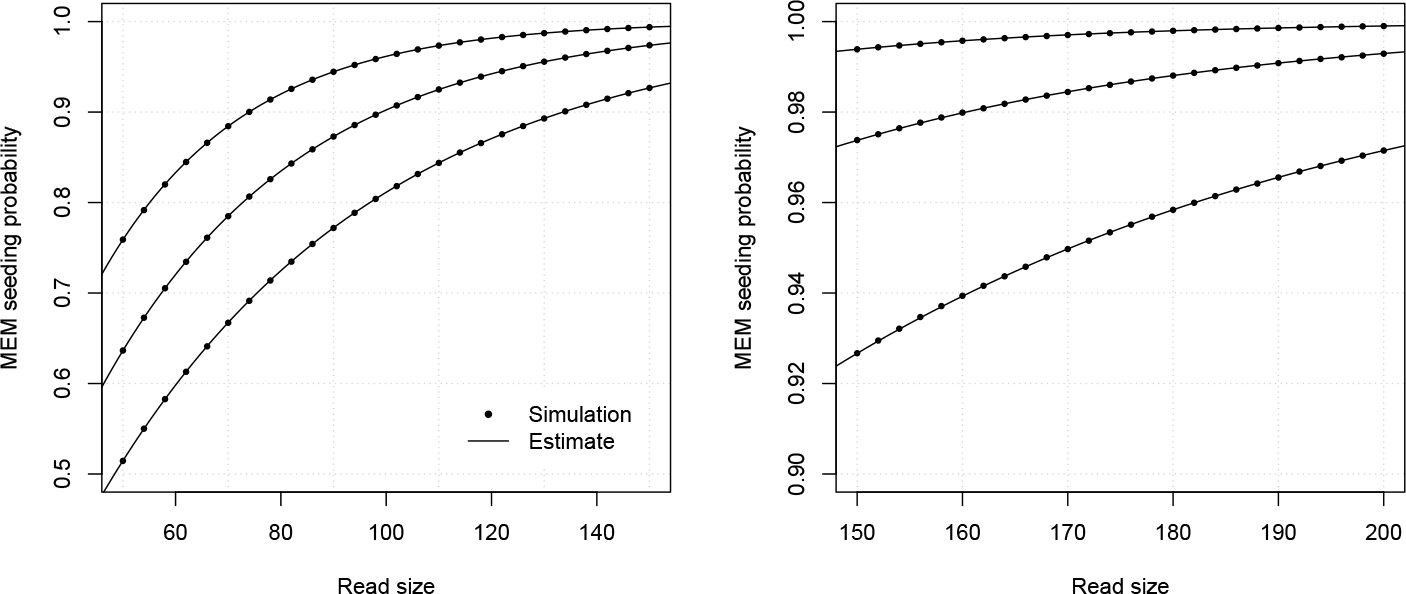
Example estimates of MEM seeding probability (one duplicate). The analytic combinatorics estimates are benchmarked against random simulations. Shown on both panels are the probablities that a read of given size will contain a MEM seed, either estimated by 10,000,000 random simulations (dots), or by the method described above (lines). The curves are drawn for γ = 17 and *κ* = 0.10, *p* = 0.08, *p* = 0.10 or *p* = 0.12 (from top to bottom). Notice how close the values are to those shown in figure 7 (probabilities that the read contains an exact seed with the same parameter values).

Let us clarify the last statement. Call *C* the event that the read has no γ-MEM seed and B the event that it does not contain an exact γ-seed *and* does not contain any stretch of size γ or greater matching the duplicate sequence. First observe that *B* ⊂ *C* so *P*(*C* ⋂ *B*^c^) = *P*(*C*) – *P*(*B*). Then, *B*^c^ is the event that the read contains an exact γ-seed, a stretch of size γ or greater matching the duplicate sequence, or both. Another way to describe *B*^c^ is that “some hit is found in the read”. So *C* ⋂ *B*^c^ is the event that “some hit is found, but it is not the target”, which is the definition of a type I error. Since *P*(*C*) ≈ *P*_3_ and *P*(*B*) ≈ *P*_2_, the probability of type I errors is indeed approximately equal to *P*_3_ – *P*_2_.

#### Remark 22.

The rationale above shows that the probability of type II error with MEM seeds is approximately equal to the probability *P*_2_ computed at section 4.2.

The example below shows how the probabilities of type I errors when using MEM seeds are computed in practice.

#### Example 23.

Let us approximate the probability of type I error with MEM seeds (with one duplicate) for a read of size *k* = 100 with γ = 17 and for substitution rates *p* = 0.10 and *κ* = 0.10. In example 20 we found *P*_2_ ≈ 1.4032791/1.0272930245^101^ and in example 22 we found *P*_3_ ≈ 1.3893346/1.0266331946^101^. The type I error rate is then approximately equal to P_3_ – P_2_ ≈ 0.00521934. For comparison, a 99% confidence interval obtained by performing 10 billion random simulations is 0.0052166 – 0.0052203.

Figure 25 illustrates the precision of the estimates for different values of the substitution rate *κ* and of the read size *k*. The approximation is grossly inaccurate for *κ* = 0.05, where it is even negative for reads under 40 nucleotides. The issue here is that the approximate coefficients of *S*+(*z*) *converge too slow*. To illustrate the point, for *k* = 40 and *κ* = 0.05, the analytic combinatorics estimate *P*_3_ is approximately 3% too low, but this error is already 5 times the target type I error rate.

**Figure 25:**
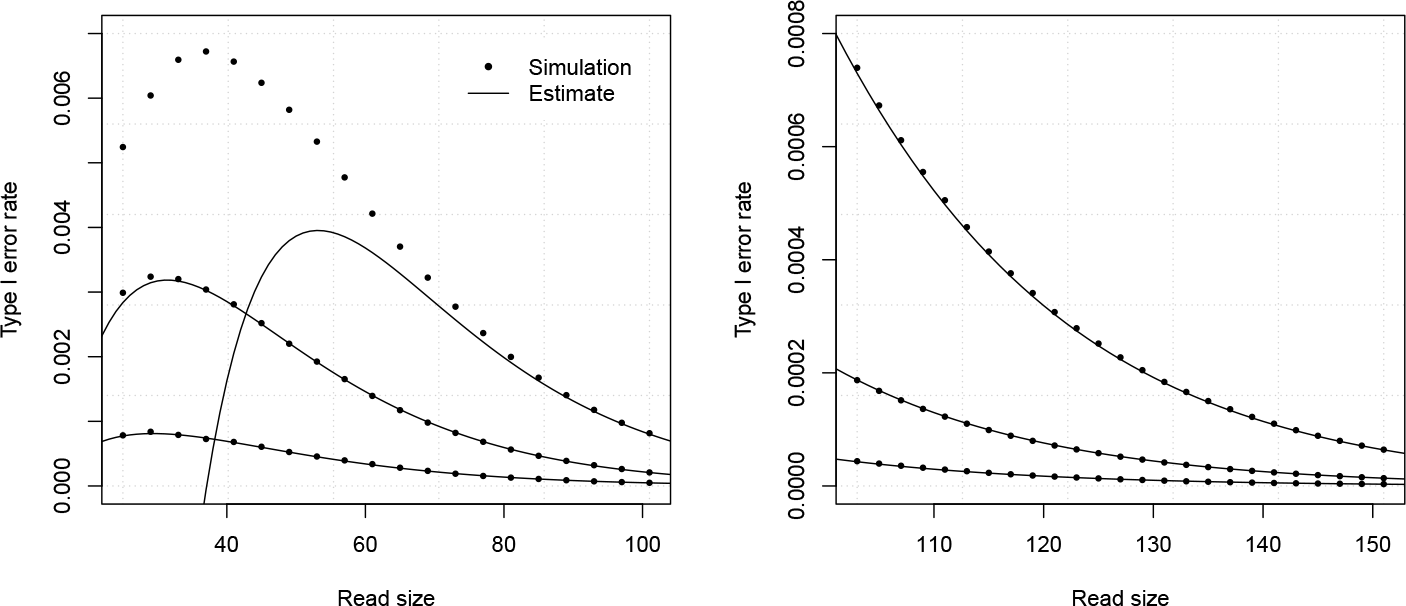
Example estimates of MEM seeding probability (one duplicate). The analytic combinatorics estimates are benchmarked against random simulations. Shown on both panels are the probablities that a read of given size will yield a type I error when using MEM seeds, either estimated by 10,000,000 (left) or 100,000,000 (right) random simulations (dots), or by approximating the coefficients of 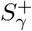 (lines). The curves are drawn for γ = 17 and *p* = 0.05, *κ* = 0.05, *κ* = 0.15 or *κ* = 0.25 (from top to bottom).

Recall that the proof of proposition 5 shows how we can obtain the exact value of 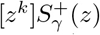 by making use of all the singularities of the weighted generating function (see remark 9). For a rational function *W*(*z*) = *P*(*z*)/*Q*(*z*),

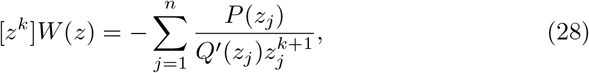

where *z*_1_, *z*_2_,…, *z*” are the singularities of *W* sorted by increasing order of *modulus*.

It seems that we could improve the estimate by using more than just the dominant singularity But here we are on slippery ground because using more singularities can also make the estimate *less accurate*. This may come as a surprise, since the proof of proposition 5 suggests that each singularity improves the approximation. However, the fact that the asymptotic estimate converges faster does not mean that it is more accurate for small values of *k*.

Figure 26 illustrates this very vividly. It represents the same data as the left panel of figure 25, except that the analytic combinatorics estimates are computed with the first three terms of (28) instead of just the first. For *κ* = 0.15 and *κ* = 0.25 (bottom two curves), the approximations were reasonable with only one singularity, but they became disastrous with three. In contrast, for *κ* = 0.05 (top curve) adding two singularities made the estimates much more accurate.

**Figure 26:**
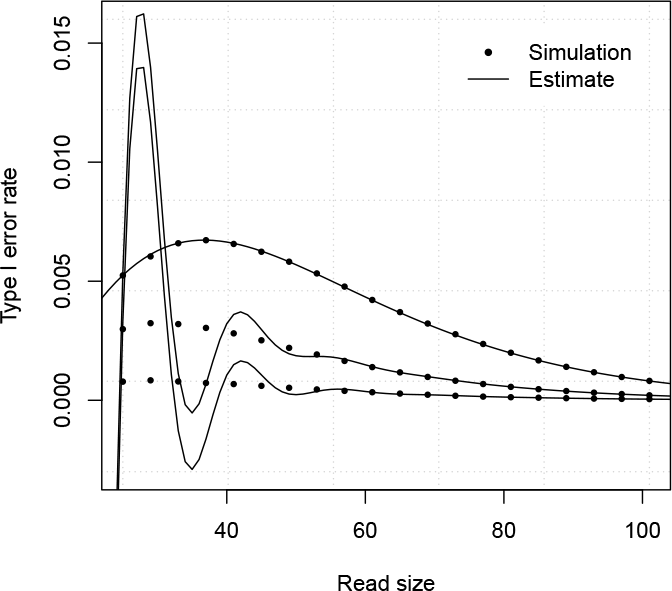
Higher order asymptotic approximations. The data is the same as that represented in the left panel of figure 25, except that the analytic combinatorics estimtates (lines) are computed from three singularities instead of one, *i.e*. using the first three terms of (28).

Looking in more detail, we observe that the estimates are improved when the singularities of 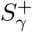 have a relatively small imaginary part, and they are degraded when the singularities have a large imaginary part. Fortunately, these cases are very easily distinguished because the second singularities orbit around only two clusters, as shown in figure 27. The trend persits for all the tested values of γ up to 30; *z*_2_ is either in the real spectrum or it is a complex number with a large imaginary part.

For some values of *p* and *κ*, the coefficients of 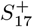 have a two-phase decay, whereas for others they have a simple exponential decay. Why this is the case is unclear, but two-phase decays are better approximated using more than one singularity. If the decay is a simple exponential, the second singularity captures some oscillations of the coefficients that are irrelevant for the problem at hand and that are detrimental to the approximation.

**Figure 27:**
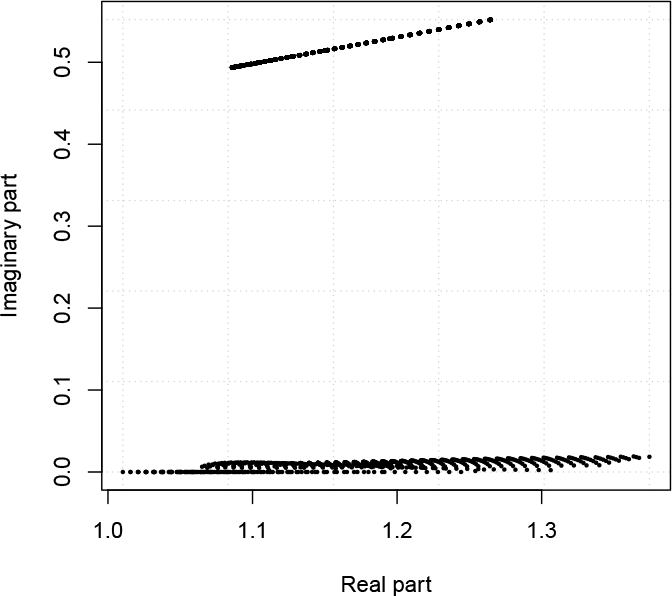
Non-dominant singularities of 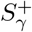. The dots represent the second singularity of 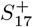 plotted in the complex plane for 2,500 values of *p* and *κ* between 0.005 and 0.25. The imaginary part of *z*_2_ is either close to 0 or close to 0.5.

In any event, this distinction between real (or near real) versus complex *z*_2_ gives a practical method to approximate the coefficients of S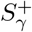 with higher precision and from there estimate the type I error rate associated with MEM seeds. The overall performance of this approach is illustrated in figure 28.

Extending the estimates to more than one duplicate is more challenging that in section 4.2 because conditional independence does not hold for MEM seeds. But we can still use the estimates above as a lower bound for the probability of type I error. If the proportion of the genome that is repeated is *x* and *P*\* is the maximum of *P*_3_ – *P*_2_ relative to unknown *κ*, then *xP*\* is again a reasonable lower bound on the probability of type I errors.

As in section 4.2, the methods presented here suffer from the difficulty to know the real number of duplicates of the target. Progress on parallel research lines will be needed in order to develop a workable theory of the relationships between repeated sequences of a genome.

## 5 Average quantities

### 5.1 General approach

The analytic combinatorics approach allows computing the average of many quantities of interest. In the seeding problem, one such quantity is the number of errors for reads of different kinds. So far, weighted generating functions only marked the size of the reads through the variable *z*. We now need to introduce another variable to mark the second quantity, which means that we will deal with bivariate weighted generating functions. For concreteness, if *a*_*k*_,_*n*_ is the total weight of reads of size *k* with n errors and without exact seed, the average number of errors for such reads is

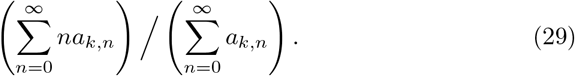

**Figure 28:**
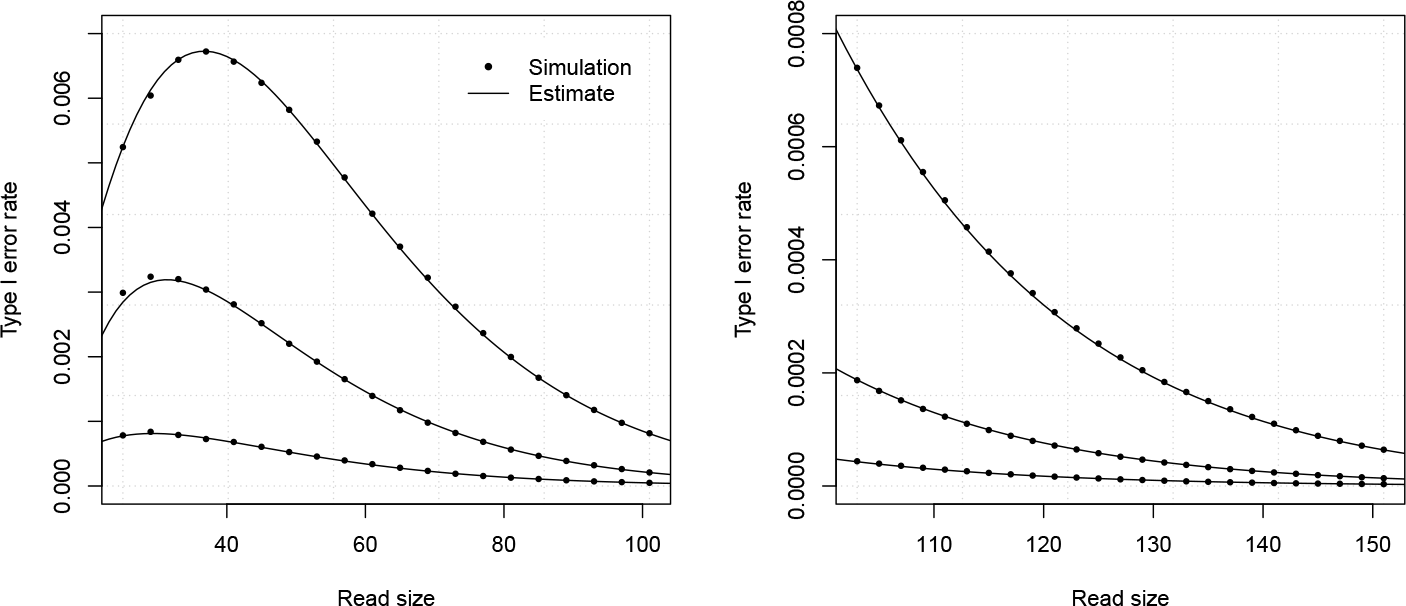
Example multi-singularity estimates of type I error rates. The analytic combinatorics estimates with either one or three singularities (see text) are benchmarked against random simulations. Shown on both panels are the probablities that a read of given size will give a type I error when using MEM seeds, either estimated by 10,000,000 (left) or 100,000,000 (right) random simulations (dots), or by the method described above (lines). The curves are drawn for γ = 17 and p = 0.05, *κ* = 0.05, *κ* = 0.15 or *κ* = 0.25 (from top to bottom). The simulation data points are the same as those shown in figure 25. Three singularities are used for *κ* = 0.05, and one for *κ* = 0.15 and *κ* = 0.25.

To compute this quantity, we introduce the variable *u* marking the number of substitutions and we write

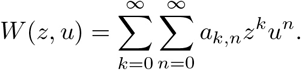

Finding an explicit formula for *W*(*z*, *u*) is the focus of sections 5.2 to 5.4. For now, observe that (29) can be expressed from *W*(*z*, *u*) as follows. On the one hand, we have

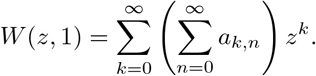

So the denominator of (29) is the coefficient of *z*^*k*^ in *W*(*z*, 1). On the other hand, by taking the derivative of *W*(*z*, *u*) with respect to *u*, and setting *u* = 1, we obtain

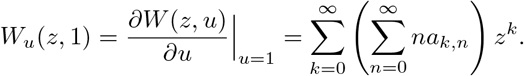

So the numerator of (29) is the coefficient of *z*^*k*^ in *W*_*u*_(*z*, 1). Note that *W*(*z*, 1) and *W*_*u*_(*z*, 1) are actually univariate weighted generating functions, so we can use the methods developed earlier to obtain asymptotic estimates for their coefficients. However, proposition 5 will not apply to approximate the coefficients of *W*_*u*_(*z*, 1). Since all the weighted generating functions considered here are ratios of polynomials of the form *W*(*z*, *u*) = *P*(*z*, *u*)/*Q*(*z*, *u*), the derivative *W*_*u*_(*z*, 1) can be expressed as (*P*_*u*_(*z*, 1)*Q*(*z*, 1) – *P*(*z*, 1)Q_*u*_(*z*, 1))/*Q*(*z*, 1)^2^. Because of the term *Q*(*z*, 1)^2^ at the denominator, the singularities of *W*_*u*_(*z*, 1) are not simple poles.

We need to work out new asymptotic estimates. Proposition 10 below shows how to extract the coefficients of weighted generating functions in this particular case.

#### Proposition 10.

If a function *W*(*z*) is the ratio of two polynomials *P*(*z*)/*Q*(*z*)^2^, and *Q* has only simple roots, then

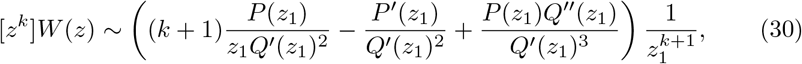

where *z*_1_ is the root of *Q* with smallest modulus.

As in proposition 5, we start by proving the lemma that will give the functional expression of the asymptotic expansion.

#### Lemma 3.

For |*z*| < a we have

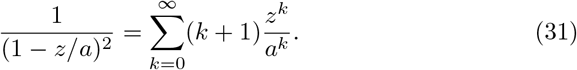

*Proof*.

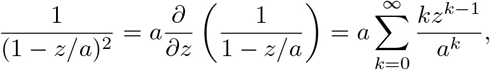

where the last equality is obtained by applying lemma 2 and differentiating with respect to z.

We now prove proposition 10.

*Proof*. As in the proof of proposition 5, let *z*_1_, *z*_2_,…, *z*_*n*_ be the complex roots of *Q*, sorted by increasing order of *modulus*. Since the roots of *Q*(*z*)^2^ all have multiplicity 2, there exists constants *α*_1_,…, *α*_*n*_ and *β*_1_,…, *β*_*n*_ such that the partial fraction decomposition of the rational function *P*(*z*)/*Q*(*z*)^2^ can be written as

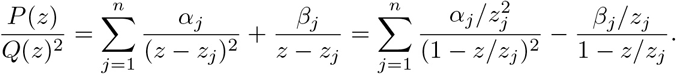

As in the proof of proposition 5, we assumed without loss of generality that the degree of *P* is lower than the degree of *Q*^2^. Now applying lemmas 2 and 3, we see that the coefficient of *z*^*k*^ in *P*(*z*)/*Q*(*z*)^2^ can be expressed as

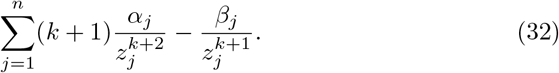

The sum above is dominated by the term with the highest exponential rate of increase, *i.e. j* = 1 because zi has smallest *modulus* by definition. Thus, the coefficient of *z^k^* in *P*(*z*)*/Q*(*z)^2^* is asymptotically equivalent to

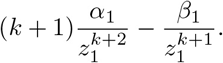

We now need to find the values of *α*_1_ and *β*_1_. As *z*_1_ is a root of *Q*, there exists a polynomial *Q*_1_(*z*) such that

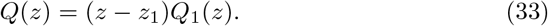

We can thus write *P*(*z*)/*Q*(*z*)^2^ as

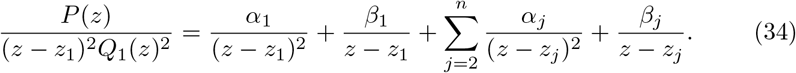

Multiplying both sides of (34) by (*z* – *z*_1_)^2^ and setting *z* = *z*_1_ shows that *α*_1_ = *P*(*z*_1_)/*Q*_1_(*z*_1_)^2^. To find the value of *Q*_1_(*z*_1_) we differentiate (33) and let *z* = *z*_1_, yielding *Q*′(*z*_1_) = *Q*_1_(*z*_1_). We thus obtain *α*_1_ = *P*(*z*_1_)/*Q*′(*z*_1_)^2^.

To find the value of *β*_1_, we subtract *α*_1_/(*z* – *z*_1_)^2^ on both sides of (34) and obtain

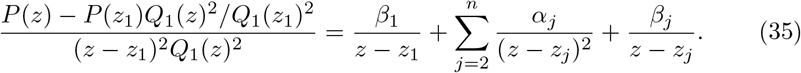

Since *P*(*z*_1_) – *P*(*z*_1_)*Q*_1_(*z*_1_)^2^/*Q*_1_(*z*_1_)^2^ = 0, there exists a polynomial *Q*_2_(*z*) such that

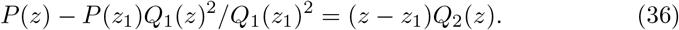

Multiplying (35) by *z-z*_1_ and setting *z* = *z*_1_, we obtain = *Q*_2_(*z*_1_)/*Q*_1_(*z*_1_)^2^. To find the value of *Q*_2_(*z*_1_) we differentiate (33) two times and (36) one time, finally obtaining *β*_1_ = *P*′(*z*_1_)/*Q*′(*z*_1_)^2^ – *P*(*z*_1_)*Q*“(*z*_1_)/*Q*^/^(*z*_1_)^3^, which concludes the proof.

#### Remark 23.

Expression (30) is asymptotically equivalent to the simpler expression

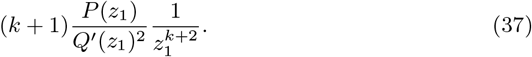

However, the convergence of expression (37) is slow. Indeed, dividing the exact expression (32) by (37) gives an error term in *O*(1/*k*). In comparison, dividing (32) by (30) gives an error term in *O*(|*z*_1_/*z*_2_|^*k*^), which decreases exponentially, as in proposition 5.

We are now ready to approximate the mean number of errors in reads of different kinds, treating the distinct types of errors separately.

### 5.2 Substitutions

We fist find the average number of substitutions in the simple error model of section 3.3. Applying a now familiar strategy, we write the generating function of simple objects that we combine into more complex objects. Recall that in the uniform substitution model, the weighted generating function of substitutions is *pz*, where *p* is the error rate. Marking substitutions with the variable u is as simple as replacing their weighted generating function by *pzu*. For every substitution in the read, the power of *u* increases by 1, while the power of *z* still increases by 1 for every nucleotide.

We could derive the weighted generating function by the transfer matrix method described in section 3.3, but we can get an immediate result by using equation (12), where we observed that the weighted generating function of reads without exact seed can be expressed as

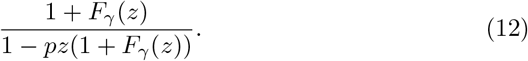

Here the weighted generating function of substitutions appears explicitly as *pz*. We can replace it by *pzu* to obtain directly

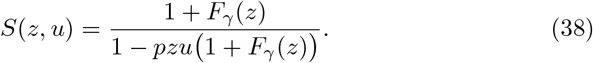

*S*(*z*, 1) is simply the weighted generating function of reads without seed derived in section 3.3, and for which we already know the coefficient asymptotics. So we already have the denominator of (29). Now differentiating (38) with respect to *u*, we obtain

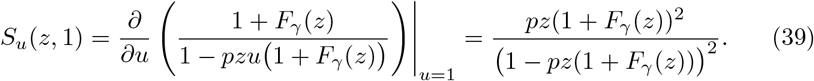

As mentioned above, the form of the weighted generating function calls for using proposition 10 (and not proposition 5). The dominant singularity is the same for both the numerator and the denominator of (29), so the terms in 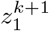 cancel out. What remains is an expression of the form *C*_1_*k* + *C*_2_, where *C*_1_ and *C*_2_ are constants. In other words, the average number of substitutions in reads without exact seed increases as an affine function of the size. We make this result more accurate in the following proposition.

#### Proposition 11.

Under the assumptions of the error model of section 3.3, the average number of substitutions in reads without seed is asymptotically equivalent to

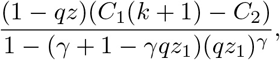

where *k* is the size of the read, where

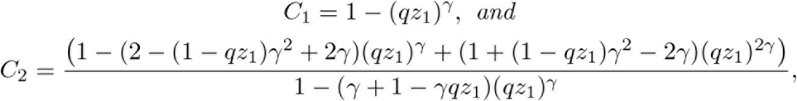

and where *z*_1_ is the only real positive root of the polynomial 1 – *pz*(1 + *F*_γ_(*z*)).

*Proof*. Apply proposition 10 to (39), use proposition 7 and simplify.

**Figure 29:**
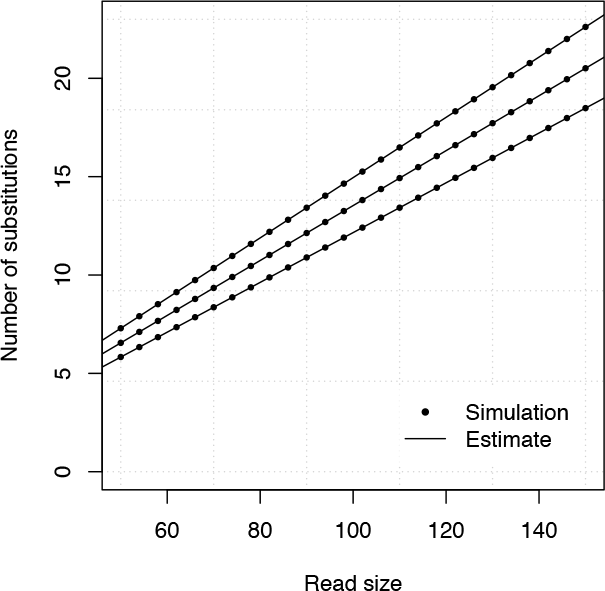
Estimating the average number of substitutions. The average number of substitutions in reads without γ-exact seed is shown for different read sizes, either estimated by 10,000,000 random simulations (dots), or by analytic combinatorics (lines). The curves are drawn for γ = 17, *p* = 0.08, *p* = 0.10 or *p* = 0.12 (from bottom to top).

Figure 29 illustrates the accuracy of the estimate. The affine relationship between the average number of errors and the size of the read is apparent. Also, the approximations are very close to the target values.

#### Remark 24.

What about the average number of substitutions in reads that contain a γ-exact seed? We can compute it indirectly from the quantities that we already have defined,.

The average number of substitutions for reads of size *k* (regardless of whether they contain a seed) is *p_1_k*. Call *P*_1_,*_k_* the probability that a read of size *k* contains no seed and *E_k_* the average number of substitutions in such reads. The quantity of interest, × follows the relationship

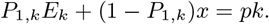

From this we see that the average number of substitutions in reads that contain a γ-exact seed is (*pk – P_1_,_k_E_k_*)/(1 – *P*_1_,*_k_*). This relationship is not affine because *P*_1_,*_k_* decreases exponentially as k increases. But for long reads *P*_1_,*_k_* vanishes so the average is approximately linear and equal to *p_1_k*.

### 5.3 Deletions

To count the average number of deletions, we could take the weighted generating function of the reads without seed in the error model of section 3.4, namely

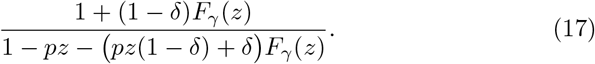

But here it is important to remember that, equation (17) ignores the deletions that occur immediately before and after substitutions, because they have no impact on the presence of a seed. For the purpose of counting deletions, we need to take them *all* into consideration, even those that do not interrupt any error-free interval. This requires working out the proper weighted generating function.

A read can be thought of as a walk on the graph shown in figure 30. On this graph, deletions appear as a separate vertex (unlike in figure 8). Their size is 0 and they occur only between nucleotides, which implies that a read can neither start nor end with a deletion. Also, a deletion must be followed by either a match or a substitution; it cannot be followed by a deletion. The reason is that a deletion has size 0 regardless the number of deleted nucleotides. In the symbolic representation of the read, either there is a deletion of any size between two nucleotides, or there is no deletion.

Here, reads are sequences of error-free intervals withs symbol Δ_0_ and weighted generating function *F*(z) or substitutions with symbol S and weighted generating function *pz*, or deletions with symbol *D* and weighted generating function *δ*. Two error-free intervals cannot follow each other (together they form a single error-free interval) and two deletions cannot follow each other (together they form a single deletion). Here we also need to account for the absence of deletions in transitions between error-free intervals and substitutions. This is done by weighing the corresponding transitions by a factor (1 – *δ*).

**Figure 30:**
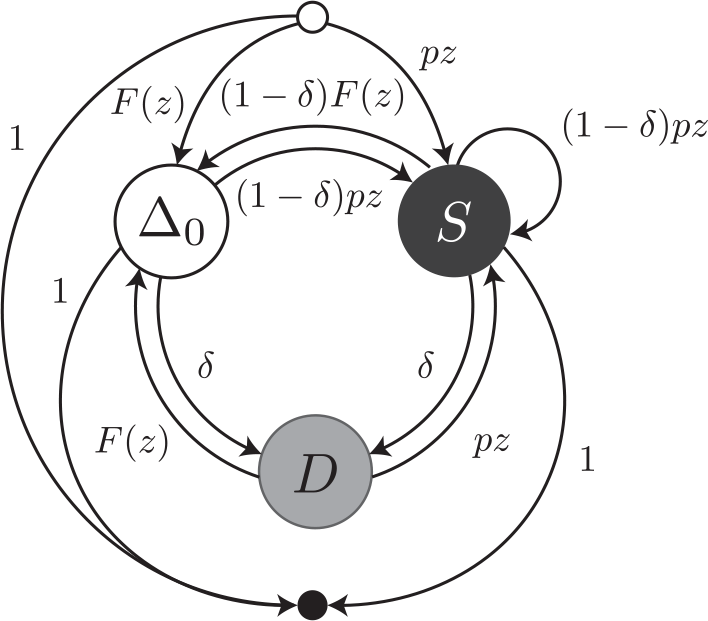
Transfer graph with explicit deletions. Reads are viewed as sequences of error-free intervals (symbol Δ_0_) or substitutions (symbol *S*) or deletions (symbol *D*), with respective weighted generating functions *F*(*z*), *pz* and *δ*.

As in the previous section, in order to count deletions with parameter *v*, we replace their weigted generating function by *δv*. For every deletion the power of *v* increases by 1 (and the power of *z* does not change). The transfer matrix then becomes a function of *z*, marking the size of the reads, and of *v*, marking the number of deletions. The final expression of the transfer matrix is

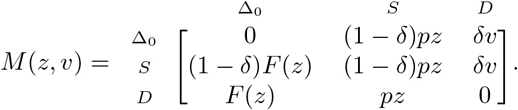

Recall that reads can neither start nor end by a deletion because they are undetectable at those positions. Thus the head vector is *H*(*z*, *v*) = (*F*(*z*), *pz*, 0)^⊤^ and the tail vector is *T*(*z*, *v*) = (1,1, 0)^⊤^. The only sequence we need to add is the empty sequence *ε*, so *ψ*(*z*, *v*) = 1. Following proposition 4, the weighted generating function *S*(*z*, *v*) is equal to *ψ*(*z*, *v*)+*H*(*z*, *v*)^⊤^ ·(*I*— *M*(*z*, *v*))^−1^ ·*T*(*z*, *v*).

Following the general strategy to count average quantities, we first compute *S*(*z*, 1). Naturally, we obtain expression (17), which is the weighted generating function of reads without exact seed. We also differentiate *S*(*z*, *v*) with respect to v and let *v* = 1 to obtain

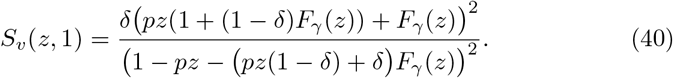

As in the section 5.2, we derive the asymptotics by applying proposition 10 to (40) and by applying proposition 5 to (17). The terms 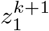 cancel out when computing the ratio as per equation (29), so the number of deletions in reads without seed is asymptotically affine, *i.e*. of the form *C*_1_*k* + *C*_2_, where *C*_1_ and *C*_2_ are constants. In both cases, C_1_ and *C*_2_ have cumbersome expressions, but they are easy to compute from propositions 5 and 10.

**Figure 31:**
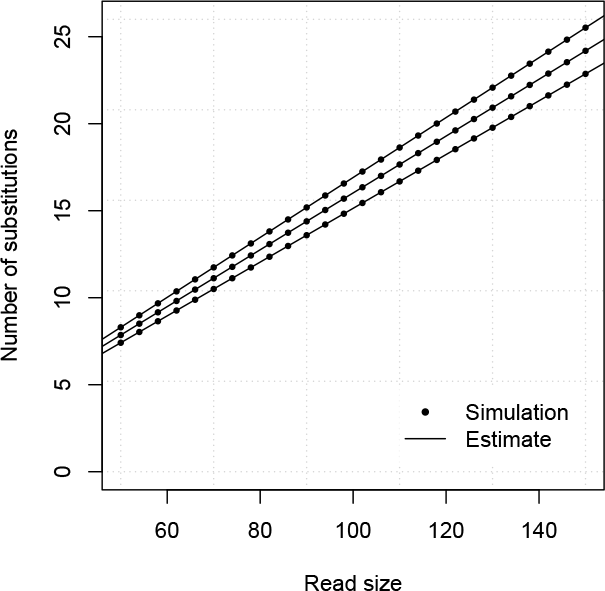
Example estimates of average number of deletions. The average number of deletions in reads without γ-exact seed is shown for different read sizes, either estimated by 10,000,000 random simulations (dots), or by analytic combinatorics (lines). The curves are drawn for γ = 17, *p* = 0.05 and *δ* = 0.14, *δ* = 0.15 or *δ* = 0.16 (from bottom to top).

The accuracy of the estimates is illustrated in figure 31. Once again the affine relationship is a good fit and the approximations are close the target values.

### 5.4 Insertions

Finally, we compute the average amount of insertions for reads without exact seed in the full error model of section 3.5. The simplest way to do this is to mark insertions in the transfer matrix where they appear explicitly. For this, we append the extra variable w to their weighted generating functions and obtain the following transfer matrix

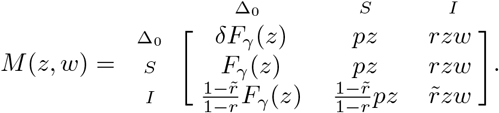

In the expression above, we have used *F*_γ_(*z*) instead of *F*(*z*) because we are directly considering reads without seed. We also update the head and tail vectors *H*(*z*, *w*) = (*F*_γ_(*z*), *pz*, *rzw*)^⊤^ and *T*(*z*, *w*) = (1,1,1)^⊤^, and we add the empty sequence so *ψ*(*z*, *w*) = 1. Following proposition 4, *S*(*z*, *w*) = *ψ*(*z*, *w*) + *H*(*z*, *w*)^⊤^ · (*I* – *M*(*z*, *w*))^−1^ · *T*(*z*, *w*).

Once again, we obtain the average number of insertions by computing ratio (29) from the coefficients of *S*(*z*, 1) and of those of *S*_*w*_(*z*, 1). Computing *S*(*z*, 1) yields expression (22), *i.e*. the weighted generating function of reads without exact seed. By differentiating *S*(*z,w*) and setting *w* = 1 we find *S_w_ (z*, 1) to be equal to

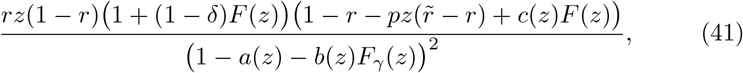

where *a*(*z*) and *b*(*z*) are defined as in (20), and where *c*(*z*) is a first degree polynomial defined as

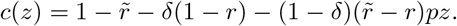

As in section 5.2, we derive the asymptotics by applying proposition 10 to (41) and by applying proposition 5 to (22). The terms 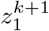 cancel out when computing the ratio as per equation (29), so the number of insertions in reads without seed is asymptotically affine, *i.e*. of the form *C*_1_*k* + *C*_2_, where *C*_1_ and *C*_2_ are constants. In both cases, *C*_1_ and *C*_2_ have cumbersome expressions, but they are easy to compute from propositions 5 and 10.

**Figure 32:**
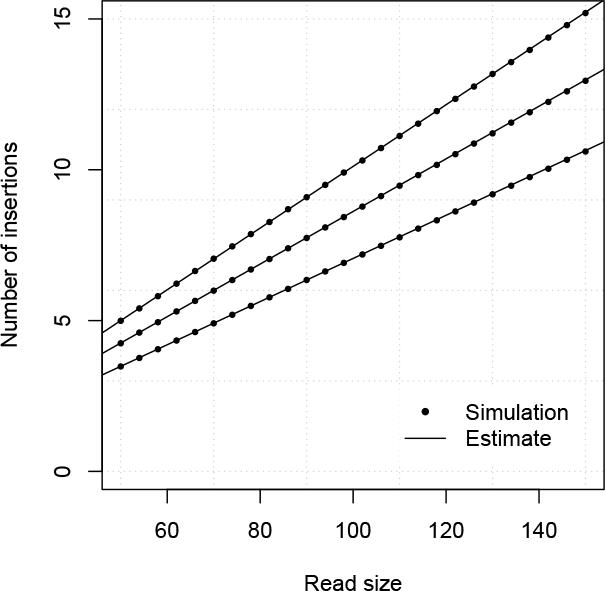
Example estimates of average number of isnertions. The average number of insertions in reads without γ-exact seed is shown for different read sizes, either estimated by 10,000,000 random simulations (dots), or by analytic combinatorics (lines). The curves are drawn for γ = 17, *p* = 0.05, *δ* = 0.15, *r̃* = 0.45 and *r* = 0.04, *r* = 0.05 or *r* = 0.06 (from bottom to top).

The accuracy of the estimates is illustrated in figure 32. Once again the affine relationship is a good fit and the approximations are close the target values.

It is also meaningful to compute the average number of substitutions or deletions in the full error model. For this, one can start from the transfer matrix, mark the quantity of interest and proceed with the general strategy exposed in section 5.1.

To compute the average number of substitutions, we use the transfer matrix

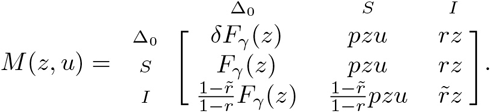

To compute the average number of deletions, we use the transfer matrix

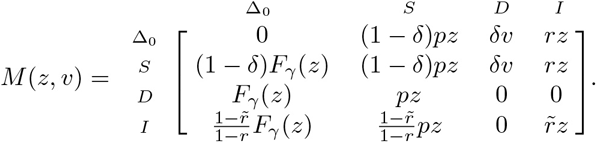

Finally, we can also consider all errors together (deletions always counting as a single error). For this we mark all the types of errors with the same variable, say w. In this case, we use the tansfer matrix

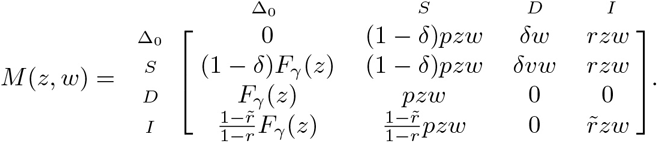

In all the cases, we obtain an affine relationship between the size of the read and the average number of error of a given type.

## 6 Conclusion

This concludes our introductory tour of the applications of analytic combinatorics to seeding methods. There is of course much more to say. Our purpose here is to show how the general strategy of analytic combinatorics gives reasonable solutions to problems that were previously difficult to address.

We have seen through multiple applications that the general strategy is to define combinatorial “atoms” with simple weighted generating functions (*e.g*. nucleotide symbols), combine these atoms into objects of increasing complexity (*e.g*. error-free intervals or reads without seed), construct their weighted generating functions from simple rules (mostly through proposition 4), and finally analyze the singularities of the weighted generating functions to approximate the quantities of interest (through propositions 5 and 10).

When using exact seeds, inexact seeds or MEM seeds, the estimates of the seeding probabilities are robust and relatively straightforward. The data collected during the alignment can be used to estimates the parameters of the error model to auto-tune the seeding heuristic during the run. This gives tight control over the seeding probability. More challenging is to control the type I error rate. The main difficulty is the lack of theoretical background on the seeding process when the target is duplicated. Significant progress on this line may not come from analytic combinatorics, but from new data structures or algorithms to seed such sequences more efficiently.

Seeding is not only used in mapping, but also in other alignment problems. In this regard, the work presented above can be applied to different contexts. That said, mapping high throughput sequencing reads is a “sweet spot” for analytic combinatorics because the sequences are usually long enough for the approximations to be accurate.

Finally, the concepts developed above can also be used to generate random reads of different kinds. The analogy between reads and walks on graphs with weighted edges allows us to generate reads by simulating a random walk. The trick is to convert the transfer matrix into a transition matrix by using the weights of the objects as their probabilities of occurrence.

In summary, analytic combinatorics is a powerful strategy that comes with a rich toolbox with many applications in modern bioinformatics. I expect more applications to see the light as new algorithms and new heuristics are developed in the future.

## Acknowledgements

I would like to thank Eduard Valera Zorita and Patrick Berger for their critical comments and their useful suggestions about the manuscript. I acknowledge the financial support of the Spanish Ministry of Economy and Competitiveness (Centro de Excelencia Severo Ochoa 2013-2017, Plan Nacional BFU2012-37168), of the CERCA Programme / Generalitat de Catalunya, and of the European Research Council (Synergy Grant 609989).

## References

[1] S F Altschul, W Gish, W Miller, E W Myers, and D J Lipman. Basic local alignment search tool. J. Mol. Biol., 215(3):403–10, October 1990.

[2] Richard Durbin, Sean R Eddy, Anders Krogh, and Graeme Mitchison,. Biological sequence analysis: probabilistic models of proteins and nucleic acids. Cambridge university press, 1998.

[3] P. Ferragina and G. Manzini. Opportunistic data structures with applications. In Proceedings of the 41st Annual Symposium on Foundations of Computer Science, FOCS ‘00, pages 390—, Washington, DC, USA, 2000. IEEE Computer Society.

[4] Philippe Flajolet and Andrew Odlyzko. Singularity analysis of generating functions. SIAM Journal on discrete mathematics, 3(2):216–240, 1990.

[5] Philippe Flajolet and Robert Sedgewick. An introduction to the analysis of algorithms, 1996.

[6] Philippe Flajolet and Robert Sedgewick. Analytic Combinatorics. Cambridge University Press, New York, NY, USA, 1 edition, 2009.

[7] Sara Goodwin, John D McPherson, and W Richard McCombie. Coming of age: ten years of next-generation sequencing technologies. Nat. Rev. Genet., 17(6):333–51, 05 2016.

[8] S Karlin and S F Altschul. Methods for assessing the statistical significance of molecular sequence features by using general scoring schemes. Proc. Natl. Acad. Sci. U.S.A., 87(6):2264–8, March 1990.

[9] S Karlin and S F Altschul. Applications and statistics for multiple high-scoring segments in molecular sequences. Proc. Natl. Acad. Sci. U.S.A., 90(12):5873–7, June 1993.

[10] Ben Langmead, Cole Trapnell, Mihai Pop, and Steven L Salzberg. Ultrafast and memory-efficient alignment of short DNA sequences to the human genome. Genome Biol., 10(3):R25, 2009.

[11] Heng Li and Richard Durbin. Fast and accurate short read alignment with Burrows-Wheeler transform. Bioinformatics, 25(14):1754–60, July 2009.

[12] Heng Li and Nils Homer. A survey of sequence alignment algorithms for next-generation sequencing. Brief. Bioinformatics, 11(5):473–83, September 2010.

[13] Kensuke Nakamura, Taku Oshima, Takuya Morimoto, Shun Ikeda, Hirofumi Yoshikawa, Yuh Shiwa, Shu Ishikawa, Margaret C Linak, Aki Hirai, Hiroki Takahashi, Md Altaf-Ul-Amin, Naotake Ogasawara, and Shigehiko Kanaya. Sequence-specific error profile of Illumina sequencers. Nucleic Acids Res., 39(13):e90, July 2011.

[14] Javier Quilez, Enrique Vidal, Francois Le Dily, Francois Serra, Yasmina Cuartero, Ralph Stadhouders, Thomas Graf, Marc A. Marti-Renom, Miguel Beato, and Guillaume Filion. Parallel sequencing lives, or what makes large sequencing projects successful. bioRxiv, 2017.

[15] Jason A Reuter, Damek V Spacek, and Michael P Snyder. High-throughput sequencing technologies. Mol. Cell, 58(4):586–97, May 2015.

[16] Yanni Sun and Jeremy Buhler. Choosing the best heuristic for seeded alignment of DNA sequences. BMC Bioinformatics, 7:133, March 2006.

